# A rigorous and versatile statistical test for correlations between time series

**DOI:** 10.1101/2022.01.25.477698

**Authors:** Alex E. Yuan, Wenying Shou

**Author notes:** (AY); (WS).

## Abstract

In disciplines from biology to climate science, a routine task is to compute a correlation between a pair of time series, and determine whether the correlation is statistically significant (i.e. unlikely under the null hypothesis that the time series are independent). This problem is challenging because time series typically exhibit autocorrelation, which cannot be properly analyzed with the standard iid-oriented statistical tests. Although there are well-known parametric tests for time series, these are designed for linear correlation statistics and thus not suitable for the increasingly popular nonlinear correlation statistics. Among nonparametric tests, the conditions that guarantee correct false positive rates are either restrictive or unclear. Here we describe the truncated time-shift (TTS) test, a nonparametric procedure to test for dependence between two time series. We prove that this test is valid as long as one of the time series is stationary, a minimally restrictive requirement among current tests. The TTS test is versatile because it can be used with any correlation statistic. Using synthetic data, we demonstrate that this test performs correctly even while other tests suffer high false positive rates. In simulation examples, simple guidelines for parameter choices allow high statistical power to be achieved with sufficient data. We apply the test to data sets from climatology, animal behavior and microbiome science, verifying previously discovered dependence relationships and detecting additional relationships.

## Introduction

Researchers routinely look for correlations between variables to identify potentially important relationships, or to use as a starting point for downstream modeling and experiments. In fields such as climatology, ecology, and physiology, data are often collected as time series, and so correlation analyses using time seres are common.

Interpreting a correlation between time series can be challenging because it is easy to obtain a seemingly high correlation between two time series that have no “meaningful” relationship [1, 2]. For example, the population densities of replicate exponentially-growing bacterial cultures may be correlated over time, but this correlation is driven by a temporal trend rather than any causally meaningful relationship. To avoid being fooled by spurious correlations, it helps to distinguish between the concepts of “correlation” and “dependence”, and how each relates to causation. In time series research, “correlation” is often defined procedurally [1, 3, 4]. That is, a correlation function is any function that takes a pair of time series and returns a number. We call this number a correlation statistic, and it is usually interpreted as a measure of similarity or relatedness. Examples include Pearson’s correlation coefficient, local similarity [5], and cross-map skill [6]. Whereas a correlation statistic is a description of an observed dataset, statistical dependence (or independence) is a hypothesis about the relationship between variables.

Two variables *x* and *y* are (statistically) dependent if the probability distribution of *x* while statistically controlling for *y* (the conditional distribution of *x* given *y*) differs from the distribution of *x* while not controlling for *y* (the marginal distribution of *x*). For example, lung cancer and smoking are dependent if the probability of lung cancer among smokers is different from that in the entire population. The concept of dependence applies not only to pairs of univariate variables, but also to pairs of vectors such as time series. Importantly, dependence is linked to causality (as defined in the usual sense: *x* causes *y* if perturbations in *x* can alter *y*). The link between dependence and causality is due to Reichenbach’s common cause principle, which states that if two variables are dependent, then they are causally related: Either they share a common cause, or one variable causes the other (possibly indirectly) [7, 8]. Thus, before seizing upon causal explanations, it is useful to first test whether the observed correlation is strong enough to indicate dependence.

To test against the null hypothesis of independence, two ingredients are needed: (i) a correlation statistic and (ii) a method to estimate the distribution of that statistic under the null hypothesis of independence. This article focuses on the second ingredient.

In the simpler (non-temporal) case where measurements of two variables are independent and identically distributed (iid), the permutation test provides a general way to test for dependence between the two variables. Specifically, let (*x_i_, y_i_*) be the *i*th pair of measurements of variables *x* and *y*, and let θ be the observed correlation between the *x* and *y* measurements. The permutation test randomly shuffles the index of one of the variables, and then recomputes the correlation. This process is then repeated many times, essentially producing a null distribution. Under the null hypothesis that *x* and *y* are independent, correlations obtained from the original and the shuffled data follow the same distribution. Thus, a *p*-value can be calculated as (see, for example, section 6.2.5 of [9]):

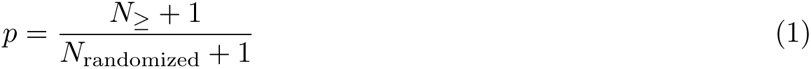

where *N*_randomized_ is the total number of “randomized correlations” (i.e. correlations obtained from shuffled data), and where *N_≥_* is the number of randomized correlations that are at least as strong as the original correlation. The “+1” terms account for the original correlation. This test has three especially desirable properties. First, the test is *valid*: If we infer dependence only when *p* is less than some significance level α, then our false positive rate (i.e. the chance of erroneously reporting dependence) will be no more than α [9, 10]. Second, the test is distribution-free: It does not require that the variables or the correlation statistic follow a particular probability distribution [11]. Lastly, the test is *without* critical parameters: Its validity does not depend on any parameters that must be estimated or chosen by the user.

Dependence testing is less straightforward when applied to a pair of time series. Although a time series can have many data points, these data are not independent of each other in the sense that they are often autocorrelated (e.g. what occurs today influences what occurs tomorrow). A permutation test carried out by shuffling data within time series will generally not be valid. This is because temporal shuffling destroys autocorrelation, which often leads to artificially weak randomized correlations and thus an unacceptably high false positive rate (Fig 2 in [12]). If multiple independent and identical systems (trials) are available, we can instead perform a valid test by comparing within-trial correlations to between-trial correlations [13, 14, 12]. However, many important questions focus on a pair of single time series only (i.e. they have the “*n*-of-one” challenge). For instance, global-scale environmental studies are necessarily *n*-of-one because replicate Earths do not exist, and the *n*-of-one perspective has been advocated in psychology because statistical patterns within one individual might differ from patterns in other individuals [15].

If a correct model of the autocorrelation happens to be known, this model can sometimes be used to remove the autocorrelation (“prewhitening”) so that standard correlation tests may be applied [16]. However, simulation benchmarks have shown that prewhitening-based dependence tests can have worryingly high false positive rates [17] and certain data types cannot be prewhitened [18].

An alternative approach to testing for dependence between time series is to use a test that takes auto-correlation into account (rather than eliminating it during data preprocessing). One can do so via either a parametric or nonparametric test. Parametric tests assume that the data follow a particular distribution (e.g. Gaussian), and use this assumption to analytically derive a null distribution for a particular statistic (such as Pearson’s correlation coefficient) [19, 20, 21, 17]. However, an analytical null distribution is not available for some increasingly popular statistics such as cross-map skill [6, 22, 23, 24, 25, 26] (see also [27, 13] for a broader overview of nonlinear dependence statistics).

When parametric tests are unavailable or inappropriate, it is common to test for dependence between time series by using a rank-based resampling approach called surrogate data testing [27]. Surrogate data tests are nonparametric (i.e. they do not rely on an analytically derived null distribution) and can thus accommodate any correlation statistic. This approach begins with two time series {*x*_1_*,x*_2_*,… x_n_*} and {*y*_1_*,y*_2_*,… y_n_*} (abbreviated {*x_t_*}and {*y_t_*}), and computes some measure of correlation between them. Next, one uses a computer to simulate independent replicates of {*y_t_*}. These simulated {*y_t_*} time series are called “surrogate” {*y_t_*} series. Finally, one computes the correlation between {*x_t_*} and each surrogate {*y_t_*}. A *p*-value is then given by the proportion of surrogate {*y_t_*} series that produce a correlation equal to or larger than the real {*y_t_*}. More precisely, the *p*-value is given by Eq. 1, but where where *N_≥_* is now the number of surrogates that produce a correlation statistic at least as large as the original {*y_t_*} series and *N_randomized_* is the total number of surrogates [28].

Several procedures have been used to generate time series surrogates, each with different strengths and limitations. For instance, the random phase procedure decomposes the {*y_t_*} time series into sine waves, randomly shifts these sine waves in time, and finally sums them up to produce surrogates [29, 23] (Appendix 3-Figure 1 of [12]). This procedure is valid when {*y_t_*} is a Gaussian process (meaning that any subsequence follows a multivariate Gaussian distribution) and is stationary (meaning that this distribution does not change over time) [27, 13]; see [30] for precise validity conditions. A more general version of the random phase procedure, called the iterative amplitude-adjusted Fourier transform (IAAFT) procedure, is valid when {*y_t_*} is a stationary Gaussian process that has been passed through an invertible but possibly nonlinear “measurement function” [28, 27]. Yet even this more general condition would seem to exclude processes that were never Gaussian to begin with. Surrogates can also be produced by a block bootstrap method in which random subsequences are selected from {*y_t_*} and joined together [31, 32]. However, junctions between the blocks can produce disruptions, rendering the test inexact [31]. A sophisticated variant of the block bootstrap method, called the twin method, attempts to position blocks so that the disruptions are minimized [33]. However, even for this method, validity depends on “embedding parameters” [34] that must be appropriately chosen by the user, a potentially difficult task [35]. Overall, such surrogate procedures do not embody the three desirable properties listed above (being valid, distribution-free, and without critical parameters).

The procedure we propose in this article is most similar to a class of “time-shift” procedures. These procedures produce surrogates by shifting the original {*y_t_*} in time [36, 37, 38, 39, 24, 32, 13]. One can in principle use these surrogates to calculate a *p* value according to Eq. 1. However, Bartlett [40] noted that such an approach is generally invalid because the surrogates are statistically dependent on each other.

Here we describe the truncated time-shift (TTS) test which tests for dependence between two time series. We mathematically prove that the TTS test is valid as long as one of the two time series is strict-sense stationary. The TTS test is compatible with any correlation statistic and its validity does not require the assumption of a particular probability distribution, nor does it require that a user correctly select some parameter. Although the statistical power of the TTS test can depend on user-selected parameters, we demonstrate that in common benchmark systems, simple guidelines for parameter choices allow high power to be achieved (as long as sufficient data are available). Lastly, we demonstrate the use of the TTS test by applying it to real data from climatology, animal behavior and microbiome science.

We note that after we uploaded an earlier version of this manuscript to bioRxiv, we happened to discover an arXiv preprint that independently conceived and proved an equivalent test [18]. The preprint, whose primary focus is a description and proof of the TTS test, additionally shows that the TTS test is not excessively conservative. That is, using the same procedure with a less stringent cutoff will always produce an invalid test for finite data (although a less stringent cutoff does become valid in the limit of infinite data). We nevertheless provide our version of the proof in Appendix 1 because (1) our proof is more complete in the sense that each statement is justified; (2) our proof applies more directly to finite-time processes, as we use a definition of stationarity that applies explicitly to finite time series (instead of stationarity in standard stochastic process literature which is defined only for infinite time series [41, 42, 43]); and (3) our proof is intended to be relatively accessible, with graphical illustrations of intermediate lemmas and relevant background concepts (Appendix 1.2).

## Results

### The truncated time shift (TTS) test

We say that the temporal processes {*x_t_*} and {*y_t_*} are independent if

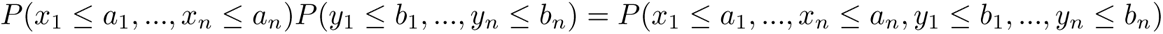

for all {*a_t_*} and {*b_t_*}. This is one of several equivalent definitions (see for instance pg. 127 in [44]).

The truncated Time Shift (TTS) test is designed to test for dependence between {*x_t_*} and {*y_t_*}, with the null hypothesis being that they are independent. The test is based on time-shifted surrogates, and requires shifting the original {*y_t_* } series in time without altering its length. One way to achieve this is to use cyclic permutations [24, 32]. That is, if the original {*y_t_* } series were {1, 2, 3, 4}, then there would be 3 surrogates, given by {2, 3, 4, 1}, {3, 4, 1, 2}, and {4, 1, 2, 3}. However, these surrogates artificially force the first and final points of the original {*y_t_* } series to become neighbors, which can distort the dynamics [27].

Instead, we will truncate time series and then shift them to generate surrogates [27]. Starting with {*x*_1_*,x*_2_*,… x_n_* }, we delete *r* time points from each end of the sequence, and obtain:

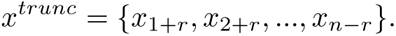

We call *r* the “truncation radius”. We then define a collection of truncated and shifted *y* time series, which all have the same length as *x^trunc^*:

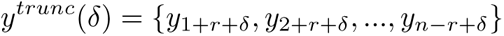

where the shift *δ* can take on integer values between *-r* and *r* (Fig 1A-B, step 1). Note that when *δ* = 0, *x^trunc^* and *y^trunc^* (*δ*) are aligned. Thus, we can think of *y^trunc^* (*δ*) as the original time series and *y^trunc^* (*δ*) for *δ ≠* 0 as the surrogate time series. For each value of *δ* between *-r* and *r*, we compute the correlation between *x^trunc^* and *y^trunc^* (*δ*) (Fig 1A-B, step 2). We then define *B* (for “Bigger”) as the number of shifts *δ* that produce a correlation at least as large as when *δ* = 0 (Fig 1A-B, step 3). *B* is bounded between 1 and 2*r* + 1: We have *B* = 1 if the strictly greatest correlation is obtained when *δ* = 0. Conversely, *B* is equal to 2*r* + 1 if the lowest correlation (or a correlation that is tied for lowest) is obtained when *δ* = 0.

**Figure 1:**
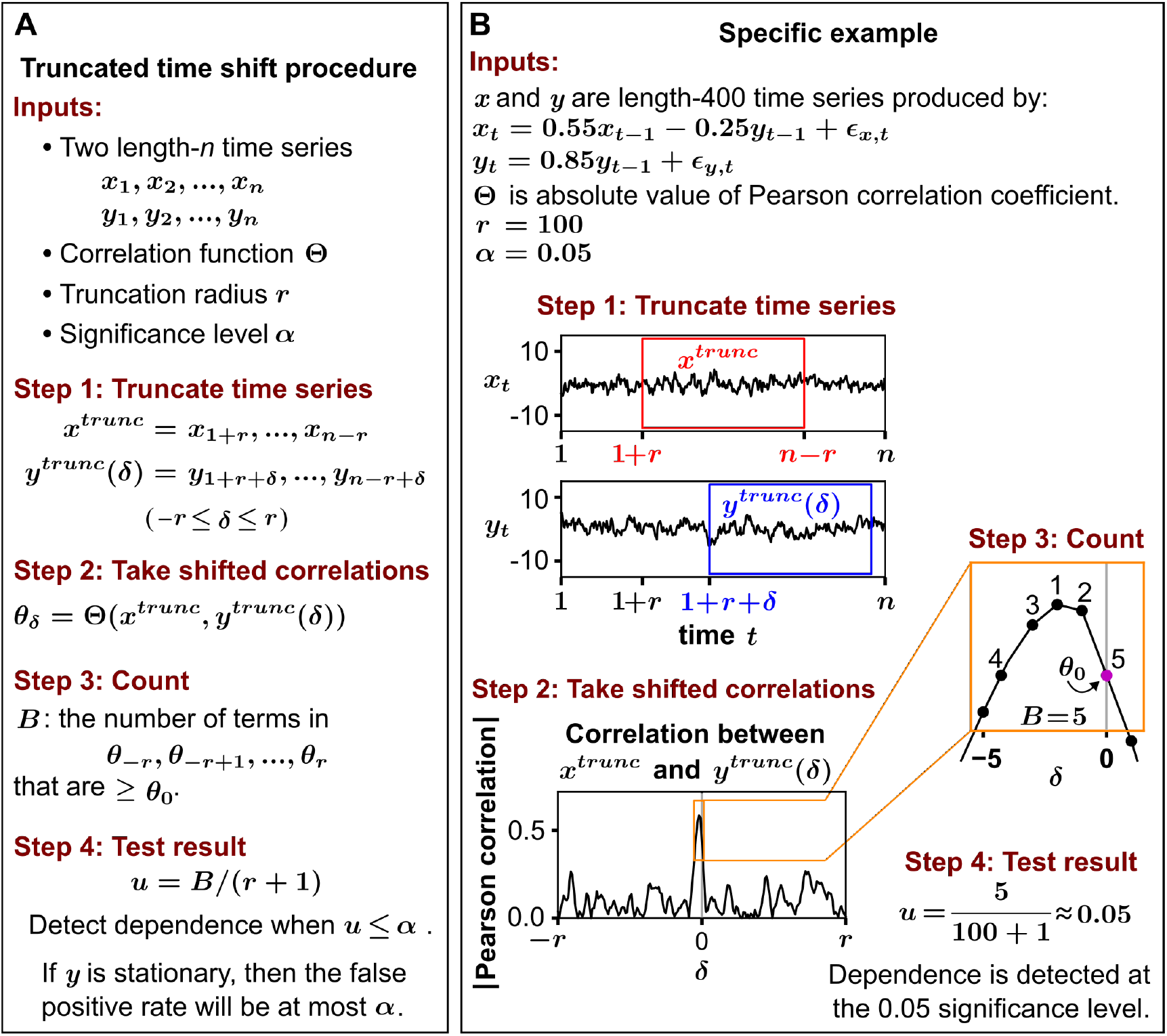
The truncated time shift procedure. (**A**) Stepwise description of the procedure. (**B**) A worked example. In the example, the process noise terms ε*_x,t_* and ε*_y,t_* are independent and identically distributed normal random variables with variance of 1 and zero mean. Note that in step 4, the statistic *u* can exceed 1 (e.g. the maximum *B* can be 2*r* + 1, in which case the null hypothesis of independence would certainly not be rejected). If one insists on reporting a *u* value between 0 and 1, then min(*u,* 1) instead of *u* can be used, giving what is sometimes called a “superuniform” *p*-value [45, 46].

If one were to naively apply the traditional logic of surrogate data testing (Eq 1), a *p*-value could be written as simply the proportion of correlations (shifted or not) that are at least as large as as the unshifted correlation:

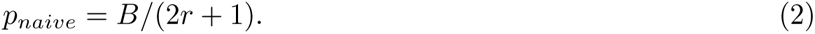

As Bartlett [40] noted, *p_naive_*is not a valid *p*-value because the surrogate *y* series are not independent of each other (e.g. two consecutive shifts are nearly identical). Instead, our approach relies upon the following statistic:

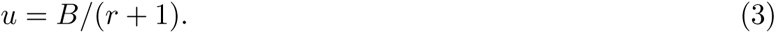

We refer to this procedure as the truncated time shift (TTS) test. Although *u* is not a *p*-value in the usual sense (e.g. *u >* 1 is possible), *u* can be used in the same way to establish evidence against the null hypothesis that {*x_t_*} and {*y_t_*} are independent. That is, if the null hypothesis is true, then the probability of *u ≤* α is no more than α (Fig 1A-B, step 4). In Appendix 1 we prove that this property holds under the assumption that *y* is stationary. Roughly speaking, a temporal process is stationary (also called strict-sense stationary) if its probability distribution does not change over time (see Appendix 1.2 for a precise mathematical definition). Stationary processes include many equilibrium processes, noise processes, chaotic processes, and periodic processes with random phases.

The above mathematical result may also provide a touch of comfort to analyses performed using the naive test: Comparing the formulas for *u* and *p_naive_* (Eq. 3 and Eq. 2), we can see that as long as the requirement of the TTS test is satisfied (i.e. the time series used to generate surrogate data is stationary), the false positive rate of the naive test will not be inflated above the significance level by more than a factor of 2. However, many applied studies do not use the naive TTS test as we have described it, but instead use a number of variations on it ([36, 38, 39, 32]). In Appendix 2, we consider two possible variants of the naive TTS test and use simulation examples to show that these can in principle be miscalibrated by far more than twofold.

### The TTS test correctly controls false positive rates when similar tests do not

Here, using simulated systems consisting of pairs of independent time series, we compare the false positive rates for the TTS test and several existing surrogate data tests, as well as several parametric tests. Whereas all other tests fail in at least one stationary system, the TTS test performs correctly in all stationary systems (as expected). The TTS test also performs well in the two nonstationary systems considered here.

Systems are shown in Fig 2A and indexed by Roman numerals. Systems i-ii are a first order autoregressive process and the logistic map, which are two stationary systems commonly used for benchmarking (e.g. [29, 6]). Systems iii-vi are four stationary systems with a combination of periodic dynamics and noise designed to challenge existing tests. Systems vii-viii are two biologically-inspired nonlinear systems: a stochastic FitzHugh-Nagumo neuronal model [47] and a competitive Lotka-Volterra system with chaotic behavior [48]. These two systems are likely to be stationary, although formal proofs are generally difficult for multivariate nonlinear systems [49]. Systems ix-x are two systems known to be nonstationary: a random walk and a first order autoregressive process (same as system i) with an additive term that increases over time. In all cases, the two time series are independent by construction. See Appendix 3 for mathematical details.

**Figure 2:**
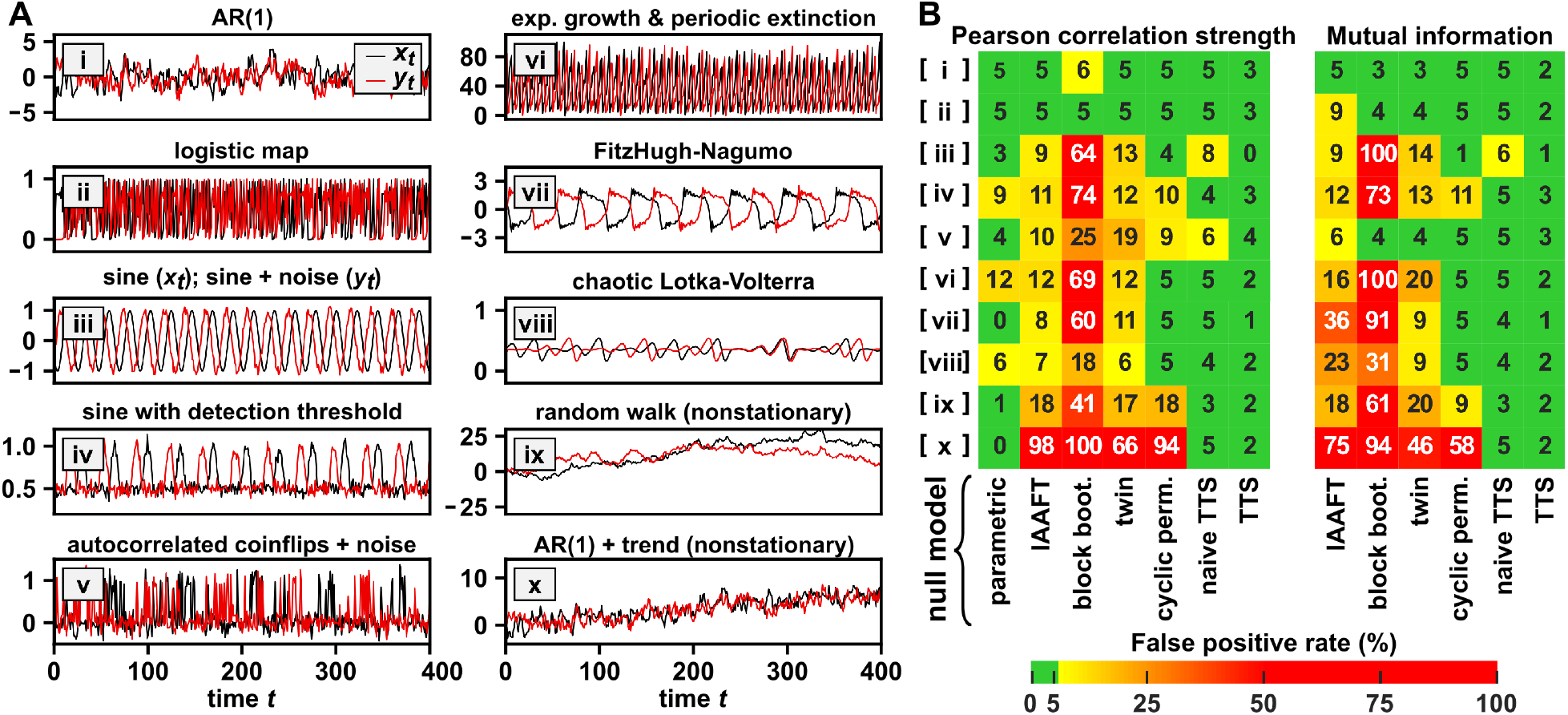
The truncated time shift (TTS) test controls the false positive rate in stationary systems and some nonstationary systems. (**A**) Example time series from different benchmark systems, some of which (i.e. systems i-vi) can be proven to be stationary (Appendix 3). Systems vii-viii are probably approximately stationary, but it is in general difficult to prove strict stationarity for multivariate nonlinear stochastic systems [49]. Systems ix-x are not stationary. A system can have process noise (noise whose effect can propagate to subsequent time steps) and/or measurement noise (whose effect does not propagate). (i) First-order autoregressive process: current values depend linearly on values one step ago and process noise. (ii) Logistic map: a deterministic discrete-time model of population dynamics with growth and a carrying capacity [54]. (iii) Two sine waves, one with measurement noise whose strength varies with a slow “sawtooth wave” [55]. (iv) A sine wave with a detection threshold and measurement noise. (v) A series of coin flips with measurement noise (with ‘heads’ and ‘tails’ coded as 1 and 0 respectively), where coins are autocorrelated because the probability of a ‘heads’ varies over time in a stationary way. (vi) A simple model of a population with exponential growth, periodic extinction events, and constant immigration. (vii) A stochastic discrete-time FitzHugh-Nagumo model, which is a nonlinear oscillator inspired by neural voltage dynamics. (viii) Chaotic Lotka-Volterra model: an ecological model where species engage in intra- and inter-species competition. (ix) A random walk with Gaussian steps (i.e. *x_t_ x_t-_*_1_ follows a zero-mean normal distribution). (x) The same process as in (i), but with an additive temporal trend. (**B**) False positive rates of dependence tests are calculated from 1.5 × 10^4^ trials. For each system, two independent time series were simulated and correlated using either the absolute value of the Pearson correlation coefficient or an estimate of mutual information. Each surrogate data test was then used to test for significance (at the 0.05 level) of the correlation under the null hypothesis of independence. The labels “block boot.” and “cyclic perm.” are shorthand for stationary block bootstrap and cyclic permutation surrogates. See Appendix 3 for further details.

Two different correlation statistics were used: Pearson correlation strength (the absolute value of the sample Pearson correlation coefficient; Fig 2B, left half), which is a linear form of correlation, and an estimator of mutual information [50], which is a popular nonlinear form of correlation (Fig 2B, right half). We do not use statistics based on the Granger causality framework as correlation statistics in this paper. This is because the Granger causality framework requires tests of conditional dependence [51], whereas the TTS test and most surrogate data procedures provide tests of (unconditional) dependence and thus are generally inappropriate for Granger causality testing (i.e. Figure 6 of [52]).

For each statistic, we compared the following surrogate data tests: IAAFT [28], stationary block bootstrap [53, 31, 32], twin surrogate test [33], cyclic permutation, naive TTS test (Eq. 2) and TTS test (Fig 1). For the first four tests, we use circularization to reduce a potential discontinuity caused by wrap-around effects (Methods), as recommended by [28, 13]. We also tested four autocorrelation-aware parametric tests: A *t*-test for Pearson correlation strength (Methods) given by [19], a modified version of the same test using a procedure suggested by [20] (Methods), and two recently described tests for Pearson correlation and mutual information respectively in linear systems [17]. In this benchmark, the latter three tests exceeded the 5% false positive rate threshold more often than the first test (and performed similarly to each other; supplementary data file 1). For this reason, we show only the original test of [19] in Fig 2, and use only this test as our parametric test for all subsequent comparative analyses.

The TTS test has a false positive rate below 5% for all stationary systems, as expected given our mathematical proof. All procedures other than the TTS test mistakenly detect dependence at rates above 5% in one or more stationary systems (Fig 2B). In Appendix 3.4, we examine possible reasons behind failure modes of the cyclic permutation procedure (a commonly used procedure from the time-shift method class). The TTS test also showed low false positive rates with the two nonstationary systems (Fig 2B, last two rows), although it is not guaranteed to be valid when surrogates are generated from a nonstationary process (e.g. Fig S11).

For time series that can be decomposed into a deterministic “trend” and a stationary component (e.g. Fig 2Ax), the TTS test can be modified to be rigorous by first detrending followed by retrending (similar to [56]). The basic idea is to: (1) extract the stationary component by removing the trend, (2) generate surrogates from the stationary component, and then (3) add the trend back to the surrogates. In Appendix 4, we demonstrate that the detrending-retrending TTS procedure produces a valid surrogate data test for time series that are nonstationary due to a deterministic trend.

In summary, whereas the previous section established that the TTS test is universally valid for dependence testing in stationary time series, here we have shown using simulation examples that popular surrogate data tests are not universally valid.

### Simple guidelines for choosing TTS parameters to achieve high statistical power

Having shown that the TTS test always correctly controls the false positive rate (as long as one series is stationary), we now consider its true positive rate (detection power; the probability of detecting a true dependence relationship). Apart from the choice of correlation statistic, the power of the TTS test is sensitive to two parameters. One parameter is the truncation radius *r*, which is specific to the TTS test. The second parameter (the pre-shift amount *s*), which we introduce later, arises from the general problem of delayed coupling (e.g. when the effect of one variable on another occurs after a lag). However, simple guidelines for choosing *r* and *s* allow high statistical power to be achieved when sufficient data are available, as we demonstrate below using simulation examples.

The truncation radius *r* simultaneously determines the length of the truncated time series (which is 2*r* less than the total number of time points) and the number of time-shifted surrogates (which is 2*r*). A rearrangement of Eq. 3 gives *r* = *B/u −* 1. If *r* is exactly 19, significance will be detected at the *u* = 0.05 level only when *B* = 1, meaning that the original correlation would need to be strictly greater than all shifted correlations. If 20 ≤ *r* ≤ 38, significance at the 0.05 level still requires *B* = 1 (since if *B* = 2 and *r* = 38, then *u* = *B/* (*r* + 1) = 2*/* 39 *>* 0.05). Thus, choosing 20 *r* 38 will never achieve higher power than setting *r* = 19. In general, for a significance level α, power is maximized when *r* is one less than an integer multiple of 1*/* α. Note that the data length needs to be more than 2*r*, and consequently more than 2*/* α. As *r* grows larger, the TTS test will be able to detect dependence with progressively greater values of *B.* However, if *r* is too large, the truncated time series will become short, causing the correlation statistic to become noisy.

Lagged dependence arises when one variable affects another after a delay, or when a common driver affects two variables with different delays (Fig 3A black curves, *x* lagging behind *y*). Detecting lagged dependence can be challenging for many statistical tests (i.e. all tests in Fig 2), because although two variables may have a strong shifted correlation (between *x_t_*_+_*_s_* and *y_t_* for some shift *s*), the unshifted correlation might be very low. The problem is especially severe for tests that rely on shifted correlations as their null models (including the TTS test). This is because the existence of a coupling lag means that some of the null (shifted) correlations will exceed the original (unshifted) correlation (Fig 3B black curve; many black dots above the red dot), potentially leading to low power.

**Figure 3:**
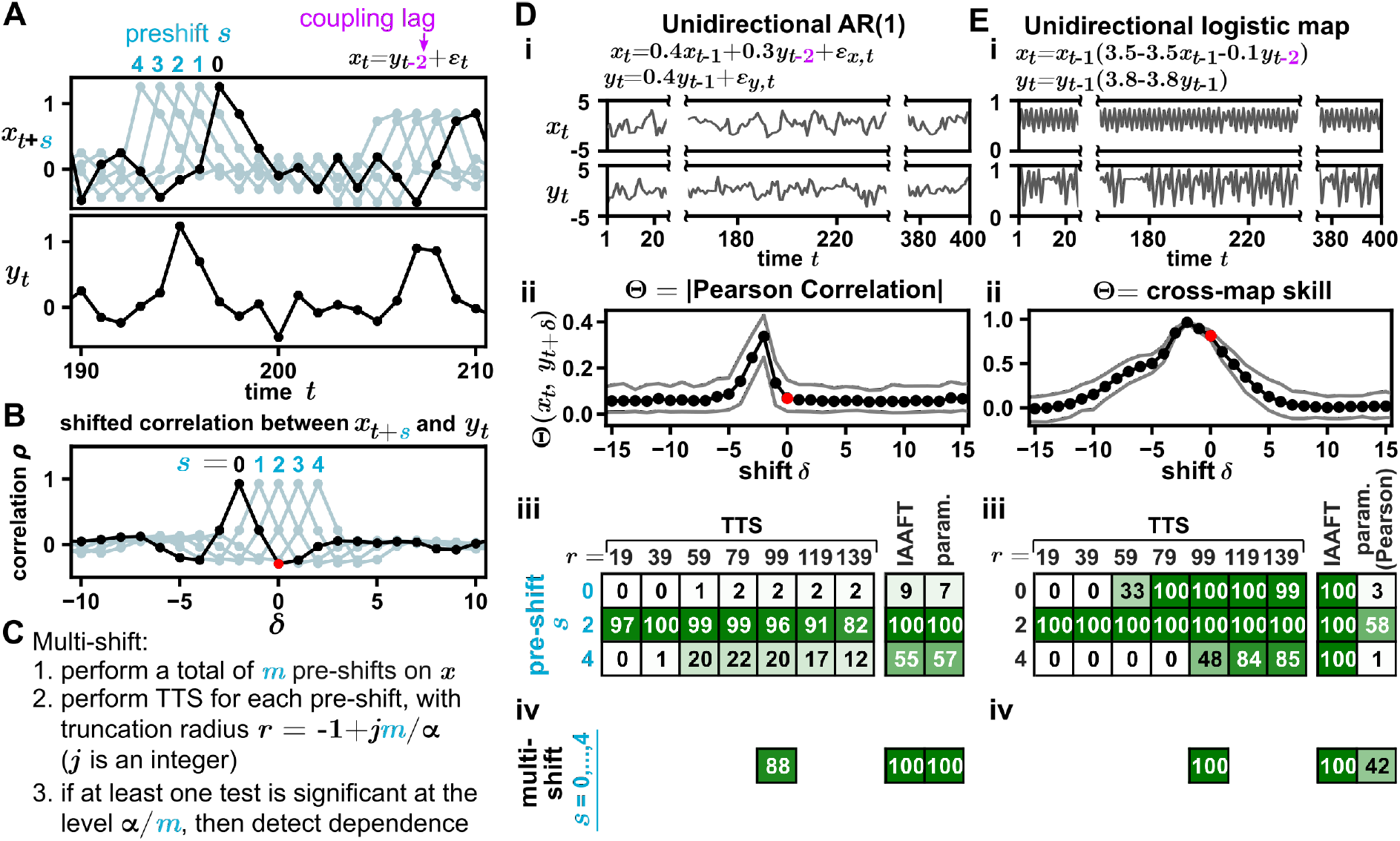
Strategies for increasing power in the challenging setting of a coupling lag. (**A-C**) Conceptual illustration of pre-shift and multi-shift. (**D, E**) Various pre-shifting strategies to account for coupling lag increase the detection power of the TTS test. (**i**) Time series in which *y* influences *x* with a lag of 2. We compare a coupled linear autoregressive process (D) and a nonlinear logistic map (E) [6, 58], which is a model inspired by ecological competition. (**ii**) Shifted correlation plots. The absolute value of the Pearson correlation (D) or cross-map skill (E) was computed between *x_t_*and *y_t_*_+_*_δ_* for various shifts δ. Detecting dependence in D is more challenging than in E since the unshifted correlation (red dot) is at the foothill in D but near the maximum in E. To calculate cross-map skill, we used *x* to estimate *y* (as appropriate when *y* influences *x* [6]), and the embedding dimension and the embedding lag were set to 2 and 1 respectively following prior works [6, 58]. Black dots indicate the mean correlation (and grey lines indicate the upper and lower 10%) from 100 trials with a truncation radius of 79. (**iii**, **iv**) Power comparison. Tests in Diii and Eiii used the correlation statistic specified in Dii and Eii respectively, except for the parametric test, which always used Pearson correlation. Prior to performing the tests, the *x* series was pre-shifted by *s* (i.e. testing for dependence between {*x*_1+_*_s_,…, x_n_*} and {*y*_1_*,…, y_n-s_}*, where *s* = 0, 2, or 4 for single pre-shift at significance level 0.05 (iii) or *s* = 0, 1, 2, 3, 4 for multi-shift at a Bonferroni-corrected significance level 0.05*/* 5 = 0.01 (iv). Detection power was computed as the proportion of 10^4^ simulations in which dependence was detected.

The problem of low power associated with lagged dependence can be mitigated by pre-shifting one of the time series before applying the TTS test (Fig 3 A, light teal curves). When the pre-shift amount is similar to the coupling lag, correlation at *δ* = 0 reaches a high value (Fig 3B, *s* = 2), enabling detection of dependence. If we know a range of likely values for the coupling lag, we can perform a “multi-shift” test: Test for dependence between {*x_t_*_+_*_s_*} and {*y_t_*} for several different values of *s*, and perform a Bonferroni correction to account for multiple tests. If *m* different pre-shifts are used, then dependence may be reported if any of the *m* tests is significant at the α*/m* level. Correspondingly, the truncation radius will need to be 1 less than an integer multiple of *m/* α (Fig 3 C). As the range of possible lags becomes larger, the minimum *r* value will increase and longer series may be needed.

To demonstrate how pre-shifting can improve the statistical power of the TTS test, we use an “unforgiving” case (an autoregressive process where *y* influences *x* with a delay of τ = 2 time steps in Fig 3Di). This process is unforgiving in the sense that the shifted correlation plot exhibits a sharp peak, and consequently the unshifted correlation is not higher than most of the shifted correlations (Fig 3Dii). Consequently, the TTS test does not obtain a significant Pearson correlation (Fig 3Diii, first row). The challenge is not entirely unique to the TTS test, as the IAAFT and parametric tests also suffer low power. Pre-shifting *x* by τ = 2 allows all three tests (including the TTS test at different truncation radius *r*) to detect the dependence with high power (Fig 3Diii, second row). If the pre-shift amount *s* is too large, the power again declines as expected, and compared to the IAAFT and parametric tests, the power of the TTS test is more sensitive to the incorrect pre-shift (Fig 3Diii, third row). The multi-shift approach described in the previous paragraph (Fig 3C) achieved high power (Fig 3Div). With sufficient data, multi-shift also allowed high detection power when two variables affect each other with different and uncertain lags, as long as the range of pre-shifts covers the maximal lag uncertainty (Appendix 5.2).

Pre-shifting may not be necessary in “forgiving” cases. One such case is a coupled bivariate logistic map (Fig 3Ei), a nonlinear system whose coupling can be readily detected using the “cross-map skill” correlation statistic [6, 23, 57]. As before, the two processes are coupled with a lag of 2. The shifted correlation plot exhibits a peak that is broader than the coupling lag (Fig 3Eii). In this case, since *B* = 4 (as three of the shifted correlations are greater than the unshifted one), a truncation radius of *r* = 4*/* 0.05 1 = 79 or above can detect dependence even without any pre-shifting (Fig 3Eiii first row). Similar to Fig 3E, other studies have also found broad peaks when using nonlinear correlation statistics to detect coupling between nonlinear deterministic systems [27, 58]. Note that it is inappropriate to visually inspect the plot of shifted correlations and then decide what pre-shift to use, since this uses test output to select test parameters and will invalidate the test. A systematic method to diagnose whether a system and/or correlation statistic will likely give rise to a “broad peak” could be a useful goal for future efforts.

In summary, although the TTS test often has lower power than other tests (Fig 3D), the multi-shift strategy can dramatically improve the power as long as sufficient data are available (Fig 3Diii; Appendix 5.2). For nonlinear systems (e.g. the logistic map), the TTS and other nonparametric tests share similar power, and are far superior to the parametric test (which suffers from the constraint of using a linear statistic). These conclusions hold when we varied parameters such as the time series length, interaction strength, and autocorrelation (Appendix 5.1).

In the next three sections, we apply the TTS test to existing data sets from climatology, microbiome science, and animal behavior science. These case studies serve as examples of real systems wherein the TTS test is sufficiently powerful to detect dependence relationships, some (but not all) of which were detected by the original authors of the data sets.

### An example from climate science

Pre-industrial climatological change is widely understood to have been driven largely by variability in the Earth’s orbit around the sun. The Earth’s rotation around the Sun is characterized by three parameters known as eccentricity, obliquity, and the climatic precession index (here simply “precession”). Eccentricity describes the shape of the orbit, which varies from nearly circular to slightly elliptical over a cycle with a period of approximately 96, 000 years (96 kiloyears or 96 kyr). Obliquity is the angle between Earth’s rotational axis and the normal to the orbital plane, which cycles over roughly 41 kyr in a band roughly bounded between 22*^0^* and 24.5*^0^*. Precession is 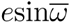 where *e* is eccentricity and 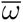 is the longitude of perihelion (the angle between the vernal equinox and the perihelion [59]), with a cycle period of about 21 kyr [60, 61]. Each of these parameters is thought to play a role in Earth’s climate, although some parameters may be more influential than others, and the extent of a parameter’s influence may change over time [62, 63, 61]. The climate record is characterized by repeated episodes of cooling followed by warming events called deglaciations. Until about one million years ago, deglaciations occurred with a period of about 41 kiloyears, which is the period of obliquity cycles. Because of this, obliquity is often said to “pace” glacial cycles [64, 65]. Yet, two time series with shared periodic elements can be statistically independent (e.g. Fig 2Aiii).

Using the TTS procedure, we tested for dependence between orbital parameters and deglaciations with only the assumption that the time series of the three orbital parameters are stationary (Fig 4A). We used the entire past 2 million years (2 Myr) of deglaciation series as our “truncated” *x* time series, and generated surrogates using orbital parameter time series spanning from 12 Myr in the past to 10 Myr in the future (i.e. with a truncation radius of 10 Myr; Fig 4A). Future orbital parameter values can be used due to the availability of an accurate physical model of Earth’s orbit [60]. We use a large truncation radius because the available orbital parameter time series are far longer than the available deglaciation time series, in contrast to Fig 3 (wherein time series have equal lengths). To test for dependence we next sought an appropriate correlation statistic. Pearson correlation seems unnatural in this context because it may be near zero (and thus fail to detect dependence) if, for exmaple, deglaciations occur at the midpoints between peaks and troughs of orbital series. To avoid this problem, we used a prediction-based nonlinear correlation statistic (similar but not identical to the cross map skill statistic used in Fig 3E; see Appendix 6.2 for more details). Using this statistic, the TTS test detected a dependence between deglaciation times and obliquity (*u <* 0.05), but not the other two orbital parameters (Fig 4B), similar to Huybers’ original period-based analysis [65].

**Figure 4:**
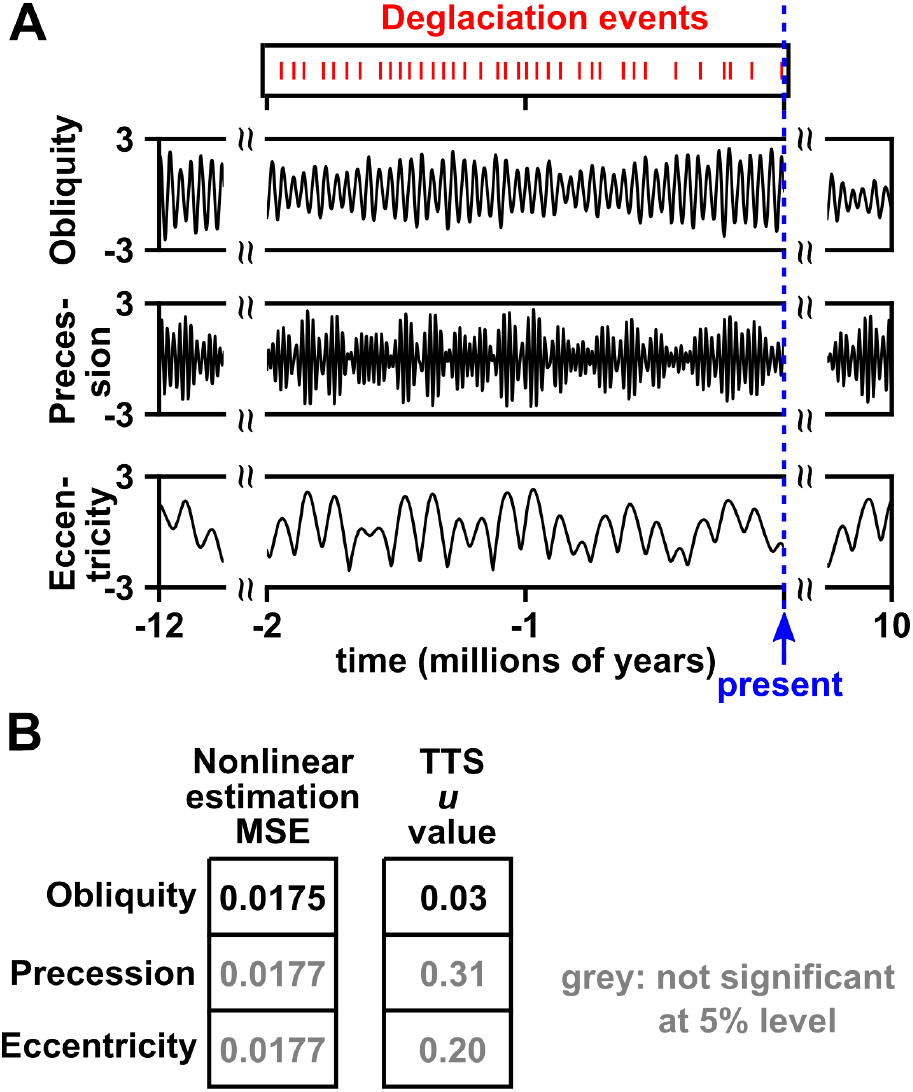
The TTS test detected dependence between deglaciation and obliquity, but not between deglaciation and precession or eccentricity. (**A**) Time series of deglaciation events (from [65]) and the three orbital parameters (estimated from the model in [60]). To convert the 36 deglaciation events ([65]) in the last 2 million years, we used a “sampling frequency” of 1 kyr by assigning a 1 to the deglaciation variable if a kiloyear contained a deglaciation event and a 0 otherwise. Our deglaciation time series thus has 2000 time points. We did not use a higher sampling frequency due to uncertainty in deglaciation timing, and avoided a lower sampling frequency (e.g. 10 kyr) to adequately capture the shapes of the obliquity and precession cycles. To estimate obliquity, precession, and eccentricity, we used the numerical solution from [60], which provides accurate estimates of orbital parameters over at least tens of millions of years (and also predicts future values). Orbital values are standardized to a mean of 0 and variance of 1. To obtain unshifted correlations, we used truncated time series with times between 1999 kyr and 0 kyr (present time), yielding a total of 2000 time points. Orbital parameter time series were used to generate time-shifted surrogates, with a truncation radius of 10 million years (20, 000 time-shifted surrogates). (**B**) Testing for dependence between orbital parameters and deglaciation. The correlation statistic for the TTS test is the mean squared error (MSE) when using orbital parameters to estimate deglaciation events via a state space-based technique. This nonlinear statistic is similar to the cross-map skill statistic used in other largely deterministic nonlinear systems (Appendix 6.2).

We did not pre-shift time series. The temporal lag between orbital parameters and temperature has been estimated in several studies [66, 67, 68]. For instance, Imbrie et al. [66] found that during the last half-million years, ice minima tended to lag fluctuations in orbital parameters by about 9 and 6 kyr for obliquity and precession respectively. However, using an empirically estimated lag as a pre-shift for the TTS test would be “cheating” and could invalidate the test. On the other hand, our simulations (Appendix 6.3) suggest that the statistical power of our test might not be especially sensitive to most pre-shifts less than or equal to the lags obtained by Imbrie et al.

### An example from human microbiome science

The human microbiome is spatially structured, with different body sites playing host to distinct microbial communities [69]. Transfer of microbes among body sites is thought to have important consequences for human health and disease [70]. Here we apply the TTS test to data from a year-long daily microbiome survey [71] to look for dependence relationships among the local microbiomes at different sites on the body of a healthy human subject. Such dependence relationships would be consistent with cross-site microbial transfer.

Importantly, we focus in this example on whether there is any detectable migration among body sites, rather than studying the migratory status of each individual species. This is because many microbiome surveys (including the one analyzed here [71]) measure only the relative abundance of different species, and this makes it difficult to perform species-level analyses (Fig S14; see also [72]). Put simply, the relative abundances of independent species will be dependent (since they sum to 1). However, testing for overall dependence among body sites is not impeded by the nature of relative abundance data. Intuitively, this is because if the absolute abundances of microbial species on two body sites are independent, then the relative abundances must also be independent. As this statement’s contrapositive, if the relative abundances of microbial species on two body sites are dependent, then the absolute abundances must also be dependent (see Appendix 7.2 for a formal argument).

We applied the TTS test to the time series from [71] to look for dependence between the microbial communities living on the left palm, right palm, tongue, and gut. We first obtained OTU (operational taxonomic unit) relative abundance tables from the data in [71] using the online Qiita platform [73] (see Appendix 7.6). We then preprocessed data (Fig 5A) by removing or filling gaps in time series, and by removing OTU abundance time series that were either mostly absent or likely nonstationary. Analyzing nonstationary processes is an important problem, but requires strong assumptions, and is outside the scope of this example. After pre-processing, the number of remaining OTUs ranged from 180 (tongue) to 507 (right palm). Finally, we performed a TTS test between each pair of body sites. Note that this example deviates from earlier examples in that both the {*x_t_*} and {*y_t_*} time series are now multivariate (due to the existence of multiple species; Fig 5B), but the TTS test still applies (Appendix 1).

**Figure 5:**
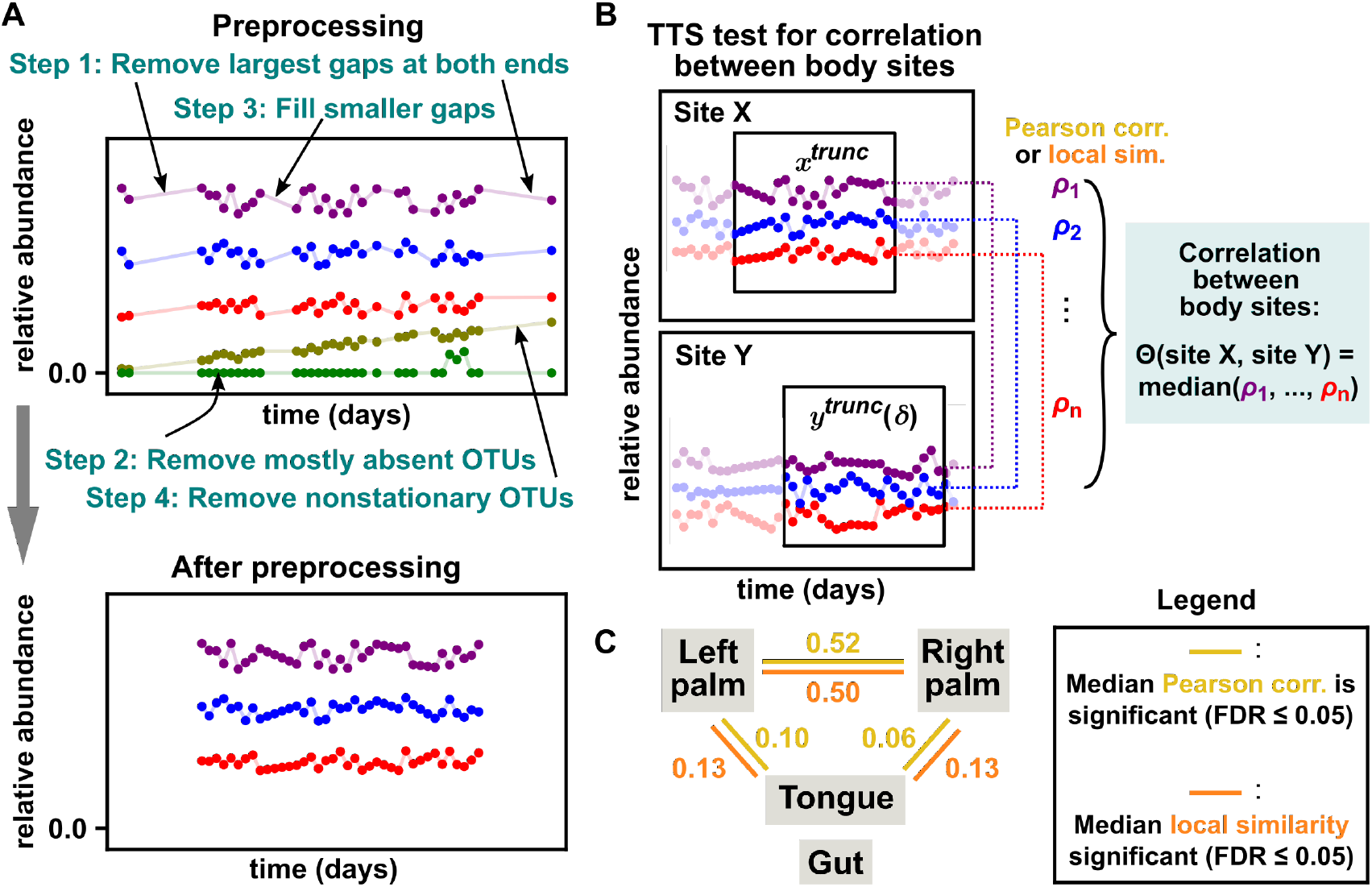
The TTS test applied to longitudinally sampled microbiomes from four body sites detects dependence between body sites. (A) Data preprocessing. To remove long (*>* 6-day) gaps at the beginning and end of time series, we only used measurements from day 42 to day 418. Remaining gaps were filled by linear interpolation or by randomly resampling abundance values. OTUs were removed if they were absent in over half of measurements or if they were not considered stationary (at the 0.05 significance level) by an augmented Dickey-Fuller (ADF) test implemented in the Statsmodels Python package [75]. OTUs removed from one body site were not necessarily removed from the other body sites so that correlations could still be computed between the other sites. (B) TTS test procedure for correlation between body sites. We used an intermediate truncation radius *r* of 79 days (inspired by Fig 3C). We quantified correlation between two body sites as follows: For each shared OTU *i*, we computed ρ*_i_* (the Pearson correlation or local similarity score of OTU *i* between the two sites). We then chose the median of {ρ_1_*,…,*ρ*_m_*} (where *m* is the total number of shared OTUs) to be the between-site correlation (i.e. our “Θ” in the notation of Fig 1). This setup avoids the need to perform a separate test for each OTU. (C) The TTS test detected dependence between the two palms and between palms and the tongue. Numbers denote the median intraspecies correlation as measured by Pearson correlation (in gold) or local similarity (in orange) when gaps were filled by linear interpolation. All links shown were detected with a significance level of 0.05 after a Benjamini-Hochberg false discovery rate (FDR) adjustment for performing 6 tests [76, 77]. Note that the gut shares few species with the other sites (Fig S16). The same network of significant correlations was obtained for (C) regardless of either the gap-filling method in (A) or whether Pearson correlation versus local similarity was used in (B). Additionally, for each correlation, we obtained the same result regardless of which body site was used to generate surrogates. See Appendix 7.6 for more details.

To correlate datasets from two body sites (Fig 5B), we first listed all of their shared OTUs. Then for each shared OTU, we computed the sample Pearson correlation coefficient between the relative abundance series of that OTU in the two sites. Our correlation statistic Θ was the median of these correlation coefficients across all shared OTUs. This statistic has the natural interpretation as the cross-site correlation for the “typical” species. Moreover, in simulations that capture the expected properties of the microbial time series (e.g. microbe transfers between sites occurring more frequently than once-per-day sampling frequency; relative abundance data), correlations tend to be positive and thus taking a median does not cancel out positive and negative correlations (Appendix 7.3). Similarly, we did not pre-shift time series because we expect dependence among body sites to be largely driven by migration on time scales likely faster than the sampling period of one day.

We detected dependence between the microbial communities on the left and right palms, and between the palms and the tongue (Fig 5C). This result was the same if instead of Pearson correlation, we used local similarity, a correlation statistic designed to detect transient temporal correlations that is popular in microbiome science [5, 74]. These results reveal more cross-site dependences than the original analysis of [71], which detected dependence only between the left and right palms. The original study first computed the phylogenetic distance between temporally adjacent microbiomes within body sites, and then calculated correlations between the phylogenetic distances of different body sites [71]. Whereas that analysis relied on parametric tests with assumed null distributions, our analysis relies on correlation statistics (i.e. the median of many Pearson correlation coefficients or local similarity scores) whose parametric sampling distributions are, to our knowledge, unknown. Yet, due to the flexibility of the TTS test, we are nevertheless able to perform a valid statistical test, assuming only that abundance time series are stationary.

### An example from animal behavior science

A major goal of animal behavior research is to understand the rules that govern how animals act, at the levels of both individuals and groups (e.g. swarms of insects or shoals of fish). Video tracking techniques enable measurements of variables such as an individual’s position and velocity [78]. These quantitative measurements have helped researchers detect subtle or complex behaviors, enable useful analogies between animal group behavior and materials physics [79, 80], and connect individual-level and group-level phenomena [78, 81, 82, 83].

We applied the TTS test to a set of zebrafish trajectories recorded by Romero-Ferrero et al. [84]. 100 juvenile zebrafish were placed in a shallow circular tank and tracked by an overhead video camera at 32 frames per second, yielding 2-dimensional trajectories of each fish (height was not measured; Fig 6A). We here use the TTS test to ask whether there is a dependence between the speed of a fish and its direction of motion.

**Figure 6:**
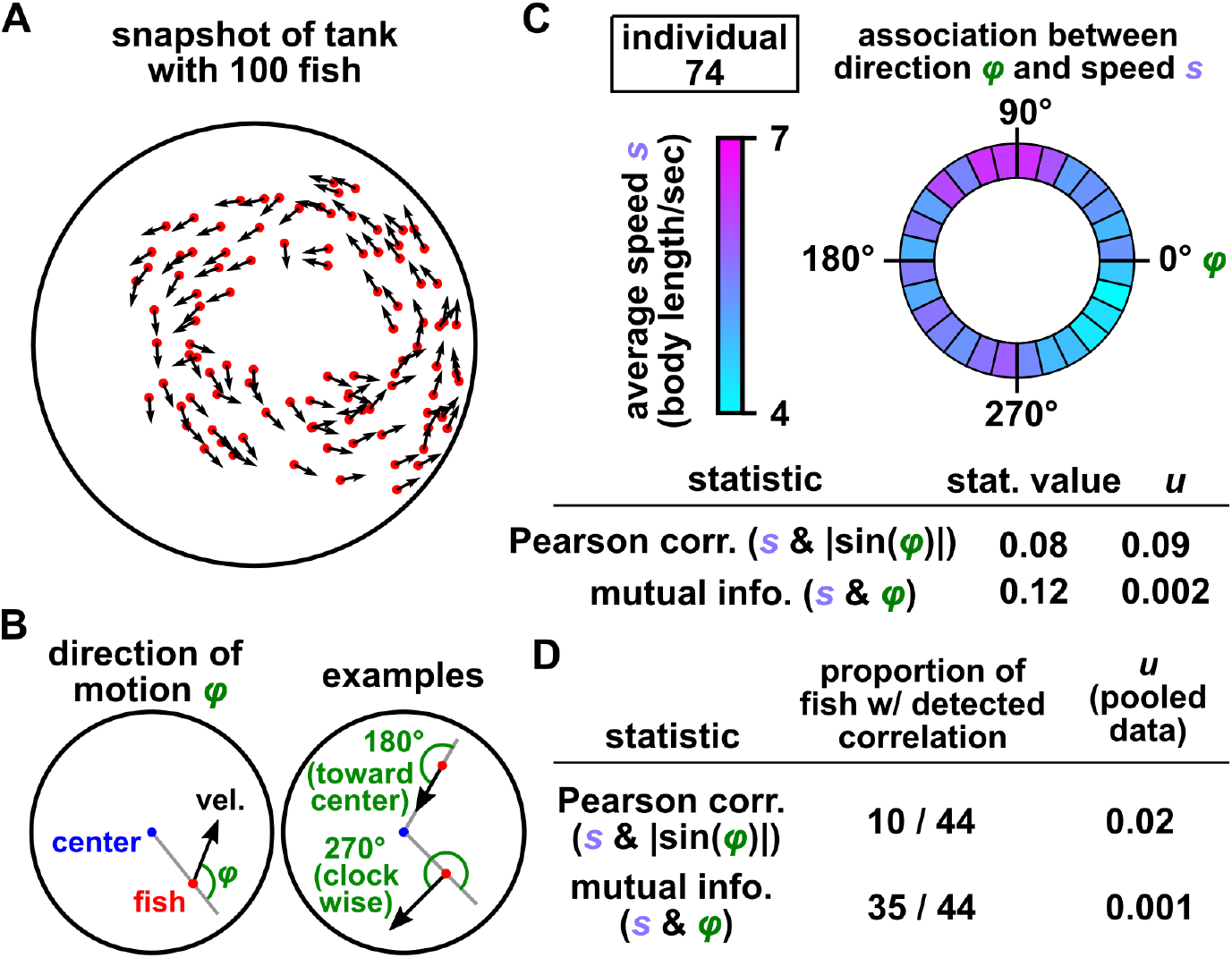
(A) A snapshot of fish positions in a 100-fish tank. (B) A fish’s direction of motion (*φ*) is defined as the angle between the fish’s position vector and velocity vector. Direction values of *φ* = 0*^0^* and 180*^0^* correspond to motion exactly away from the center or toward the center respectively. Direction values of *φ* = 90*^0^* and 270*^0^* correspond to motion exactly counterclockwise and clockwise respectively. (C, D) Association between direction (*φ*) and speed (*s*) for an arbitrary individual (“individual 74”) and for 44 fish in which recorded trajectories did not contain gaps. See Fig S18 for time series. The TTS procedure tests for dependence using the absolute value of the Pearson correlation coefficient between *s* and |sin(*φ*)|, or using the mutual information between *s* and *φ*. Mutual information detects correlations more readily in this case. When testing for correlations in 44 fish (D), detections were made at a significance level of 0.05 with a Benjamini-Hochberg FDR correction [76, 77] (middle column). We also performed a TTS test that incorporated all 44 trajectories (right column; “pooled data”). This test was analogous to the test of Fig 5B. That is, for each time shift, we calculated the overall correlation between speed and direction as the median correlation among all 44 trajectories. For all tests we used data from the first of the three replicate videos in [84] and limited our analysis to the first 10^4^ frames (about 5 minutes) since this data segment appeared approximately stationary by visual inspection, although fish speeds are likely nonstationary overall (see Appendix 8.2 and [85]). We used the speed variable for time-shifted surrogates and used a truncation radius of one tenth the total time series length (1000 frames). See Appendix 8 for further details.

To define a coordinate system for direction of motion, we noticed that large groups of fish often swam parallel to the perimeter of the tank (Fig 6A). To capture this behavior, we define an individual’s direction of motion *φ* as the angle between the individual’s position vector and velocity vector (Fig 6B).

For an arbitrary fish, different directions appear to correspond to different speeds (Fig 6C). It is unnatural to quantify this association using the Pearson correlation between speed and direction *φ* because speed is a linear quantity whereas direction is a circular quantity. We can make linear correlation more appropriate by first transforming the direction variable *φ* to |sin(*φ*)|, which is the largest when *φ* = 90° or 270*^0^* (swimming parallel to the perimeter), and the smallest when *φ* = 0° or 180° (swimming away from or toward the center), and then compute the Pearson correlation between |sin(*φ*)| and speed. Alternatively, we can use mutual information as our statistic, which is not limited to linear dependence. Fig 6C shows the statistic values and TTS test results for one arbitrary fish. For this fish, mutual information between speed *s* and direction *’* is significant, while Pearson correlation between *s* and |sin(*φ*)| is not (Fig 6C). Overall, the TTS test detected dependence between speed and direction in 10 out of 44 fish using Pearson correlation and in 35/44 using mutual information (at the 0.05 level after a Benjamini-Hochberg false discovery rate correction), respectively (Fig 6D). The TTS tests with either statistic detected dependence in pooled data (Fig 6D). We did not pre-shift time series because we had no prior expectation of a coupling delay between swimming speed and direction.

## Discussion

A statistical hypothesis test for dependence between two time series requires a correlation statistic and a null model. These two ingredients seem to have received different levels of attention over the past couple of decades. Recent years have witnessed the development and rapid adoption of new correlation statistics that can detect transient or nonlinear forms of dependence, some of which even attempt to infer the direction of causation [5, 74, 6, 58, 86, 13, 12]. In practice however, these correlation statistics have often been paired with inappropriate surrogate data null models. For example, analyzing an arbitrary non-Gaussian time series with the popular IAAFT surrogate procedure can be dangerous (Fig 2B rows ii-viii, IAAFT columns; see also [27, 87]). Even more concerning, studies have tested dependence hypothesis in time series using a naive permutation test, which assumes that a time series consists of independent and identically distributed data. Overall the general problem of assigning statistical significance to nonlinear correlations between time series does not appear to have a broadly-accepted solution [87, 88, 8].

The TTS can serve as a (provably) conservative solution to this problem since it is valid as long as one of the time series is stationary (or can be made stationary by detrending; Appendix 4). This is a minimally restrictive requirement among valid nonparametric tests of dependence between time series. This test was sufficiently powerful to verify the previously observed dependence between obliquity and deglaciation timing (Fig 4) as well as dependence between the microbiome compositions of the left and right palms (Fig 5). In the microbiome dataset, we could even use it to identify additional relationships that went undetected by the original analysis of Caporaso et al. [71]. Importantly, the TTS can be applied in cases where linear parametric tests are particularly unnatural such as when correlating fish swimming speed (which lies on the number line) to direction (which lies on the circle between 0 and 2⇡). Since surrogate data tests are presently used in disciplines ranging from Earth System science to physiology [29, 37, 36, 38, 23, 89, 13, 25, 90], we expect the TTS test to find utility in diverse application domains.

Although the TTS test is one-tailed, it may still be used to detect high or low correlations (or both). For example, if the correlation function is the Pearson correlation coefficient, then correlations are deemed “impressive” if they are positive and large. Alternatively, the demonstrations in this article primarily used the absolute value of the Pearson correlation coefficient, for which correlations are deemed impressive if they are extreme (i.e. large in magnitude, whether positive or negative). A third possibility is to simply carry out two TTS tests, one using the Pearson correlation and one using the negative Pearson correlation, and then perform a Bonferroni correction for two tests. If the distribution of correlations is symmetric and centered around zero, then the last two strategies are similar.

The main limitation of the TTS test relative to other tests is that the TTS test tends to require more data or stronger coupling to achieve high detection power. Under the TTS test, the required length of the time series increases with the reciprocal of the desired significance level. Even more data are required if the multi-shift procedure is used to handle an uncertain optimal coupling delay. Moreover, whereas other surrogate data tests often specify the desired false positive rate (at least approximately), the TTS test only specifies an upper bound on the false positive rate, and the actual false positive rate is often substantially lower (Fig 2). We can therefore expect other tests for dependence to typically achieve high detection power with fewer data than the TTS test. Indeed, this is what we have found in numerical experiments (Appendix 5). Additionally, since the naive TTS test seems to usually perform well (Fig 2), we think it is reasonable for authors to choose the naive TTS test for stationary time series, provided they note that under pathological conditions the naive TTS test may underestimate the false positive rate by up to twofold.

Since the stationarity assumption is so central in dependence testing, it would be convenient to have a statistical test for stationarity. However, a single time series can in theory be described by either a stationary process (e.g. Fig S19A) or a nonstationary one (e.g. Fig S19B). Nevertheless, there are many statistical methods that attempt to test for stationarity or a similar property in a single time series, albeit with various modeling assumptions or other caveats [91, 92, 93, 94]. For example, although the popular augmented Dickey Fuller (ADF) test [91, 94] is sometimes used as a pragmatic means of assessing stationarity (e.g. Fig 4; [95, 96]), its null hypothesis is not exactly nonstationarity. In fact, a rejection of the ADF test’s null hypothesis indicates that a time series is free from some sources of nonstationarity (e.g. a random walk), but other sources of nonstationarity (e.g. time-varying parameters) may in principle still be present. Overall, statistical tests cannot guarantee that a time series is stationary, but can provide supporting evidence. Background knowledge of the process can also be used to support the stationarity assumption: Mathematical work has shown that the stationarity condition is often met by stochastic processes that tend to relax toward a stable equilibrium [97]. Periodic processes, measurement noise processes, and chaotic processes can also be stationary (e.g. Fig 2B), as can combinations of these processes.

The subject of conditional dependence has been conspicuously absent from our discussion. Tests of conditional dependence (i.e. whether two variables are dependent after we statistically control for a third variable) can help to rule out possible common-cause explanations, and can sometimes even be used to reveal the direction of causation [98, 12]. We initially motivated the TTS test by noticing that it enjoys the rigor and generality of the permutation test, but applies to time series rather than iid data. Could a test for conditional dependence between time series be devised with the same rigor and generality? This seems difficult. Even for continuous iid data, it has been proven under general conditions that no valid test for conditional dependence can both avoid making assumptions and have statistical power [99]. Thus, we expect most tests of conditional dependence among time series to be less rigorous or require more assumptions than the TTS test. Nevertheless, there have been promising recent advances on this front, such as a test based on constrained shuffling [100, 101] and the “knockoff” testing approach, which has been recently applied to sequential data [102, 103]. Exploring methods that can robustly test conditional dependence between time series is an important future direction.

In sum, an important and longstanding problem is that of nonparametrically testing for the statistical significance of a correlation between autocorrelated datasets such as time series. The TTS test provides an approach that imposes relatively minimal requirements onto both the correlation statistic and the data-generating process. This gives researchers the freedom to apply a large arsenal of correlation statistics to a wide array of processes without sacrificing test validity. This freedom will become more valuable in the future as both correlation techniques and data availability continue to proliferate across diverse fields of research.

## Acknowledgement

We are grateful to Sean Gibbons, Benjamin Kerr, Mark Maslin, Sarah Holte, and Nathan Kutz for valuable discussions and suggestions. We also thank members of the Shou Lab for critiques of the presentation of ideas.

## Methods

### Surrogate data tests for comparative benchmarks

For surrogate data tests based on the IAAFT, stationary block bootstrap, or twin procedures, we used 499 surrogates for the empirical null distribution, unless otherwise specified. Custom Python scripts were used to generate stationary block bootstrap surrogates, cyclic permutation surrogates, TTS surrogates, and naive TTS surrogates. We used the Pyunicorn package [104] to generate IAAFT and twin surrogates.

For IAAFT surrogates [28], we used the ‘refined_AAFT_surrogates’ function in the Pyunicorn package. We set the ‘n_iterations’ argument to 200 and the ‘output’ argument to ‘true_spectrum’. For stationary block bootstrap surrogates, we set the parameter known as *p* in [53] to 0.05. This parameter sets the average block length, which is approximately 1*/p*. Cyclic permutation surrogates were generated by shifting in time with a wraparound. If the original time series was of length *n*, then *n −* 1 cyclic permutation surrogate were produced. For example, if the original time series was {*x*_1_*,x*_2_*,x*_3_*,x*_4_}, then there would be 3 cyclic permutation surrogates: {*x*_2_*,x*_3_*,x*_4_*,x*_1_}, {*x*_3_*,x*_4_*,x*_1_*,x*_2_}, and {*x*_4_*,x*_1_*,x*_2_*,x*_3_}.

For twin surrogates [33], we used the ‘twin_surrogates’ function in the Pyunicorn package. This function requires four parameter arguments: the minimum temporal distance between twins, the delay vector lag, the delay vector embedding dimension, and the “recurrence threshold”. We set the minimum distance between twins to 1. To choose the delay vector lag and embedding dimension, we used the function ‘takens_embedding_optimal_parameters’ in the Python package giotto-tda [105]. Briefly, this function first selects the delay lag τ so that the mutual information between values τ steps apart is minimized. The function then selects the embedding dimension by the widely used ‘false nearest neighbors’ algorithm, which is difficult to describe concisely, but is explained clearly by its inventors [106]. The function “takens_embedding_optimal_parameters” requires two arguments, the maximum delay lag and the maximum embedding dimension, and these were both set to 8. Finally, we chose the recurrence threshold parameter by a method advocated by the original authors of the twin procedure [33], which is to select the recurrence threshold that causes the recurrence plot to have between 5% and 20% “black points”. We used 12% black points as this is in the middle of the recommended range.

Following previous works [28, 13], we preprocessed time series to reduce a potential mismatch between the earliest and latest times before applying certain surrogate data tests. We used this preprocessing step (henceforth called ‘circularization’) for four surrogate procedures: IAAFT, cyclic permutation, stationary bootstrap, and twin. Circularization is recommended for IAAFT surrogates to avoid artifacts due to the periodic nature of Fourier components, and it is recommended for cyclic permutation surrogates because they directly join the beginning and ends of the time series [13]. We also used circularization for the stationary bootstrap and twin methods because these techniques also sometimes join the extremes of the time series.

To circularize a time series {*y*_1_*,y*_2_*,…, y_n_*}, we truncate it to {*y_kstart_, y_k__start_*_+1_*,…, y_k__end-_*_1_}, where *k_start_* and *k_end_* are chosen to minimize the mismatch between the beginning and end of the truncated time series using a formula quoted in [13]:

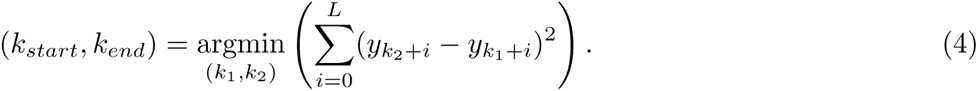

We used *L* = 10. Additionally, to ensure that the circularized time series is not too short, we require that *k*_1_ and *k*_2_ be near the beginning and end of the time series respectively. Specifically, we impose the constraints *k*_1_ ≤ 40 and *n − L − k*_2_ + 1 ≤ 40.

When circularization was used, *k_start_*and *k_end_*were chosen based on the time series that was used to generate surrogates, but both the *x* and *y* time series were circularized using the same choice of *k_start_*and *k_end_*. Additionally, both the original and surrogate correlations were calculated from circularized time series. Circularization generally improved the false positive rates of tests (compare Fig 2 to Fig S7).

### Parametric significance test

We used a parametric test described by [19]. Under the null hypothesis that two stochastic processes {*x*_1_*,…, x_n_* } and {*y*_1_*,…, y_n_* } are independent, this method estimates the variance of the sample Pearson correlation coefficient (denoted 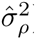) as:

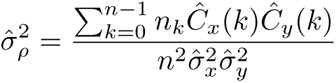

where 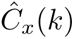 is the estimated autocovariance of the time series *x* at lag *k*:

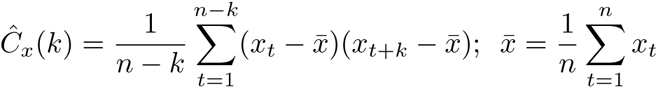

and 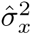 is the estimated variance of time series *x*:

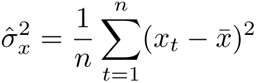

and *n_k_*is the number of entries *A_ij_*in an *n*-by-*n* matrix *A* such that *|i - j |* = *k*. In other words, *n_k_*= *n* if *k* = 0 and *n_k_*= 2(*n - k*) if 0 *< k < n*.

Since 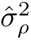 is estimated from finite data, it can under some circumstances be negative, which is nonsensical. If this occurs, 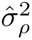 is simply set to 1*/n*, as is often recommended [19, 107, 20]. Note that 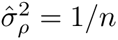 corresponds to the case without autocorrelation (to see this, plug into the equation for 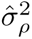 the following: 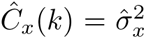 if *k* = 0 and 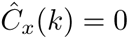 otherwise).

Next, the autocorrelation-corrected “effective sample size” is given by 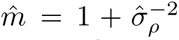, and a standard *t*-test of the Pearson correlation is performed using 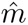 in place of *n*. That is, the test statistic is 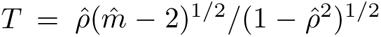, where 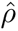 is the sample Pearson correlation coefficient, and a two-tailed *p*-value is computed by comparing *T* to a Student’s *t*-distribution with 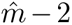 degrees of freedom [19]. This test is what is referred to as the “parametric test” unless otherwise stated.

In supplementary data file 1, we compare the false positive rate of this test to those of three other parametric tests. In that file, “Test 1” refers to the test described above. “Test 2” refers to a variant of this test in which 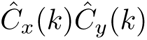 is set to zero for *k > n /* 4, as suggested by [20, 21] (but otherwise, all steps are the same as in Test 1). The rationale for this is that estimating 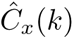 for large values of *k* is difficult. “Test 3” and “Test 4” refer to recently described tests from [17] for Pearson correlation and mutual information respectively.

### Mutual information

To estimate mutual information (except in Fig 6; see next paragraph), we used the internal function ‘_compute_mi_cc’ in the Scikit-Learn package [108] (available at: github.com/scikit-learn/scikit-learn/ blob/2beed55847ee70d363bdbfe14ee4401438fba057/sklearn/feature_selection/_mutual_info.py#L18), which implements “estimator *I*^(1)^” of [50]. The estimator requires a choice of distance metric for each of the variables being correlated, and one parameter (the number of neighbors). We used 3 neighbors and regular (i.e. Euclidean) distance for both variables.

For the fish behavior example in which we correlated speed with direction (Fig 6), the circular nature of the direction variable required a slightly different mutual information estimator. Specifically, we again used the *I*^(1)^ estimator of [50] with 3 neighbors and used Euclidean distance for speed. For direction, we used angular distance so that, for instance, the angles of 0.1π and 1.9π would have a distance of 0.2π rather than 1.8π. Mathematically, we took the distance between two direction angles α and *β* to be:

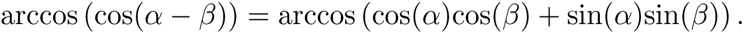

This restricts the range of possible angular distances to (0, π). We implemented the mutual information estimator using this distance metric as a custom Python script accelerated with the Numba compiler [109].

