## Appendix for "A rigorous and versatile statistical test for correlations between time series"

**9 Difficulty of testing for stationarity 65**

### 1 Mathematical justification of the truncated time-shift test

Appendix 1.1 provides a justification for the claim that the TTS test correctly controls the false positive rate as long as one of the time series under study is strictly stationary. Appendix 1.2 defines background terms (e.g. "stochastic sequences", "stationarity", and "measurability"), and provides auxiliary results that support the arguments in Appendix 1.1.

#### 1.1 Proof of the truncated time-shift test

To make this proof more accessible, we have replaced some of the most tedious aspects of mathematical notation with two terms: "$r$-neighborhood" and "$r$-locally top-$b$". We begin with definitions for these terms.

**Definition 1** *$r$-locally top-$b$*

- For a point $a_k$ in a sequence, the "$r$-neighborhood" of $a_k$ is the subsequence $\{a_{k-r}, ..., a_{k+r}\}$. That is, $a_k$'s $r$-neighborhood contains all points that are no more than $r$ steps away from $a_k$.

- A point $a_k$ in a sequence is "$r$-locally top-$b$" if $a_k$ is among the top $b$ points within $a_k$'s $r$-neighborhood. That is, $a_k$ is $r$-locally top-$b$ if no more than $b$ points in $a_k$'s $r$-neighborhood (including $a_k$ itself ) are greater than or equal to $a_k$. The edge case of $a_k$ being $r$-locally top-0 (i.e. at most zero points in $a_k$'s $r$-neighborhood are at least as large as $a_k$) never occurs.

Fig S1A-B illustrates these definitions and how they behave in the presence of ties. Note that in order for these definitions to make sense, the $r$-neighborhood must not "fall off the edge" of the sequence. For instance, if our sequence is $\{a_1, a_2, ..., a_{10}\}$, then it does not make sense to talk about the 3-neighborhood of $a_8$, as this would contain $a_{11}$, which does not exist. To avoid this problem, when writing about the $r$-neighborhood of an element $a_k$ of a sequence $\{a_1, ..., a_m\}$, we will require $1 + r \leq k \leq m - r$.

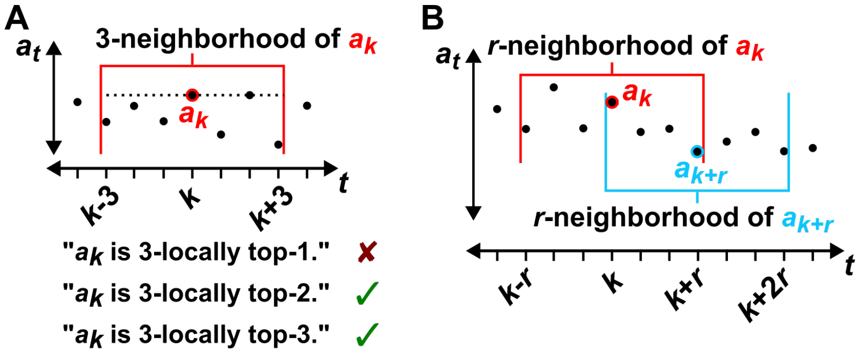

Figure S1: Illustration of "$r$-locally top-$b$". (A) The 3-neighborhood of $a_k$ includes $a_k$ itself as well as the three points to the left or right of $a_k$. In this example, $a_k$ is not 3-locally top-1, but is 3-locally top-2, 3-locally top-3, 3-locally top-4 etc. (B) $a_k$ is the leftmost point whose $r$-neighborhood contains $a_{k+r}$ and $a_{k+r}$ is the rightmost point whose $r$-neighborhood contains $a_k$. Thus, the $r$-neighborhood of any point between $a_k$ and $a_{k+r}$ contains the sequence $\{a_k, ..., a_{k+r}\}$. This fact is used in the proof of lemma 2.

**Lemma 2**

Let $\{a_1, a_2, ...a_m\}$ be a sequence. For nonnegative integer $r$, consider a subsequence $\{a_k, a_{k+1}, ..., a_{k+r}\}$ such that $1 + r \leq k \leq k + r \leq m - r$ (i.e. $1 + r \leq k \leq m - 2r$). Let $b$ be a non-negative integer. Then, at most $b$ points within the subsequence $\{a_k, a_{k+1}, ..., a_{k+r}\}$ are $r$-locally top-$b$.

This lemma is represented graphically by Fig S2A.

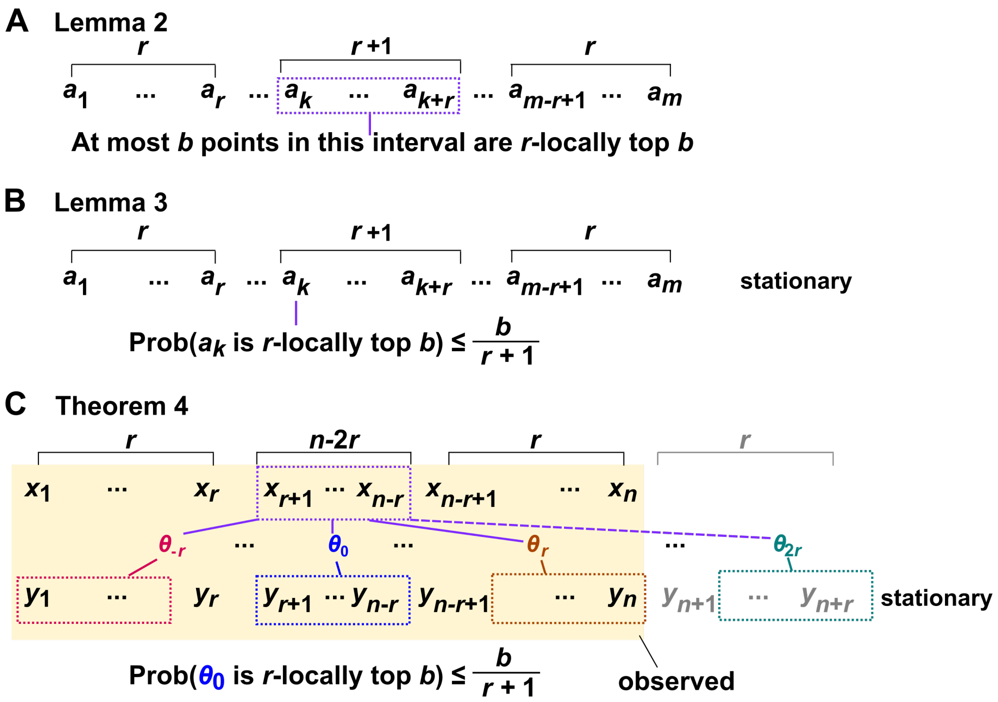

Figure S2: Illustration of elements of the proof.

**Proof:** We prove the lemma by contradiction. Suppose that at least $b{+}1$ points within $\{a_k, a_{k+1}, ..., a_{k+r}\}$ are $r$-locally top-$b$. Let $a_s$ be the smallest of these $b+1$ points (possibly tied for smallest). Then, $\{a_k, a_{k+1}, ..., a_{k+r}\}$ is within the $r$-neighborhood of $a_s$. (In detail, $s - r \leq k \leq k + r \leq s + r$ since $k \leq s \leq k + r$). Since there are at least $b+1$ points greater than or equal to $a_s$ in $a_s$'s $r$-neighborhood, $a_s$ is not $r$-locally top-$b$. Thus, we have a contradiction, which completes the proof. We required $1{+}r \leq k \leq m{-}2r$ to ensure that the $r$-neighborhoods of $\{a_k, a_{k+1}, ..., a_{k+r}\}$ are properly defined.

**Lemma 3**

Let $\{a_1, a_2, ...a_m\}$ be a stationary sequence of random variables. Consider an element $a_k$ such that $1 + r \leq k \leq m - 2r$ for some nonnegative integer $r$. Then, for some nonnegative integer $b$, the probability that $a_k$ is $r$-locally top-$b$ has an upper bound of $b/(r + 1)$.

This lemma is represented graphically by Fig S2B.

**Proof:** Define the variable $T_i(r, b)$ to indicate whether the $i$th point is $r$-locally top-$b$. Specifically, $T_i(r, b) = 1$ if $a_i$ is $r$-locally top-$b$ and $T_i(r, b) = 0$ otherwise. Since $1 + r \leq k \leq m - 2r$, lemma 2 says that at most $b$ points within $\{a_k, a_{k+1}, ..., a_{k+r}\}$ are $r$-locally top-$b$. Written in terms of $T_i(r, b)$, this condition is:

$$b \geq \sum_{i=k}^{k+r} T_i(r, b).$$

Since $\{a_1, a_2, ...a_m\}$ is stationary, $\{T_k(r, b), T_{k+1}(r, b), ..., T_{k+r}(r, b)\}$ must also be stationary. (This fact is intuitive, but a rigorous proof is given by lemma 13.) Denote by $E[\cdot]$ the expected value of a random variable (i.e. the mean). Since the mean of a stationary sequence is independent of time, we have:

$$b = E[b] \geq E\left[\sum_{i=k}^{k+r} T_i(r, b)\right] = \sum_{i=k}^{k+r} E\left[T_i(r, b)\right] = (r + 1)E\left[T_k(r, b)\right].$$

So $E\left[T_k(r, b)\right] \leq b/(r + 1)$. But $E\left[T_k(r, b)\right]$ is the probability that $a_k$ is $r$-locally top-$b$, so the proof is complete.

We are now ready to turn our attention to the theorem which justifies the TTS test. Since we will need to frequently mention specific sequences, we will make use of the abbreviation $\{a_1, a_2 ..., a_n\} = \{a_t\}_1^n$.

The statement of the following theorem parallels the TTS procedure closely, but with two main differences. The first difference is that we specify that the $x_t$ and $y_t$ terms are in $\mathbb{R}^l$ and $\mathbb{R}^k$. Essentially, this just means that the time series are allowed to be multivariate and are not required to have the same dimension. In the simple case where we are dealing with two univariate time series, we can just set both $l$ and $k$ to 1. Alternatively, it is possible to correlate time series with different dimensions (i.e. $l \neq k$) using correlation functions based on distances or predictions ([110, 6, 12]). An example where this may be useful is in Fig S12, where a series of one-dimensional $x_t$ terms is correlated to a two-dimensional time series whose terms are $v_t = (z_t, z_{t-1})$. In this example, a correlation is high if pairs of points that are nearby in $v_t$ space have similar $x_t$ values.

The second difference between the theorem below and the TTS procedure in the main text is that in the theorem we require the existence of $\{y_t\}_1^{n+r}$, despite only having data from $\{y_t\}_1^n$. This is a theoretical requirement. We do not need to obtain data from $\{y_{n+1}, ..., y_{n+r}\}$, but we must assume that the subsequence $\{y_{n+1}, ..., y_{n+r}\}$ exists in order to apply lemma 2.

**Theorem 4** *Validity of the truncated time-shift surrogate test*

Let $n$ ("the observed data length") and $r$ ("the truncation radius") be integers such that $0 < 2r < n$. Let $\{x_t\}_1^n$ be a (possibly nonstationary) stochastic sequence where each term $x_t$ is in $\mathbb{R}^l$. Let $\{y_t\}_1^{n+r}$ be a stationary stochastic sequence where each term $y_t$ is in $\mathbb{R}^k$. Let the two sequences be independent of each other. Let $\Theta$ (the "correlation function") be a real-valued function maps a pair of sequences (the "truncated $x$ series" $\{x_{1+r}, ..., x_{n-r}\}$ and the "shifted truncated $y$ series" $\{y_{1+r+\delta}, ..., y_{n-r+\delta}\}$) to a real number. Moreover, let $\Theta$ be $(\mathbb{R}^{(l+k)\times(n-2r)}, \mathcal{B}_{(l+k)\times(n-2r)})$-measurable[1]. Define the "shifted correlations" $\theta_\delta$ as

$$\theta_\delta = \Theta(\{x_{1+r}, ..., x_{n-r}\}, \{y_{1+r+\delta}, ..., y_{n-r+\delta}\}).$$

Let $B$ be the number of terms in the sequence

$$\theta_{-r}, \theta_{-r+1}, ..., \theta_r$$

that are greater than or equal to $\theta_0$. Then

$$P(B \leq b) \leq \frac{b}{r+1}$$

for any non-negative integer $b$.

This theorem is represented graphically by Fig S2C.

**Proof:** Since $\{x_t\}_1^n$ and $\{y_t\}_1^{n+r}$ are independent, and $\{y_t\}_1^{n+r}$ is stationary, the sequence

$$\theta_{-r}, \theta_{-r+1}, ..., \theta_{2r}$$

is also stationary (by theorem 12). Since $\theta_0$ is a member of a stationary sequence and is flanked by $r$ values to its left and $2r$ values to its right, we may apply lemma 3 to obtain:

$$\frac{b}{r+1} \geq P(\theta_0 \text{ is } r\text{−locally top−}b) = P(B \leq b)$$

[1] Although we have tried to avoid potentially difficult concepts in this proof, measurability does appear here as a necessary condition. We give a brief explanation here and a formal definition in section 1.2. Here, the phrase "$\Theta$ is a $(\mathbb{R}^{(l+k)\times(n-2r)}, \mathcal{B}_{(l+k)\times(n-2r)})$-measurable function" can be translated as: "for any pair of stochastic sequences $\{x_{1+r}, ..., x_{n-r}\}$ and $\{y_{1+r+\delta}, ..., y_{n-r+\delta}\}$ where each term $x_t$ is in $\mathbb{R}^l$ and each term $y_t$ is in $\mathbb{R}^k$, it is guaranteed that $\Theta(\{x_{1+r}, ..., x_{n-r}\}, \{y_{1+r+\delta}, ..., y_{n-r+\delta}\})$ is a random variable with a well-defined cumulative distribution function". If $\Theta$ is not measurable, then we may not be able to define the probability distribution of $\Theta(\{x_{1+r}, ..., x_{n-r}\}, \{y_{1+r+\delta}, ..., y_{n-r+\delta}\})$, which is a serious problem for statistical analysis. Measurability is a theoretical requirement, not a practical one: Essentially any function we choose in a practical correlation analysis will be measurable unless we intentionally engineer it to be otherwise.

which completes the proof.

Why does theorem 4 justify the TTS test? Recall that the TTS procedure defines the $u$ statistic to be:

$$u = \frac{B}{r+1}$$

with $B$ being the number of correlations ($\theta_\delta$'s) that are greater than or equal to the unshifted correlation ($\theta_0$) as in theorem 4. In the main text, we claimed that we can use $u$ like a $p$-value. That is, for a significance level $\alpha$, we have $P(u \leq \alpha) \leq \alpha$. To see why this is true, choose $b = \text{floor}(\alpha(r+1))$, where floor($a$) rounds $a$ down to the nearest integer. Then, as long as the requirements of theorem 4 are met, we have:

$$\begin{aligned} P(u \leq \alpha) &= P\left(\frac{B}{r+1} \leq \alpha\right) \\ &= P(B \leq \alpha(r+1)) \\ &\leq P(B \leq \text{floor}(\alpha(r+1))) \\ &= P(B \leq b) \leq b/(r+1) \leq \alpha \end{aligned}$$

where we applied theorem 4 and the fact that $b \leq \alpha(r+1)$ in the last line.

#### 1.2 Background definitions and supporting proofs

Here we review relevant definitions and properties from probability theory. Note that our definition of stationarity departs from much of the literature in that we work with a notion of stationarity that applies to finite time series. The final two results of this section, theorem 12 and lemma 13, are auxiliary results that support the arguments made in Appendix 1.1. Definitions 5, 6, and 8 are illustrated graphically in Fig S3.

**Definition 5** *Probability spaces (chapter 2 of [111])*

A probability space is any triple $(\Omega, \mathcal{F}, P)$ where $\Omega$, $\mathcal{F}$, and $P$ have the following names and meet the following requirements:

- $\Omega$ is called the sample space and is a nonempty set.
- $\mathcal{F}$ is called the $\sigma$-algebra and is a set of subsets of $\Omega$. $\mathcal{F}$ contains $\Omega$ itself and the empty set $\emptyset$. $\mathcal{F}$ is closed under the formation of complements and countable unions, meaning that $\mathcal{F}$ contains the complements and countable unions of any elements of $\mathcal{F}$.
- $P$ is called the probability measure and is a mapping from $\mathcal{F}$ to $[0, 1]$. $P(\emptyset) = 0$, $P(\Omega) = 1$, and $P$ is countably additive.

In essence, $\mathcal{F}$ (also known as the "event space") is the set of events (each event being a set) to which we can assign a probability value.

We now introduce the concept of Borel field on $\mathbb{R}$, a special type of $\sigma$-algebra generated from intervals on the real number line. This concept will be used in the formal definition of a random variable.

**Definition 6** *The Borel fields on $\mathbb{R}$ and $\mathbb{R}^k$ (definitions 3.18 and 3.19 in [112]).*

The Borel field on $\mathbb{R}$, denoted $\mathcal{B}$, is the smallest set of sets that includes:

1. all intervals $\{b : -\infty < b \leq \alpha\}$ where $\alpha \in \mathbb{R}$;
2. the complement $B^c$ of any set $B$ in $\mathcal{B}$;
3. the countable union of any sequence $\{B_i\}$ in $\mathcal{B}$.

Elements of a Borel field are called Borel sets.

The Borel field on $\mathbb{R}^k$ (where $k < \infty$), denoted $\mathcal{B}_k$, is the smallest collection of sets that includes:

1. all intervals $\{b : -\infty < b \leq \alpha\}$ where $b \in \mathbb{R}^k$ and $\alpha \in \mathbb{R}^k$;
2. the complement $B^c$ of any set $B$ in $\mathcal{B}_k$;
3. the countable union of any sequence $\{B_i\}$ in $\mathcal{B}_k$.

Above, in the multivariate case, the "$\leq$" and "$<$" inequality symbols apply element-wise.

The Borel fields are also closed under intersections. This fact will be useful later.

**Lemma 7**

If $B_1, B_2, ..., B_n$ (where $n$ is a positive integer) is a sequence of sets in $\mathcal{B}_k$, then the intersection $\cap_{i=1}^{n} B_i$ is also in $\mathcal{B}_k$.

**Proof:** For two sets $A$ and $B$, $A^C$, $A \cup B$, and $A \cap B$ denote the operations of complementation, union, and intersection respectively. From De Morgan's laws we know that

$$(A \cap B)^C = A^C \cup B^C.$$

Equivalently,

$$A \cap B = (A^C \cup B^C)^C. \tag{5}$$

Thus, since $\mathcal{B}_k$ is closed under complementation and union, if sets $A$ and $B$ are in $\mathcal{B}_k$, then so is $A \cap B$. This is the special case for two sets. The lemma itself then follows from the inductive generalization of the two-set case. Let $B_1, B_2, ...$ be a sequence of sets in $\mathcal{B}_k$. Let $\mathcal{C}(n)$ be the condition $\cap_{i=1}^{n} B_i \in \mathcal{B}_k$. We will show by induction that $\mathcal{C}(n)$ holds for all $n = 1, 2, ....$ The base case of $\mathcal{C}(1)$ is immediately satisfied as $\cap_{i=1}^{1} B_i = B_1$ is already in $\mathcal{B}_k$. For the inductive step, assume that $\mathcal{C}(s-1)$ holds, meaning that $\cap_{i=1}^{s-1} B_i \in \mathcal{B}_k$. Then, since $B_s \in \mathcal{B}_k$, we have:

$$\begin{aligned} \cap_{i=1}^{s} B_i &= (\cap_{i=1}^{s-1} B_i) \cap B_s \\ &= ((\cap_{i=1}^{s-1} B_i)^C \cup B_s^C)^C \in \mathcal{B}_k \end{aligned}$$

where the second line applied Eq. 5. Thus $\mathcal{C}(s)$ holds, completing the inductive step. We then conclude by induction that $\mathcal{C}(n)$ holds for $n = 1, 2, ...$, which proves the lemma.

Note that the empty set $\emptyset$ is in $\mathcal{B}_k$ as $\emptyset$ is the intersection of any set and its complement.

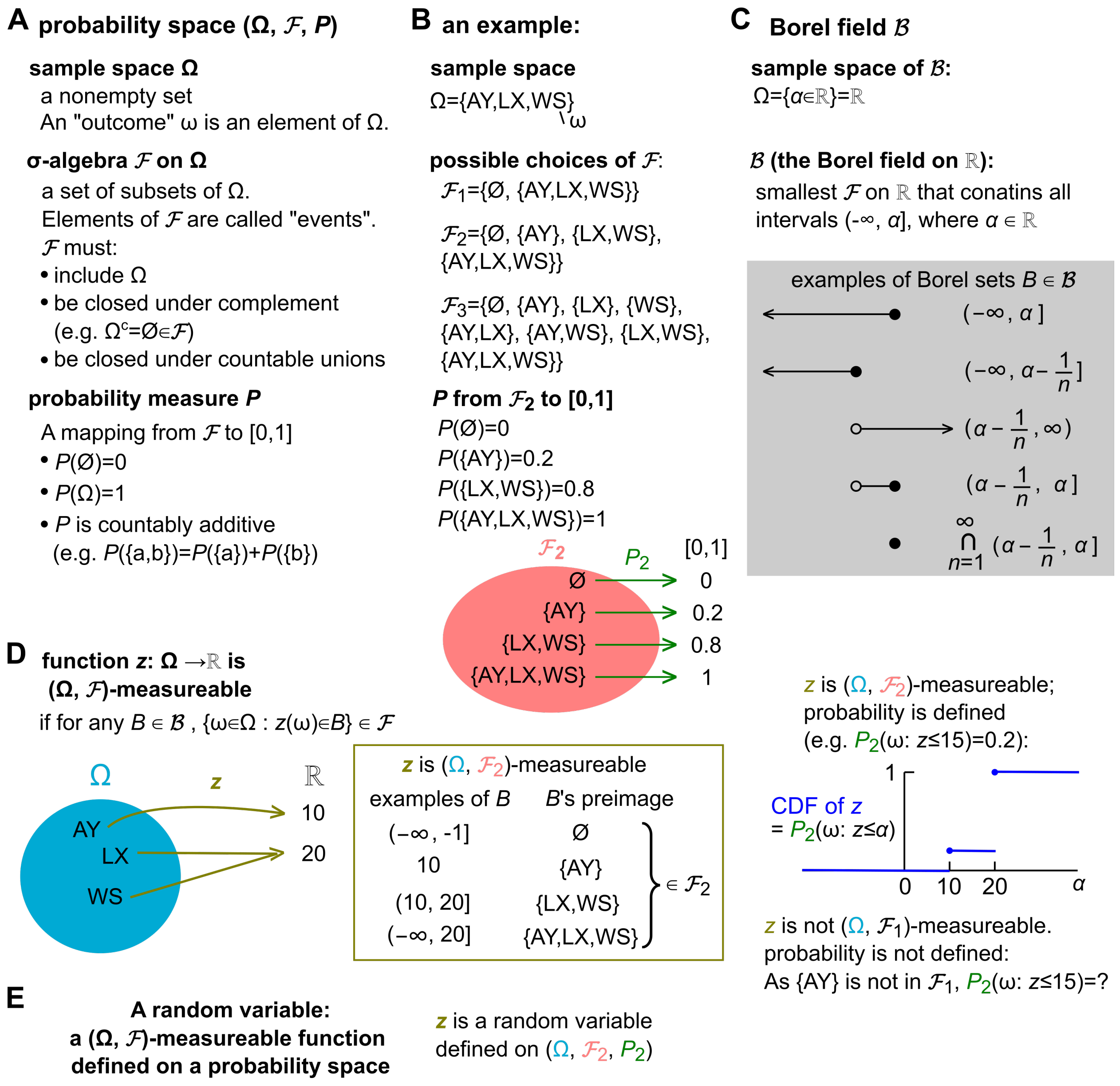

Figure S3: Some background concepts from probability theory. (**A**) A probability space includes a sample space $\Omega$, a $\sigma$-algebra $\mathcal{F}$, and a probability measure $P$. (**B**) An example of a probability space. Note that the "event" {LX,WS} occurs if either of the "outcomes" LX or WS occurs. We encourage readers to check these examples against the definitions in (A). (**C**) The Borel field $\mathcal{B}$ and some elements of $\mathcal{B}$ (which are called Borel sets). $\alpha$ is a real number, and $n$ is an integer. Rows 1 and 2 are elements of $\mathcal{B}$, as is seen immediately from $\mathcal{B}$'s definition. Row 3 is the complement of row 2. Row 4 is the intersection of rows 1 and 3. Row 5, the countable intersection of intervals $(\alpha - 1/n, \alpha]$ where $n$ ranges from 1 to $\infty$ (see, for instance, section 1.4 of [113]). This demonstrates that a real number is also an element of $\mathcal{B}$. (**D**) Measurement function: definition and illustration. Although not pictured, $z$ is also $(\Omega, \mathcal{F}_3)$-measurable. (**E**) Definition of a random variable.

The following concepts allow us to ensure that our models of probabilistic phenomena behave as they should (e.g. that their cumulative distribution functions [CDF]s exist; Fig S3D-E).

**Definition 8** *Measurable functions, random variables, and stochastic sequences*

- A real-valued function $z : \Omega \to \mathbb{R}$ is called measurable on $(\Omega, \mathcal{F})$, or $(\Omega, \mathcal{F})$-measurable, if for any set

$B$ in $\mathcal{B}$,

$$\{\omega \in \Omega : z(\omega) \in B\} \in \mathcal{F}.$$

In other words, for any $B$ in $\mathcal{B}$, its preimage (i.e. the set of all those $\omega$ in the sample space whose $z(\omega)$ value is in $B$) is an element of $\mathcal{F}$. The above definition of measurable functions is useful for proving results about functions already known to be measurable. However, another definition is equivalent (see page 29 of [111] or proposition 2.1.9 of [114]) and particularly useful for showing that a function is in fact measurable: A real-valued function $z : \Omega \to \mathbb{R}$ is measurable if for all $\alpha \in \mathbb{R}$,

$$\{\omega \in \Omega : z(\omega) \in (-\infty, \alpha]\} \in \mathcal{F}.$$

- A random variable is a $(\Omega, \mathcal{F})$-measurable real function defined on a particular probability space.
- A stochastic sequence is a family of random variables $\{z_1, z_2, ..., z_n\} = \{z_t\}_{t=1}^n$ where all such random variables are defined on the same probability space (i.e. they all share the same $\Omega$ and $\mathcal{F}$). Since in this work, we are dealing with finite-length stochastic sequences, we can alternatively think of a stochastic sequence as a vector-valued random variable.

**Definition 9** *Stationary stochastic sequences*

A stochastic sequence $\{x_1, ..., x_n\}$ is stationary if for all triples $(i, j, \tau)$ such that $1 \leq i \leq i + \tau \leq n$ and $1 \leq j \leq j+\tau \leq n$, the joint distribution of $\{x_i, ..., x_{i+\tau}\}$ is the same as the joint distribution of $\{x_j, ..., x_{j+\tau}\}$.

**Theorem 10** *Applying a measurable function to a stationary sequence produces a stationary sequence*

Let $\{x_1, ..., x_n\}$ be a stationary sequence of real-valued random variables. If $y_t = f(x_t, ..., x_{t+s})$ (where $1 \leq t \leq t + s \leq n$) for some $(\mathbb{R}^{s+1}, \mathcal{B}_{s+1})$-measurable function $f$, then $\{y_1, ..., y_{n-s}\}$ is stationary.

**Proof:** Let $B_0, B_1, ..., B_\tau$ be Borel sets in $\mathcal{B}$. We want to show that the joint distribution of $y$, $P\left((y_t \in B_0), (y_{t+1} \in B_1), ..., (y_{t+\tau} \in B_\tau)\right)$, is independent of time $t$ (i.e. identical between $t = i$ and $t = j$).

$$\begin{aligned} P\left((y_i \in B_0), (y_{i+1} \in B_1), ..., (y_{i+\tau} \in B_\tau)\right) =& P\left(\left((x_i, ..., x_{i+s}) \in f^{-1}B_0\right), ..., \left((x_{i+\tau}, ..., x_{i+\tau+s}) \in f^{-1}B_\tau\right)\right) \\ =& P\left(\left((x_j, ..., x_{j+s}) \in f^{-1}B_0\right), ..., \left((x_{j+\tau}, ..., x_{j+\tau+s}) \in f^{-1}B_\tau\right)\right) \\ =& P\left((y_j \in B_0), ..., (y_{j+\tau} \in B_\tau)\right) \end{aligned}$$

The second line follows from the stationarity of $\{x_1, ..., x_n\}$. Note that $f^{-1}(\cdot)$ denotes the preimage of $f$, not the inverse of $f$. The indexing terms $i, j, \tau, s$ are defined to prevent subsequences from falling off the edge (i.e. $1 \leq i \leq i + \tau \leq n - s$ and $1 \leq j \leq j + \tau \leq n - s$).

The last two results of this section are not background theory. Rather, they are auxiliary results that are necessary to fully detail the arguments in Appendix 1.1. Specifically, theorem 12 is used in the proof of theorem 4. Lemma 13 is used in the proof of lemma 3. In other words, we do not attempt to motivate the statements given below, and simply provide them to be used as a reference for readers working through the proofs of theorem 4 and lemma 13.

**Theorem 11** *Functions of independent random varaibles are independent (Proposition 3.2.3 of [111])*

Let $x \in \mathbb{R}^{m\times n}$ and $y \in \mathbb{R}^{m\times n}$ be random variables (or vectors of random variables, or matrices of random variables) such that $x$ and $y$ are independent. Let $f : \mathbb{R}^{m\times n} \to \mathbb{R}^{m\times n}$ and $g : \mathbb{R}^{m\times n} \to \mathbb{R}^{m\times n}$ be functions that are measurable on $(\mathbb{R}^{m\times n}, \mathcal{B}_{m\times n})$. Then, $f(x)$ and $g(y)$ are independent.

**Proof:** Let $S_x$ and $S_y$ be Borel sets in $\mathcal{B}_{m\times n}$. Suppose that $x$ and $y$ are independent. Then, considering the joint distribution of $f(x)$ and $g(y)$, we have:

$$\begin{aligned}P(f(x)\in S_x, g(y)\in S_y) &= P(x\in f^{-1}S_x, y\in g^{-1}S_y)\\ &= P(x\in f^{-1}S_x)P(y\in g^{-1}S_y)\\ &= P(f(x)\in S_x)P(g(y)\in S_y)\end{aligned}$$

Since the joint distribution of $f(x)$ and $g(y)$ is equivalent to the product of the marginal distributions, $f(x)$ and $g(y)$ are independent. This proof is given in [111] for the univariate case, but we repeat it here to stress that the same logic applies in the multivariate case.

**Theorem 12**

Let $\{x_1, ..., x_m\}$ and $\{y_1, ..., y_n\}$ be independent stochastic sequences, where $\{y_1, ..., y_n\}$ is stationary. The two sequences may be multivariate (i.e. $x_t \in \mathbb{R}^k$ and $y_t \in \mathbb{R}^r$). Construct a sequence $\{z_1, ..., z_{n-s}\}$ such that $z_t = f(\{x_1, ..., x_m\}, \{y_t, ..., y_{t+s}\})$ where $1 \le t \le t+s \le n$ and $f$ is $(\mathbb{R}^{mk+r(s+1)}, \mathcal{B}_{mk+r(s+1)})$-measurable. That is, each term $z_t$ is a function of the entire $x$ sequence and of a sliding window of size $s+1$ within the $y$ sequence (e.g. a correlation function of two time series). Then $\{z_1, ..., z_{n-s}\}$ is stationary.

**Proof:** The proof is similar in spirit to that of theorem 10. The difference is that here, $f$ is a function of two series, and thus the preimage of $f$ will consist of two sets. For a Borel set $B$ in $\mathcal{B}$, let its preimage $f^{-1}B = A$, where $A$'s $x$ component looks like $\{x_1, ..., x_m\}$ (i.e. $A^x \subseteq \mathbb{R}^{km}$ ) and $A$'s $y$ component looks like $\{y_1, ..., y_n\}$ (i.e. $A^y \subseteq \mathbb{R}^{r(s+1)}$).

As before, we proceed by showing that the joint distribution of $\{z_i, ..., z_{i+\tau}\}$ is the same as the joint distribution of $\{z_j, ..., z_{j+\tau}\}$ (where $i$ and $j$ may differ). Let $B_0, B_1, ..., B_\tau$ be Borel sets in $\mathcal{B}$.

$$\begin{aligned}P\left((z_i\in B_0),...,(z_{i+\tau}\in B_\tau)\right) &= P\left(\left(\left(\{x_t\}_1^m,\{y_t\}_i^{i+s}\right)\in f^{-1}B_0\right),...,\left(\left(\{x_t\}_1^m,\{y_t\}_{i+\tau}^{i+\tau+s}\right)\in f^{-1}B_\tau\right)\right)\\ &= P\left(\left(\left(\{x_t\}_1^m,\{y_t\}_i^{i+s}\right)\in A_0\right),...,\left(\left(\{x_t\}_1^m,\{y_t\}_{i+\tau}^{i+\tau+s}\right)\in A_\tau\right)\right)\\ &= P\left(\left(\{x_t\}_1^m\in A_0^x,\{y_t\}_i^{i+s}\in A_0^y\right),...,\left(\{x_t\}_1^m\in A_\tau^x,\{y_t\}_{i+\tau}^{i+\tau+s}\in A_\tau^y\right)\right)\end{aligned}$$

Since $x$ and $y$ series are independent, the above equation can be rewritten as

$$P\left((z_i\in B_0),...,(z_{i+\tau}\in B_\tau)\right) = P\left((\{x_t\}_1^m\in A_0^x),...,(\{x_t\}_1^m\in A_\tau^x)\right)P\left(\left(\{y_t\}_i^{i+s}\in A_0^y\right),...,\left(\{y_t\}_{i+\tau}^{i+\tau+s}\in A_\tau^y\right)\right)$$

Since the $y$ series is stationary, we have:

$$\begin{aligned}P\left((z_i\in B_0),...,(z_{i+\tau}\in B_\tau)\right) &= P\left((\{x_t\}_1^m\in A_0^x),...,(\{x_t\}_1^m\in A_\tau^x)\right)P\left(\left(\{y_t\}_j^{j+s}\in A_0^y\right),...,\left(\{y_t\}_{j+\tau}^{j+\tau+s}\in A_\tau^y\right)\right)\\ &= P\left((z_j\in B_0),...,(z_{j+\tau}\in B_\tau)\right)\end{aligned}$$

Thus, $z$ is stationary.

**Lemma 13**

Let $\{a_1, a_2, ...a_m\}$ be a stationary sequence of random variables. Let $b$ and $r$ be non-negative integers. Consider a subsequence $\{a_{1+r}, ..., a_{m-r}\}$ so that the $r$-neighborhood of each element is valid. For $1+r \le k \le m-r$, define $T_k(r,b) = \psi(a_{k-r}, ..., a_{k+r})$ to indicate whether $a_k$ is $r$-locally top-$b$. Specifically, $T_k(r,b) = 1$ if $a_k$ is $r$-locally top-$b$ and $T_k(r,b) = 0$ otherwise. Then, $\{T_{1+r}(r,b), ..., T_{m-r}(r,b)\}$ is a stationary sequence of random variables.

**Proof:** According to Theorem 10, we can prove this theorem by showing that the function $\psi$ is measurable on $(\mathbb{R}^{2r+1}, \mathcal{B}_{2r+1})$. To show that $\psi$ is measurable on $(\mathbb{R}^{2r+1}, \mathcal{B}_{2r+1})$, it is sufficient (see definition 8) to show that for all $(-\infty, \alpha]$ where $\alpha \in \mathbb{R}$,

$$\{\overrightarrow{a_k} \in \mathbb{R}^{2r+1} : \psi(\overrightarrow{a_k}) \in (-\infty, \alpha]\} \in \mathcal{B}_{2r+1} \tag{6}$$

where $\overrightarrow{a_k} = \{a_{k-r}, ..., a_{k+r}\}$. We will also refer to $\{\overrightarrow{a_k} : \psi(\overrightarrow{a_k}) \in (-\infty, \alpha]\}$ as "the preimage" of $(-\infty, \alpha]$ under $\psi$.

There are three cases to consider. First, if $\alpha < 0$, then the preimage is simply the empty set $\emptyset$, since $T_k(r, b)$ is never less than 0. $\emptyset$ is an element of $\mathcal{B}_{2r+1}$, so the condition of Eq. 6 is satisfied. Second, if $\alpha \geq 1$, since $T_k(r, b) \leq 1$, Eq. 6 is again satisfied as the preimage then becomes $\mathbb{R}^{2r+1}$, which is an element of $\mathcal{B}_{2r+1}$.

Lastly, if $0 \leq \alpha < 1$, then the preimage is the set of all $\overrightarrow{a_k}$ where more than $b$ elements are greater than or equal to $a_k$. But since one of these elements, namely $a_k$ itself, is always equal to $a_k$, we can rephrase the above statement to be more useful for the subsequent arguments: *The preimage is the set of all $\overrightarrow{a_k}$ where at least $b$ elements (other than $a_k$ itself) are greater than or equal to $a_k$.* A trivial edge case is where $b = 0$, in which case the preimage is $\mathbb{R}^{2r+1} \in \mathcal{B}_{2r+1}$. When $b > 0$, we will need a more careful strategy to show that the preimage is in $\mathcal{B}_{2r+1}$.

Our strategy will be similar to the one used in Fig S3C (grey box) where we showed that a set consisting of a single real number is a member of $\mathcal{B}$. Analogously to that argument, here we will find some "starting sets" that are known to be in $\mathcal{B}_{2r+1}$ and show that one can arrive at the preimage by taking countable unions and intersections of these starting sets.

To build our intuition, let's first consider the specific setting where $r = 2, b = 3$. The preimage is then the set of all $\overrightarrow{a_k} = (a_{k-2}, a_{k-1}, a_k, a_{k+1}, a_{k+2})$ where at least 3 elements (other than $a_k$ itself) are greater than or equal to $a_k$. We begin with "starting sets" $\mathcal{S}_i = \{\overrightarrow{a_k} : a_{k+i} \geq a_k\}$, where $i = -2, -1, 1, 2$. For example, $\mathcal{S}_1$ is the set of all $\overrightarrow{a_k}$s wherein $a_{k+1} \geq a_k$. To obtain some geometric intuition for this set, note that if $\overrightarrow{a_k}$ had been simply $(a_k, a_{k+1})$, then $a_{k+1} \geq a_k$ would correspond to the half plane on and above the line of identity $a_{k+1} = a_k$. $\mathcal{S}_1$ is in $\mathcal{B}_5$, because its complement (the set of all $\overrightarrow{a_k}$s wherein $a_{k+1} < a_k$) is an open set. Note that a set $O$ is called "open" if for every point $x$ in $O$, there is some positive "neighborhood radius" $\epsilon$ such that all of $x$'s "neighbors" (i.e. points less than $\epsilon$ away from $x$) are also in $O$ (e.g. page 385 of [114]). Since any open set is a Borel set (e.g. proposition 1.1.5 of [114]), $\mathcal{S}_1$ is in $\mathcal{B}_5$. By the same logic, all four $\mathcal{S}_i$s are in $\mathcal{B}_5$. Fig S4 shows how we can arrive at the preimage by taking unions and intersections of $\mathcal{S}_{-2}$, $\mathcal{S}_{-1}$, $\mathcal{S}_1$, and $\mathcal{S}_2$. The intersection $\mathcal{S}_{-2} \cap \mathcal{S}_{-1} \cap \mathcal{S}_2$, which is outlined in red and resembles a petal, is the set of all $\overrightarrow{a_k}$ where $a_{k-2}$, $a_{k-1}$, and $a_{k+2}$ are all $\geq a_k$. Clearly this is a subset of the preimage because it is one way in which at least 3 elements (other than $a_k$) are $\geq a_k$. There are three other such triple intersections, which are the three other overlapping "petals" in Fig S4. The preimage itself, shown as the grey shaded area, is the union of these four petals.

The same reasoning holds for other choices of $b$ and $r$. In general, the starting sets are $\mathcal{S}_i = \{\overrightarrow{a_k} : a_{k+i} \geq a_k\}$, where $-r \leq i \leq r$ and $i \neq 0$. In words, $\mathcal{S}_i$ is the set of all $\overrightarrow{a_k}$s wherein $a_{k+i} \geq a_k$. By the same reasoning as above, these starting sets $\mathcal{S}_i$ are all elements of $\mathcal{B}_{2r+1}$. Any intersection of $b$ starting sets is a situation where $a_k$ is not $r$-locally top-$b$, and the preimage is given by the union of all such intersections. Since the preimage can be constructed from starting sets in $\mathcal{B}_{2r+1}$ by taking countable unions and intersections, the preimage must be in $\mathcal{B}_{2r+1}$ for this (final) case.

Overall, for all $\alpha \in \mathbb{R}$, the condition of Eq. 6 is satisfied, which in turn establishes that

$$\{T_k(r, b), T_{k+1}(r, b), ..., T_{k+r}(r, b)\}$$

is a stationary sequence of random variables, as required.

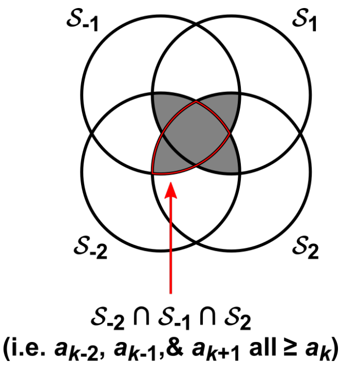

Figure S4: An example of how we can obtain the preimage by taking intersections and unions of starting sets $\mathcal{S}_i$. Here, $r = 2$ and $b = 3$, so the preimage (shaded in grey) is the set of all $\overrightarrow{a_k} = (a_{k-2}, a_{k-1}, a_k, a_{k+1}, a_{k+2})$ where at least 3 elements (other than $a_k$ itself) are greater than or equal to $a_k$.

### 2 Certain variants of the naive TTS test may be miscalibrated by more than twofold

In the main text we pointed out that the naive TTS test (Eq. 2) will never be miscalibrated by more than a factor of 2 as long as the time series used to produce surrogate data is stationary. In practice, the naive TTS test is not typically implemented exactly according to the procedure we describe, but instead several related variants are used [36, 38, 115, 39, 32]. Are variants of the naive TTS test also guaranteed to be miscalibrated by no more than twofold when applied to stationary time series? Here we describe two possible variants of the TTS test, and show that they can be miscalibrated by well over twofold. We do use rather extreme cases as examples in this section. Our purpose here is only to show that it is possible for these variants to fail catastrophically, in contrast to the naive TTS procedure described in the main text, which will not be miscalibrated by more than a factor of 2 (for stationary data). A study of how likely these failure modes are to occur in realistic systems is beyond the scope of this section.

One way in which applied works modify the time-shifting approach is by the use of what we call an "exclusion radius" [36, 38, 39, 32]. The idea is that the shifted time series that comprise the null model must all be shifted by more than some user-defined amount. We now describe this procedure in the context of the naive TTS test. As in Fig 1, the procedure begins with two time series $\{x_t\}_{t=1}^n$ and $\{y_t\}_{t=1}^n$, choose a flanking radius $r$, and define the truncated sequences $x^{trunc} = \{x_{1+r}, ..., x_{n-r}\}$ and $y^{trunc}(\delta) = \{y_{1+r+\delta}, ..., y_{n-r+\delta}\}$. Also as in Fig 1, we define shifted correlations $\theta_\delta = \Theta(x^{trunc}, y^{trunc}(\delta))$. At this point we break with the standard recipe and choose an exclusion radius $r_{ex}$, which is a number that must be less than $r$. Rather than using all shifted $y^{trunc}$ sequences to compute the naive $p$-value, we will only use those with a shift larger than $r_{ex}$. To express this idea as an equation, define $B_-(r_{ex})$ to be the number of shifted correlations within $\{\theta_{-r}, \theta_{-r+1}, ..., \theta_{-r_{ex}-1}\}$ that are greater than or equal to $\theta_0$. Similarly define $B_+(r_{ex})$ to be the number of shifted correlations within $\{\theta_{r_{ex}+1}, \theta_{r_{ex}+2}, ..., \theta_r\}$ that are greater than or equal to $\theta_0$. Finally the empirical $p$-value (corresponding as always to the null hypothesis that the two time series are independent) is written as

$$p = \frac{B_-(r_{ex}) + B_+(r_{ex}) + 1}{2(r - r_{ex}) + 1}.$$

Note that in special case where $r_{ex} = 0$, this procedure reduces to the naive TTS test (Eq. 2).

We now give a simulation example of a pair of independent stationary systems $\{x_t\}$ and $\{y_t\}$ for which this test (with $r_{ex} > 0$) is severely miscalibrated. The $\{x_t\}$ series is generated according to a periodic process afflicted with periodically varying measurement noise:

$$x_t = (1 - \lambda_t)b_t + \lambda_t \epsilon_t$$

We can think of $b_t$ as a signal with a measurement strength of $(1 - \lambda_t)$ and we can think of $\epsilon_t$ as noise with strength of $\lambda_t$. The the noise terms $\epsilon_t$ are independently and identically distributed continuous uniform random variables between 0 and 1. The noise strength $\lambda_t$ is given by:

$$\lambda_t = \begin{cases} 0.8 & \text{if } a_t \geq 450 \\ 0.5 & \text{if } a_t < 450 \end{cases}$$

$$a_t = \text{mod}(t + \phi, 950)$$

where $\text{mod}(\alpha, \beta)$ is the modulo operation, which is the remainder obtained from dividing $\alpha$ by $\beta$. The phase term $\phi$ is chosen from $\{0, 1, ..., 949\}$ with equal chance. The initial signal value $b_1$ is chosen to be 0 or 1 with equal chance and all subsequent $b_t$ values are given by $b_{t+1} = 1 - b_t$ (i.e. they alternate between 0 and 1). $\phi$, $b_1$, and the $\epsilon_t$ terms are all independent. The $\{y_t\}$ series is an independent realization of the same process as the $\{x_t\}$ series (i.e. with independent choices of $\phi$, $b_1$, and $\epsilon_t$). Sample realizations are shown in Fig S5A.

To see that $\{x_t\}$ is stationary, note that $\{a_t\}$ is stationary because $\{a_t\}$ is a periodic process whose phase is chosen uniformly from throughout its period. Similarly, $\{b_t\}$ is stationary for the same reason. Next, $\{\lambda_t\}$ is stationary by theorem 10 since $\{\lambda_t\}$ is a static function of $\{a_t\}$. Lastly, since all three of $\{b_t\}$, $\{\lambda_t\}$ and $\{\epsilon_t\}$ are stationary and independent of one another, $\{x_t\}$ must be stationary.

We tested whether the two time series were dependent using a pair of length-2000 time series, a flanking radius of $r = 500$, and an exclusion radius of $r_{ex} = 450$. We used the sample Pearson correlation coefficient as the correlation statistic. In 43% of 1000 trials, dependence was detected at the 0.05 significance level (Fig S5B).

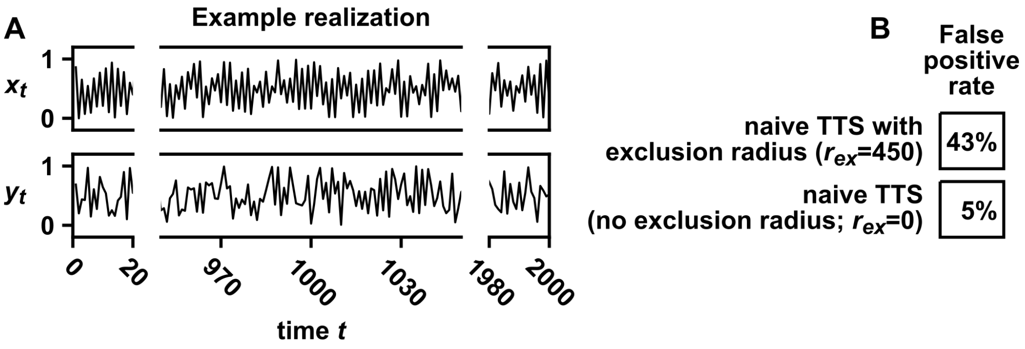

Figure S5: The exclusion radius procedure can cause the naive TTS test to be miscalibrated by more than twofold. (A) Sample dynamics of the benchmark system. See text for details. (B) False positive rates of the naive TTS test with or without an exclusion radius. For both tests, the truncation radius $r$ was set to 500, the correlation statistic was the absolute value of the Pearson correlation coefficient, and significance was evaluated at the 0.05 level.

A second possible variant of the naive TTS test concerns possible delayed coupling. If two time series are coupled but with a delay, then the time shift approach may fail to detect a dependence relationship because $\{x_t\}$ may be more strongly correlated to shifted $\{y_t\}$ series than the unshifted $\{y_t\}$ series. In the main text, our proposed solution is to pre-shift one of the two series (Fig 3). However, a seemingly natural alternative solution is the following: We define truncation radius $r$, $x^{trunc}$, $y^{trunc}(\delta)$, correlation function $\Theta$, and shifted correlation $\theta_\delta = \Theta(x^{trunc}, y^{trunc}(\delta))$ as before, but now we choose a new parameter $\Delta$ (the coupling delay we expect to be in the system). $\Delta$ is the value of $\delta$ at which we expect the correlation $\theta_\delta$ to be maximized. We then let $B_\Delta$ be the number of shifted correlations $\theta_\delta$ $(-r \leq \delta \leq r)$ that are as large as or larger than $\theta_\Delta$. We then write the empirical $p$-value as:

$$p = \frac{B_\Delta}{2r + 1}.$$

We call this the "naive TTS test with an expected coupling delay" and note that the naive TTS test is the special case of this test where $\Delta = 0$. Although we have not found this variant in the literature, we have been asked about it when presenting this work.

We now give a simulation example of a pair of independent stationary processes $\{x_t\}$ and $\{y_t\}$ for which this test is miscalibrated by more than twofold. Both are periodic process afflicted with periodically varying measurement noise. $\{x_t\}$ is given by:

$$x_t = (1 - \lambda_{x,t})b_{x,t} + \lambda_{x,t}\epsilon_{x,t}$$

$$t = 1, 2, ..., 400$$

where

$$\lambda_{x,t} = 0.1 + 0.1\left(\frac{\text{mod}(t + \phi_x, 3000)}{3000}\right)$$

$$b_{x,t} = 1 - b_{x,t-1}; b_{x,1}\text{chosen to be 0 or 1 with equal chance}$$

Next, $\{y_t\}$ is given by similar formulae, but where the equation for $\lambda_{y,t}$ has some differences in parameter choices:

$$y_t = (1 - \lambda_{y,t})b_{y,t} + \lambda_{y,t}\epsilon_{y,t}$$

$$\lambda_{y,t} = 0.3 + 0.5\left(\frac{\text{mod}(t + \phi_y, 3000)}{3000}\right)$$

$$b_{y,t} = 1 - b_{y,t-1}; b_{y,1}\text{chosen to be 0 or 1 with equal chance}$$

Here, $\phi_x$ and $\phi_y$ are chosen from among $\{0, 1, ..., 2999\}$ with equal chance. The $\epsilon_{x,t}$ and $\epsilon_{y,t}$ noise terms are independently and identically distributed continuous uniform random variables between 0 and 1. $\phi_x$, $\phi_y$, $b_{x,1}$, $b_{y,1}$, and the $\epsilon_{x,t}$ and $\epsilon_{y,t}$ terms are all independent. As with the process of Fig S6, both $\{x_t\}$ and $\{y_t\}$ are stationary here as well; this can be seen using a line of argument analgous to the one used for the process of Fig S6. Sample realizations are shown in Fig S6A.

We tested whether the two time series were dependent using a naive TTS test with a flanking radius of $r = 100$ and various choices for the expected coupling delay $\Delta$. We used the sample Pearson correlation coefficient as the correlation statistic. Fig S6B shows how the false positive rate varies as a function of the expected coupling delay. In this example, strongly negative values of $\Delta$ result in a false positive rate as high as 30%, even though detections were made at the 0.05 significance level.

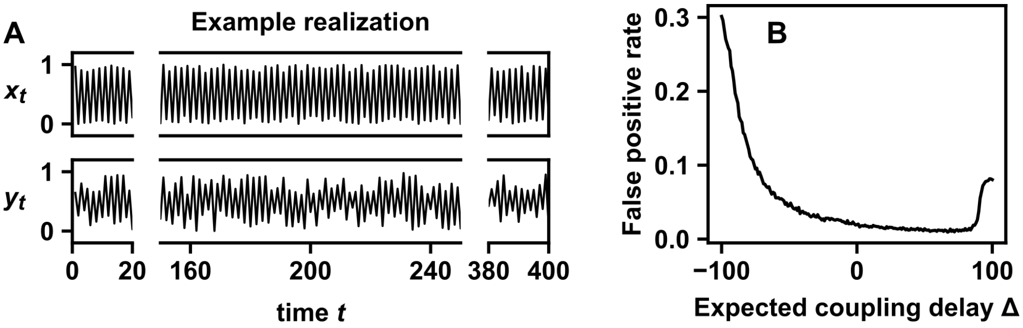

Figure S6: The naive TTS test with an expected coupling delay can be miscalibrated by more than twofold. (A) Sample dynamics of the benchmark system. See text for simulation details. (B) False positive rate of the naive TTS test as a function of the expected coupling delay $\Delta$. The truncation radius $r$ was set to 100 and the correlation statistic was the absolute value of the Pearson correlation coefficient. We varied the expected coupling delay $\Delta$ from $-100$ to 100. Detections were made at the 0.05 significance level. False positive rates were estimated from $10,000$ replicate trials.

### 3 Detailed methods and results for the simulation benchmark

Here we describe details for data-generating processes and hypothesis tests that we used in the benchmark study of false positive rates (Fig 2).

#### 3.1 Data-generating processes

For all data-generating processes except the sine wave with drifting measurement noise (system iii), the $x_t$ series and $y_t$ series were generated as independent replicates of the same process. Therefore, for all processes except those of system iii, we describe only the procedure to generate the $x_t$ series.

**Standard first-order autoregressive process (system i)**

$$t = 1, 2, ..., 400$$

$$x_{t+1} = 0.7x_t + \epsilon_t$$

where the $\epsilon_t$ terms are independent random variables with a standard normal distribution (i.e. with a mean of zero and variance of 1). The initial condition $x_1$ follows the stationary distribution of the process, which in this case is a normal distribution with a mean of zero and a standard deviation of $(1 - 0.7^2)^{-1/2}$ (see Eq. 20-6 in [43], for instance).

**Logistic map (system ii)**

$$t = 1, 2, ..., 400$$

$$x_{t+1} = 4x_t(1 - x_t)$$

where $x_1$ is chosen according to the Beta(0.5, 0.5) distribution. This system is stationary (see chapter 1 of [116], for instance).

**Sine wave with measurement noise whose strength varies as a sawtooth wave (system iii)**

$$t = 1, 2, ..., 400$$

$$x_t = \sin\left(\frac{2\pi(t + \phi_x)}{22}\right)$$

$$y_t = \sin\left(\frac{2\pi t + \phi_y}{22}\right) + a_t(\epsilon_t - 0.5)$$

where

$$a_t = \frac{\text{mod}(t + \phi_a, 2800)}{7000}$$

where mod$(\alpha, \beta)$ is the modulo operation, which is the remainder obtained from dividing $\alpha$ by $\beta$. The terms $\phi_x$, $\phi_y$, $\phi_a$ and $\{\epsilon_t\}_{t=1}^{400}$ are all independent random variables with the following distributions: $\phi_x$ and $\phi_y$ are chosen uniformly at random from $\{0, 1, ..., 21\}$. $\phi_a$ is chosen uniformly at random from $\{0, 1, ..., 2799\}$. The $\epsilon_t$ terms are independent random variables drawn from a Beta(1/3, 1/3) distribution. Note that the phase terms $\phi_x$, $\phi_y$, and $\phi_a$ are all time-invariant. More generally, we stress that throughout this section, all terms not indexed by $t$ are time-invariant.

$\{x_t\}$ is stationary since it is a periodic sequence whose initial value is chosen uniformly from among the points within a period. To see that $\{y_t\}$ is stationary, start with $\{a_t\}$. We know that $\{a_t\}$ is stationary because it is a periodic sequence whose initial value is chosen uniformly from among the points within a period. Since $\{\epsilon_t\}$ are iid (and thus stationary), and since any static function of two independent stationary sequences is stationary, the "$a_t(\epsilon_t - 0.5)$" term in the equation for $y_t$ is also stationary. The sinusoid term in the equation for $y_t$ is also stationary since it is a periodic sequence whose initial value is chosen uniformly from among the points within a period. Finally, $\{y_t\}$ must be stationary since it is the sum of two terms that are independent of each other and also individually stationary.

**Sine wave with detection threshold (system iv)**

This process describes a sine wave with a detection threshold (0.5 and additive noise). See Fig S8 for more details.

$$t = 1, 2, ..., 400$$

$$x_t = a_t + \epsilon_t$$

$$a_t = \max(\sin(\phi + \frac{2\pi t}{35}), 0.5)$$

where $\phi$ is a continuous uniform random variable drawn from between 0 and $2\pi$, and $\epsilon_t$ terms are independent normal random variables with mean of 0 and standard deviation of 0.05.

To see that $\{x_t\}$ is stationary, start with $\{a_t\}$. Note that since $\phi$ is a continuous uniform random variable drawn from between 0 and $2\pi$, it follows that $\sin(\phi)$ and $\sin(\phi + t)$ have the same distribution for any $t$ (due to the periodicity of the sin function). It then follows that $a_1, a_2, ..., a_n$ has the same distribution as $a_{1+t}, a_{2+t}, ..., a_{n+t}$ for any $t$. That is, $\{a_t\}$ is stationary. The $\{x_t\}$ process is stationary because it is a static function of two independent stationary processes ($\{a_t\}$ and $\{\epsilon_t\}$).

**Coin flips with additive noise and time-varying 'heads' probability (system v)**

$$t = 1, 2, ..., 400$$

$$x_t = b_t + \epsilon_t$$

where $\{\epsilon_t\}_{t=1}^{400}$ are independent random normal variables with mean of 0 and standard deviation of 0.15. The terms $b_t$ are Bernoulli random variables ('coin flips') with probability:

$$P(b_t = 0) = 1 - a_t$$
$$P(b_t = 1) = a_t$$

where $a_t$ is given by:

$$a_t = \frac{1}{2}\left(\frac{1}{2}\sin\left(\phi_1 + \frac{2\pi t}{75}\right) + \frac{1}{2}\right)^6 + \frac{1}{12}\left(\frac{1}{2}\sin\left(\phi_2 + \frac{2\pi t}{31\sqrt{2}}\right) + \frac{1}{2}\right)$$

and where $\phi_1$ and $\phi_2$ are independent continuous uniform random variables drawn from between 0 and $2\pi$.

To see that $\{x_t\}$ is stationary, note that both additive terms in $\{a_t\}$ are stationary because they are each periodic functions whose phase is uniformly distributed over the period, similar to the sine wave in system iv. Then, $\{a_t\}$ is itself stationary because it is a function of two independent stationary sequences. Then, $\{x_t\}$ is stationary because it is a function of $\{a_t\}$ and $\{\epsilon_t\}$, which are also two independent stationary sequences.

**Exponential growth with periodic extinction (system vi)** Consider a population with a size of $a_t$ that experiences constant immigration (50 each time step) together with exponential growth punctuated with periodic extinction (every 8 time units):

$$a_{t+1} = \begin{cases} a_t + 0.2a_t + 50 & \text{if } t \text{ is not a multiple of } 8 \\ 50 & \text{if } t \text{ is a multiple of } 8 \end{cases}$$

$$a_1 = 50.$$

Suppose that the population has already been in existence for a random amount of time when observations begin. In this case, the process during the observation window may be $b_1, ..., b_{400}$:

$$b_t = a_{t+\phi}$$

where $\phi$ is an integer drawn uniformly at random from among $\{0, 1, ..., 7\}$.

Additionally, only about 10 percent of the population is observed, via a binomial sampling process, which we approximate using a normal distribution:

$$x_t = 0.1b_t + (0.09b_t)^{1/2}\epsilon_t$$

where the $\epsilon_t$ terms are independent normal random variables with mean of 0 and variance of 1. As usual, $x_1, ..., x_{400}$ is the reported time series.

To see that $\{x_t\}$ is stationary, note that $\{b_t\}$ is stationary because it is a periodic process whose phase is chosen uniformly at random from among the period. Then, $\{x_t\}$ is stationary because it is a static function of $\{b_t\}$ and $\{\epsilon_t\}$, which are two independent stationary processes.

##### FitzHugh-Nagumo model with noise (system vii)

$$(v_{t+1} - v_t)/0.7 = v_t - v_t^3/3 - w_t + 1 + 0.2\epsilon_t$$
$$(w_{t+1} - w_t)/0.7 = 0.08(x_t + 0.7 - 0.8w_t)$$

where the $\epsilon_t$ terms are independent random variables with standard normal distribution. We started the simulation with the initial condition $(v_1, w_1) = (0, 0)$ and ran the system for 1000 time points to allow it to equilibrate, using points 1001 through 1400 for the statistical benchmark. The $x_t$ and $y_t$ series were independent realizations of $v_t$.

##### Chaotic Lotka-Volterra (system viii)

$$\frac{ds_i(t)}{dt} = 1.5r_i\left(s_i(t) - \sum_{j=1}^{4} a_{ij}s_i(t)s_j(t)\right),\ i = 1, 2, 3, 4$$

$$r = \begin{bmatrix} 1 \\ 0.72 \\ 1.53 \\ 1.27 \end{bmatrix}$$

$$a = \begin{bmatrix} 1 & 1.09 & 1.52 & 0 \\ 0 & 1 & 0.44 & 1.36 \\ 2.33 & 0 & 1 & 0.47 \\ 1.21 & 0.51 & 0.35 & 1 \end{bmatrix}$$

Unlike the other systems which are discrete-time and mostly stochastic, this is a system of differential equations. These parameter choices are taken from [48]. Although this is a continuous-time system, statistical analysis was performed on discrete time points. We initialized the system by independently choosing $s_1(1)$, $s_2(1)$, $s_3(1)$, and $s_4(1)$ from a continuous uniform distribution between 0.1 and 0.5. We then numerically integrated the system for 2000 time units to allow it to equilibrate and finally used $t = 2001, 2002, ..., 2400$ for the statistical benchmark. The $x_t$ and $y_t$ series were independent realizations of $s_4(t)$.

##### Random walk (system ix)

$$t = 1, 2, ..., 400$$

$$x_{t+1} = x_t + \epsilon_t$$

where the $\epsilon_t$ terms are independent random variables with a standard normal distribution. The initial condition $x_1$ was set to 0.

##### First-order autoregressive process with trend (system x)

This system was generated by adding $t/60$ to the stationary first-order autoregressive process. Specifically,

$$t = 1, 2, ..., 400$$

$$x_t = a_t + t/60$$

where

$$a_{t+1} = 0.7a_t + \epsilon_t$$

Here the $\epsilon_t$ terms are independent random variables with a standard normal distribution. The initial condition $a_1$ is given by a normal distribution with a mean of zero and a standard deviation of $(1 - 0.7^2)^{-1/2}$ as in system i.

#### 3.2 Hypothesis testing

Correlation statistics, surrogate data tests, and the parametric tests were implemented as described in Methods. For the naive TTS test (Eq. 2) we set the flanking radius $r$ to 50. For the TTS test (Eq. 3) we set the flanking radius $r$ to 59, since the power of the TTS test is maximized when $r$ is set to one less than a multiple of 20, as in Fig 3.

#### 3.3 False positive rates of surrogate data tests without circularization

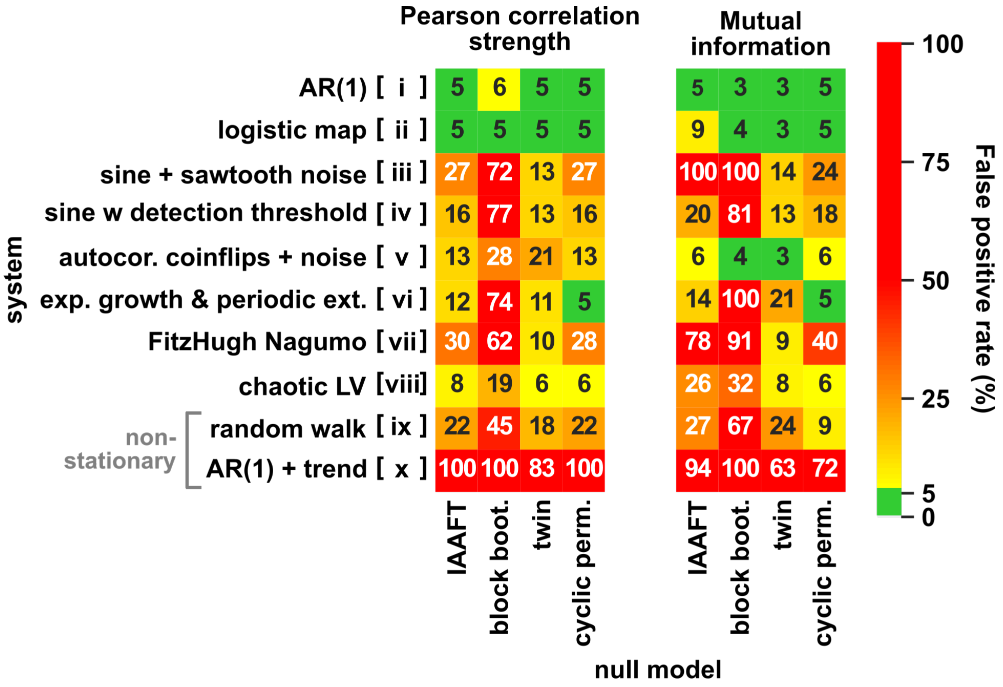

Figure S7: False positive rates of dependence tests without circularization. The same benchmark analysis was performed as in Fig 2 for IAAFT, block bootstrap, twin, and cyclic permutation tests, except that here, no circularization step was performed. Only these four surrogate tests are shown here because they were the only tests that used a circularization preprocessing step.

#### 3.4 Challenges for cyclic permutation surrogates and circularization preprocessing

The cyclic permutation method is fairly similar to the TTS test, but lacks the validity guarantee of the TTS test (as demonstrated in Fig 2). Because the two procedures are similar, we here try to gain some intuition for their sometimes different behavior by exploring specific ways in which stationary sequences can cause the cyclic permutation test to become invalid.

The cyclic permutation procedure is valid if: 1) the true process is periodic; 2) the initial condition was chosen at random from among the points within a period with equal chance, because that way, the ground truth process will be reflected by the cyclic permutation process (wherein each position of the period is sampled equally by surrogates); and 3) the length of the truncated time series is an integer multiple of the period.

Since cyclic permutation surrogates directly join the ends of sequences, it is intuitive that problems may arise when the beginning and end of a time series look very different, and therefore we may expect the cyclic permutation procedure to benefit substantially from circularization preprocessing [27, 13]. This occurs most strikingly in system vii (Fig 2), where the cyclic permutation test of mutual information has a false positive rate of either 5% with circularization (Fig 2) or 40% (Fig S7) without it. Thus, time series that are difficult to properly preprocess may pose challenges to the cyclic permutation method. Below, we illustrate two such examples. In both, we use the cyclic permutation surrogate test with circularization with the same parameters as in Fig 2.

First, a noise process or thresholding process can prevent the circularization procedure from selecting the optimal trimmed sequence length. As an example of this, consider a process in which $\{x_t\}$ and $\{y_t\}$ are generated by stationary sine waves with a detection threshold and additive noise:

$$x_t = \max(\sin(\phi_x + \frac{2\pi t}{35}), \beta) + \epsilon_{x,t}$$
$$y_t = \max(\sin(\phi_y + \frac{2\pi t}{35}), \beta) + \epsilon_{y,t}$$

Here, $\phi_x$ and $\phi_y$ are independent uniform random variables between 0 and $2\pi$. The noise terms $\epsilon_{x,t}$ and $\epsilon_{y,t}$ are independent normal random variables with zero mean and standard deviation of 0.05. The parameter $\beta$ is a threshold term. Example time series are shown in Fig S8A.

We generated independent $\{x_t\}$ and $\{y_t\}$ time series from this system and computed the proportion of trials in which the cyclic permutation test reported a significant Pearson correlation coefficient strength at the 0.05 significance level (Fig S8B purple). As the threshold increased, so did the false positive rate. This is presumably because at higher threshold values such as 0.5, much of the periodicity of the series is masked by the threshold, and so it is more difficult to determine the optimal length of the trimmed sequence via circularization (Eq. 4). Indeed, because the period of $\{y_t\}$ is 35, it would be natural to trim the sequence to a length that is a multiple of 35; this is nearly always acheieved for low threshold values, but rarely for high threshold values (Fig S8C). We can confirm that the increase in the false positive rate is due to a suboptimal trimming length because when we analyze only the trials in which the trimmed length was a multiple of 35, the false positive rate is correctly set at 0.05 regardless of the threshold value (Fig S8B, red). In this particular case, one may try to avoid this problem by estimating the period by other means, but this example is intended to illustrate the general idea that detection thresholds and noise processes may pose a challenge to circularization methods.

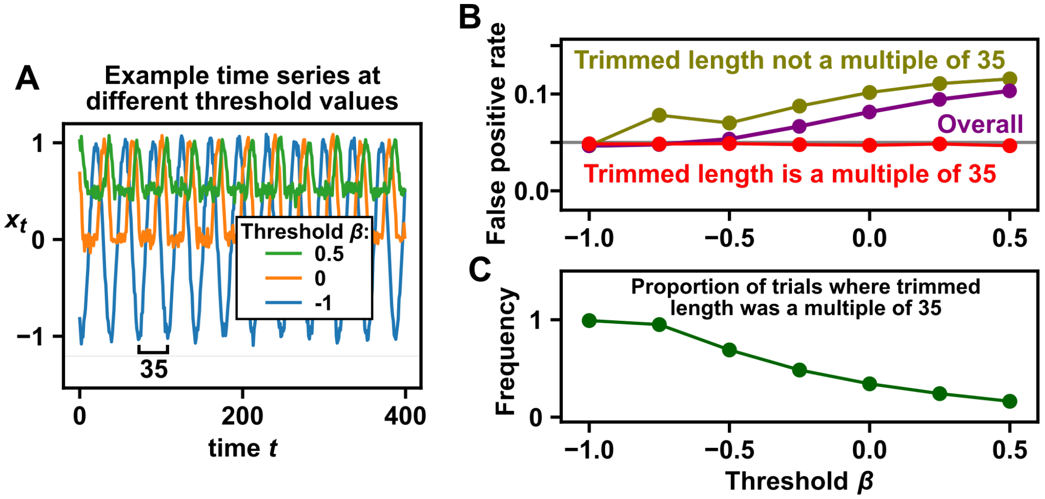

Figure S8: A process with a detection threshold is difficult for setting trimming length and thus poses a challenge for the cyclic permutation surrogate procedure. (A) Example time series of the thresholded sine process. A representative time series is shown for each of three possible threshold values. (B) False positive rate as a function of the detection threshold and the length of the trimmed time series. Specifically, we performed $10^5$ trials in which two time series were generated with the same threshold $\beta$. We used the cyclic permutation procedure (with the circularization trimming step) to test for the significance of the Pearson correlation strength at the 0.05 level. For all thresholds, there were at least $16,000$ trials in which the trimmed length was a multiple of 35. (C) The proportion of trials in which the trimmed (i.e. circularized) time series had a length that was a multiple of 35. The trimmed length is optimally a multiple of 35 because the dynamics have a period of 35.

A second potential challenge occurs when the method used to select the cutoff points ($k_{start}$ and $k_{end}$ in Eq. 4) is not appropriate for the time series under study. For instance, Eq. 4 chooses $(k_{start}, k_{end})$ as the values of $(k_1, k_2)$ that minimize the score:

$$\sum_{i=0}^{L}(y_{k_2+i} - y_{k_1+i})^2.$$

Although this score will identify subsequences at the beginning and the end of a time series that share similar values, these subsequences may not share other aspects of their distribution.

As a simple illustrative example, consider a system that toggles between a random even integer upper-bounded by the even number $\beta$ and a random odd integer upper-bounded by the odd number $\beta - 1$ . Specifically, let the phase term $\phi$ be either 0 or 1 with equal chance. Then, for $t = 1, ..., 400$, if $t$ has the same even/odd status as $\phi$ (i.e. $t$ and $\phi$ are either both even or both odd), $x_t$ will be a random even integer between 0 and $\beta$; alternatively, if $t$ does not have the same even/odd status as $t$, $x_t$ will be a random odd integer between 1 and $\beta - 1$. Lastly, because we plan to use mutual information to correlate two realizations of this process, we add a tiny amount of measurement noise (a uniform random variable between $-10^{-8}$ and $10^{-8}$) to each data point, as recommended by [50] for data sets with "tied" values. Fig S9A shows two examples of this process for $\beta = 6$. Note that this process is stationary.

We estimated the mutual information between an independent pair of such processes with the same value of $\beta$ and used the cyclic permutation procedure to test for significance at the 0.05 level. Since the process has a period of two, the circularization step should ideally trim the time series to an even length; doing so is both necessary and sufficient to ensure a false positive rate of 5% (Fig S9C). When $\beta = 2$, the possible even values (either 0 or 2) can be easily distinguished from the possible odd value (1 only), and thus the circularization procedure successfully trims the series to an even length, leading to a well calibrated false positive rate (Fig S9B, $\beta = 2$). As $\beta$ becomes larger (e.g. 6), the possible even values (0, 2, 4, or 6) cannot be easily distinguished from the possible odd values (1, 3, or 5), and thus circularization will sometimes trim the series to an odd length, leading to a high false positive rate (Fig S9B, $\beta = 6$).

When $\beta$ exceeds 10, the false positive rate begins to decline. This decline is not due to a decrease in the abundance of odd-trimmed sequences (green curve in B), but instead due to a decrease in the false positive rate among odd-trimmed sequences (olive curve in C). The explanation for this is beyond our present purpose, and will not be discussed further.

Overall, when time series have important fluctuations in aspects of their distributions that are not tied to distances between points, it may be difficult for the circularization scores such as Eq. 4 to select the optimal trimming length. We re-emphasize that the examples in both Fig S8 and Fig S9 used stationary processes. Thus, although these challenges are relevant to the circular permutation approach (and perhaps other techniques that rely on circularization), they will not result in inflated false positive rates under the TTS procedure.

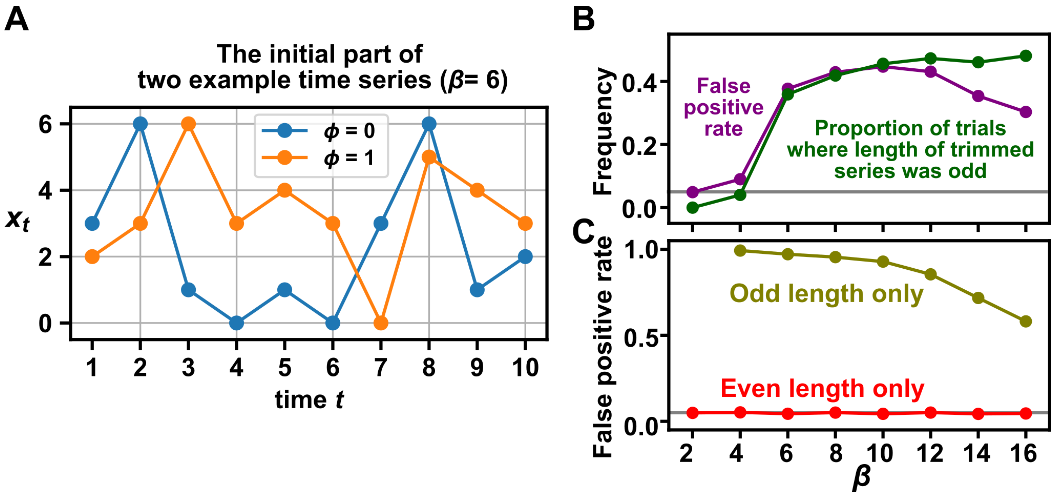

Figure S9: A process with important features that are not closely tied to interpoint distances frustrates a circularization procedure based on minimizing the distance between the beginning and the ending subsequences of a series. (A) A process $x$ toggles between a random even number (between 0 and $\beta$) and a random odd number (between 1 and $\beta - 1$) at each time step. See text for details. (B) We simulated an independent pair of realizations from this process at different values of $\beta$ and tested for significant mutual information at the 0.05 level using the cyclic permutation procedure (including the circularization step). We performed $10^4$ trials for each $\beta$. The false positive rate and the frequency of obtaining an odd-length trimmed sequence are shown. (C) The false positive rate as a function of the length of the trimmed time series. In both (B) and (C), a grey line indicates the 0.05 level.

### 4 Surrogates for some nonstationary time series: The detrend-retrend TTS test

Although the TTS test as described in Fig 1 requires that surrogates are generated from a stationary time series, it is possible to modify the TTS test for nonstationary time series that can be correctly decomposed into a deterministic trend component and a stationary component. The basic idea is that we can: (1) obtain the stationary component by subtracting the deterministic trend, (2) generate time-shifted surrogates from the stationary component, and (3) add the trend back to each of the time-shifted surrogates (Fig S10). This idea has previously been applied to random phase surrogates [56]. The next subsection describes the procedure precisely and show why it results in a valid test for independence. Afterward, simulations are used to illustrate scenarios where the this procedure is does (or does not) outperform simpler alternatives.

#### 4.1 Detailed description of the detrend-retrend TTS test

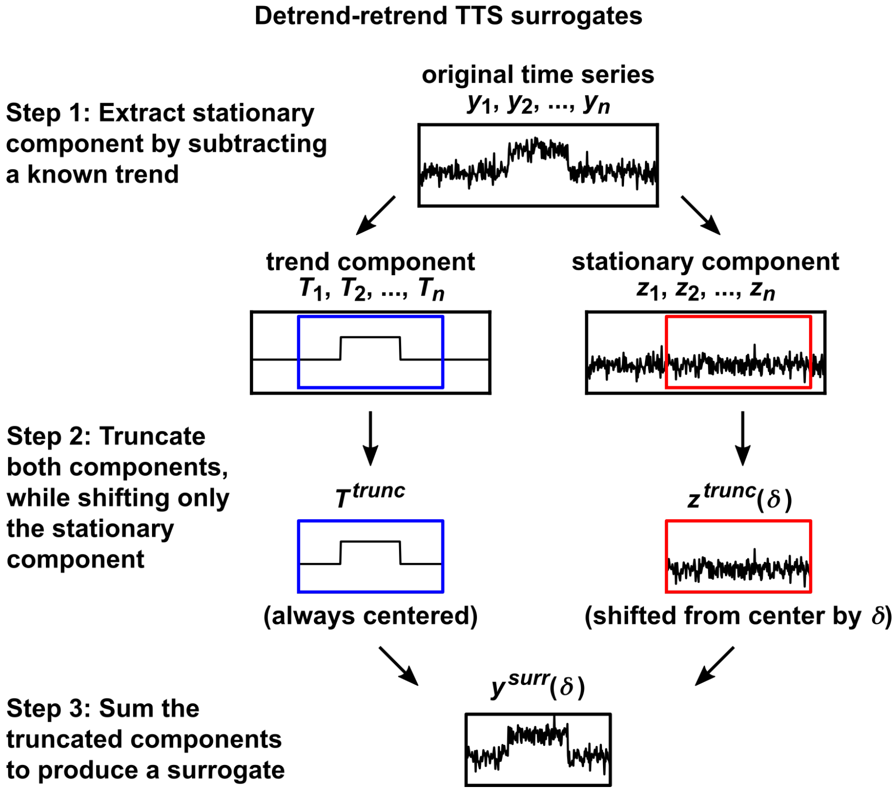

Figure S10: Procedure to generate detrend-retrend TTS surrogates.

We now describe this procedure (the "detrend-retrend TTS test") in detail. Suppose that there are two time series $\{x_t\} = \{x_1, x_2, ..., x_n\}$ and $\{y_t\} = \{y_1, y_2, ..., y_n\}$, and we wish to test whether they are independent. As in the regular TTS test (Fig 1), we choose a truncation radius $r$, a correlation function $\Theta$, and a significance level $\alpha$. Suppose now that $\{y_t\}$ is not stationary, but we know how to decompose it into a deterministic trend component and a stationary component. That is,

$$y_t = T_t + z_t \text{ for all } t = 1, 2, ..., n$$

where $\{T_1, ..., T_n\}$ is deterministic, $\{z_1, ..., z_n\}$ is stationary. We now produce truncated sequences from $\{x_t\}$, $\{T_t\}$ and $\{z_t\}$:

$$x^{trunc} = \{x_{1+r}, ..., x_{n-r}\}$$
$$T^{trunc} = \{T_{1+r}, ..., T_{n-r}\}$$
$$z^{trunc}(\delta) = \{z_{1+r+\delta}, ..., z_{n-r+\delta}\}$$

where, as before, $\delta$ takes on integers between $-r$ and $r$. Next, we produce $y$ surrogates $y^{surr}(\delta)$ as the element-wise sum of $z^{trunc}(\delta)$ and $T^{trunc}$ (Fig S10). That is, for each value of $\delta$, $y^{surr}(\delta)$ is the $(n-2r)$-length sequence whose $i$th element is given by:

$$T_{i+r} + z_{i+r+\delta}.$$

(We call these surrogates $y^{surr}(\delta)$ instead of $y^{trunc}(\delta)$ because they cannot be obtained by simply truncating $\{y_t\}$ at different shifts.) Next, we use these surrogates in the same way as before: Obtain the shifted correlation values

$$\theta_\delta = \Theta(x^{trunc}, y^{surr}(\delta)).$$

Let $B$ be the number of terms in the sequence $\{\theta_{-r}, ..., \theta_r\}$ that are greater than or equal to $\theta_0$. Let

$$u = B/(r+1).$$

Finally, the test detects dependence between $\{x_t\}$ and $\{y_t\}$ at a significance level of $\alpha$ if $u \leq \alpha$.

Let us see why the detrend-retrend TTS procedure is a valid test of dependence between $\{x_t\}$ and $\{y_t\}$ (as long as $\{y_t\}$ is the sum of a known deterministic component and a stationary component): First, note that the above procedure is equivalent to applying the (regular) TTS test for dependence between $\{x_t\}$ and the stationary component $\{z_t\}$, but with a special correlation function $\tilde{\Theta}$:

$$\tilde{\Theta}(x^{trunc}, z^{trunc}(\delta)) = \Theta(x^{trunc}, T^{trunc} + z^{trunc}(\delta)).$$

Thus, since $\{z_t\}$ is stationary, this procedure rigorously tests for dependence between $\{x_t\}$ and $\{z_t\}$. Next, since $\{y_t\} = \{T_t\} + \{z_t\}$ and since $\{T_t\}$ is nonrandom by definition, we know that $\{x_t\}$ and $\{z_t\}$ are dependent if and only if $\{x_t\}$ and $\{y_t\}$ are dependent. (Since this fact may not be immediately obvious, we have formally proven it below as lemma 14.) Thus, by testing for dependence between $\{x_t\}$ and $\{z_t\}$, the detrend-retrend TTS procedure also tests for dependence between $\{x_t\}$ and $\{y_t\}$, as promised.

**Lemma 14**

Let $\{x_t\}$, $\{y_t\}$, and $\{z_t\}$ be sequences of $n$ random variables such such that $\{y_t\} = \{T_t\} + \{z_t\}$, where $\{T_t\}$ is a sequence of nonrandom real numbers. Then, $\{x_t\}$ and $\{y_t\}$ are independent if and only if $\{x_t\}$ and $\{z_t\}$ are independent.

**Proof:** Let us begin with a notational change. Since our sequences have a finite length, we may instead represent them as vector-valued variables. Thus, let us rewrite our respective sequences as the length-$n$ random vectors $\vec{x}$, $\vec{y}$, $\vec{z}$, and the length-$n$ nonrandom vector $\vec{T}$. The proof essentially consists of applying 11 from Appendix 1.2. We first show the forward direction. To do so, suppose that $\vec{x}$ and $\vec{y}$ are independent. Define the functions $f(\vec{a}) = \vec{a}$ and $g(\vec{a}) = \vec{a} - \vec{T}$. By theorem 11, $f(\vec{x})$ and $g(\vec{y})$ are independent, but $f(\vec{x}) = \vec{x}$ and $g(\vec{y}) = \vec{z}$, so $\vec{x}$ and $\vec{z}$ are independent, as required. For the second reverse direction, suppose that suppose that $\vec{x}$ and $\vec{z}$ are independent. Define the function $h(\vec{a}) = \vec{a} + \vec{T}$. By theorem 11, $f(\vec{x}) = \vec{x}$ and $h(\vec{z}) = \vec{y}$ are independent, as required.

#### 4.2 With nonstationary data, the detrend-retrend TTS test is sometimes, but not always, superior to simpler alternatives.

We compared various flavors of the TTS test by applying them to simulated systems in which one or both series are nonstationary. We compared detrend-retrend TTS to two simpler strategies: directly applying the TTS test, or applying the TTS test after detrending the $\{y_t\}$ series (without retrending). The detrend-retrend test was superior in the case where both $\{x_t\}$ and $\{y_t\}$ are nonstationary, whereas the detrending-only strategy was superior where $\{x_t\}$ is stationary and $\{y_t\}$ is nonstationary. Below, we describe the details of the simulations.

We first simulated a system where $\{x_t\}$ and $\{y_t\}$ have related nonstationary behavior. We avoided using a system where the key problem addressed by the detrend-retrend TTS test is trivial: If $\{x_t\}$ and $\{y_t\}$ share a known common trend, then one could simply subtract this shared trend from both time series and proceed with the TTS test, obviating the need to "retrend". Avoiding that trivial case, $\{y_t\}$ has a visually apparent additive trend, but $\{x_t\}$ does not (Fig S11A).

For $t = 1, 2, ..., 400$, we generated $\{x_t\}$ and $\{y_t\}$ as follows:

$$H_t = \begin{cases} 5 & \text{if } 150 < t \leq 250 \\ 1 & \text{otherwise} \end{cases}$$

$$a_t = \begin{cases} 5 & \text{with probability } H_t/25 \\ 0 & \text{otherwise} \end{cases}$$

$$x_t = a_t + \epsilon_{x,t}$$

$$y_t = H_t + \epsilon_{y,t}$$

where the measurement noise terms $\epsilon_{x,t}$ and $\epsilon_{y,t}$ are normal random variables with a mean of 0 and variance of 1. For the case where $\{x_t\}$ and $\{y_t\}$ are independent, we set

$$\text{Cov}(\epsilon_{x,t}, \epsilon_{y,t}) = 0.$$

For the case where $\{x_t\}$ and $\{y_t\}$ are dependent, we set

$$\text{Cov}(\epsilon_{x,t}, \epsilon_{y,t}) = 0.3.$$

Both $\{x_t\}$ and $\{y_t\}$ are nonstationary. Whereas $\{y_t\}$ has a trend component given by $\{H_t\}$, the $\{x_t\}$ process does not have a deterministic trend component (Fig 1A).

We simulated $\{x_t\}$ and $\{y_t\}$ time series and tested whether they are dependent using various flavors of time-shift tests. Specifically, we either (1) tested for dependence between $\{x_t\}$ and $\{y_t\}$ using the (regular) TTS test, (2) tested for dependence between $\{x_t\}$ and the detrended series $\{\epsilon_{y,t}\}$ using the TTS test, or (3) tested for dependence between $\{x_t\}$ and $\{y_t\}$ using the detrend-retrend TTS test (i.e. detrending and retrending $\{H_t\}$ on $\{y_t\}$). For the different tests, we report the proportion of trials in which dependence was detected when the true covariance between $\epsilon_{x,t}$ and $\epsilon_{y,t}$ was either 0 (Fig S11B, "False positive rate") or 0.3 (Fig S11B, "True positive rate"). Directly applying the TTS test to $\{x_t\}$ and $\{y_t\}$ is invalid (since $\{y_t\}$ is nonstationary) and failed to control the false positive rate at the 0.05 significance level (Fig S11B top row). The other two tests are valid and correctly controlled the false positive rate. Of these, the detrend-retrend TTS test had higher detection power.

For the system where $\{x_t\}$ is stationary and $\{y_t\}$ is nonstationary (Fig S11C), we simulated an equivalent system except where $x_t$ is no longer determined by $H_t$:

$$a_t = \begin{cases} 5 & \text{with probability } 1/25 \\ 0 & \text{otherwise} \end{cases}$$

$$x_t = a_t + \epsilon_{x,t}$$

All other aspects of the system in Fig S11C are equivalent to those in Fig S11A. In this case, directly applying the TTS test did not result in a high false positive rate, and simply detrending gave grater detection power than the detrend-retrend TTS test (Fig S11D).

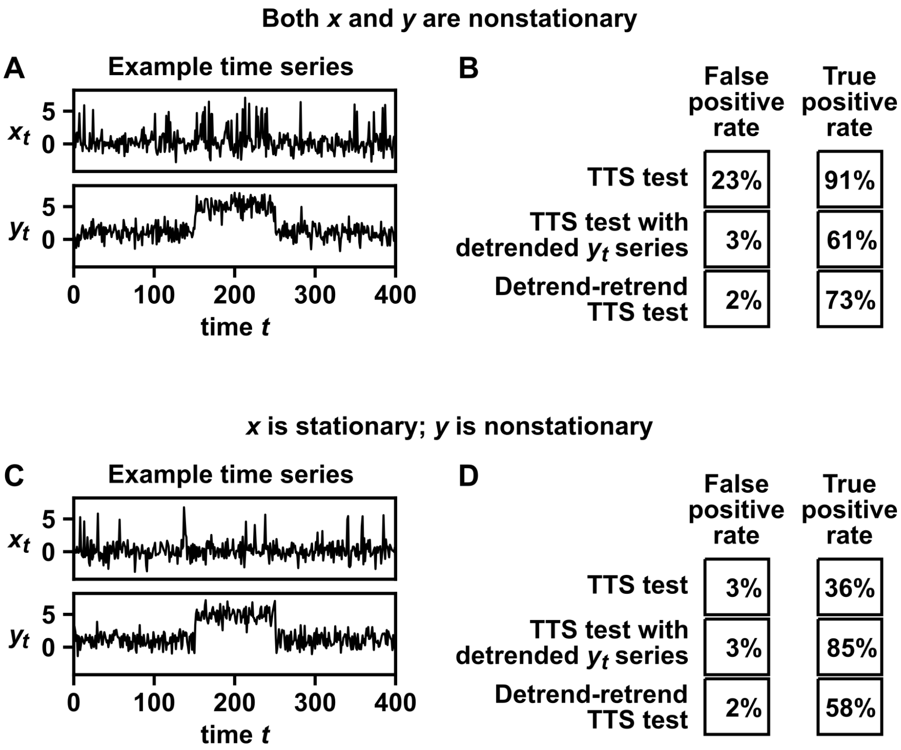

Figure S11: The detrend-retrend TTS test correctly controls the false positive rate in a benchmark simulation. (A) A simulated benchmark system in which both time series are nonstationary and where one time series ($\{y_t\}$) can be decomposed into a deterministic trend component and a stationary component. See text for simulation details. (B) Performance comparison. Different procedures were used to test whether $\{x_t\}$ and $\{y_t\}$ are dependent at the 0.05 significance level. Either the TTS test was applied directly, the TTS test was applied after detrending the $\{y_t\}$ series, or the detrend-retrend TTS test was used. To compute true positive rates, we set the covariance between the measurement noise of $x_t$ terms and measurement noise of $y_t$ to 0.3; for false positive rates this covariance was set to 0. For each test, the absolute value of the Pearson correlation coefficient was used as the correlation function, the truncation radius $r$ was set to 79, and true (or false) positive rates were reported as the proportion of $10^4$ trials in which dependence was detected. (C-D) Same as (A-B), but for a system where $\{x_t\}$ is stationary and $\{y_t\}$ is nonstationary.

### 5 Additional detection power comparisons between TTS and other tests

We performed several additional simulations examining the detection power of the TTS test, and comparing it to those of other surrogate data tests as well as the parametric test for Pearson correlation. In all of the following benchmark tests, we used the significance level of 0.05 to detect dependence. For all benchmark studies of the power of the TTS test (Fig 3, Fig S11 and this appendix section), we make sure to include $r = 79$, so that results are comparable across the various conditions. In Section 5.1, we also test $r = 19$ in order to study the effect of time series length (since a large $r$ is precluded in short series). In many of the following benchmark tests, we pre-shift data as in Fig 3. To fix notation, "pre-shifting by $s$" means that we test for dependence between $\{x_{1+s}, x_{2+s}, ..., x_n\}$ and $\{y_1, y_2, ..., y_{n-s}\}$ if $s \geq 0$, and test for dependence between $\{x_1, x_2, ..., x_{n-s}\}$ and $\{y_{1+s}, y_{2+s}, ..., y_n\}$ if $s < 0$.

#### 5.1 Unidirectionally coupled processes

**Autoregressive process:** We simulated the unidirectionally coupled autoregressive process:

$$x_t = \rho_x x_{t-1} + \rho_{yx} y_{t-1} + \epsilon_{x,t}$$
$$y_t = \rho_y y_{t-1} + \epsilon_{y,t}$$

where the $\epsilon_{x,t}$ and $\epsilon_{y,t}$ terms are independent random variables with a standard normal distribution. Unless otherwise specified, we used $\rho_x = 0.4$, $\rho_y = 0.4$, $\rho_{yx} = 0.3$, and a time series length of 400. We varied each of these four parameters ($\rho_x$, $\rho_y$, $\rho_{xy}$, and the time series length) individually. The initial conditions of $x$ and $y$ were chosen independently at random from a standard normal distribution and the system was allowed to equilibrate for 500 steps before recording measurements used for testing. We tested for dependence between the two time series using the Pearson correlation strength as the correlation statistic, and compared the detection power of each of the tests used in Fig 2. Additionally, we pre-shifted the $\{x_t\}$ series by $s = 1$ time step. For the TTS test, we used two different values for $r$ (79 and 19). We used $r = 79$ because this is an intermediate value as suggested in the main text and we used $r = 19$ in order to use the TTS test for time series lengths as low as 100. We used $r = 80$ for the naive TTS test as this is the multiple of 10 nearest to 79. For all other tests, details are the same as in Fig 3. $10^4$ trials were used to compute the true positive rate for each choice of parameters. The true positive rates are given in the file 2_coupled_ar1.xlsx.

**Coupled logistic map:** We simulated the unidirectionally coupled logistic map process:

$$x_t = x_{t-1}(r_x - r_x x_{t-1} - r_{yx} y_{t-1})$$
$$y_t = y_{t-1}(r_y - r_y y_{t-1})$$

where the initial conditions of $x$ and $y$ were chosen independently at random from a uniform distribution between 0.2 and 0.8, and the system was allowed to equilibrate for 500 steps before recording measurements used for testing. Unless otherwise specified, we used $r_x = 3.5$, $r_y = 3.8$, $r_{yx} = 0.1$ and a time series length of 400. We varied each of these four parameters ($\rho_x$, $\rho_y$, $\rho_{xy}$, and the time series length) individually. We tested for dependence between the two time series using cross-map skill with the same direction and parameters as in Fig 3 (except for when using the parametric test, which relies on Pearson correlation). Additionally, we pre-shifted the $\{x_t\}$ series by $s = 1$ time step. The values of $r$ for the TTS and naive TTS tests are the same as in the section immediately above. For all other tests, details are the same as in Fig 3. 2000 trials were used to compute the true positive rate for each choice of parameters. The true positive rates are given in the file 3_coupled_logistic.xlsx.

**Nonlinearly coupled autoregressive process:** We simulated a pair of autoregressive processes that are coupled in a nonlinear way:

$$x_t = \rho x_{t-1} + \cos(\epsilon_t)$$
$$y_t = \rho y_{t-1} + \sin(\epsilon_t)$$

where the $\epsilon_t$ terms are independent random variables drawn from a uniform distribution between 0 and $2\pi$. The state variables $x$ and $y$ were initialized at 0, and then the system was allowed to equilibrate for 500 steps before recording measurements used for testing. For the default system we used $\rho = 0.35$ and a time series length of 400. We either varied $\rho$ while keeping the length at 400 or varied the length while keeping $\rho$ at 0.35. We tested for dependence using the mutual information estimator (see Methods) and compared the detection power of each of the tests used in Fig 2. We did not pre-shift time series. The values of $r$ for the TTS and naive TTS tests are the same as in the sections immediately above. For all other tests, details are the same as in Fig 3. 2000 trials were used to compute the true positive rate for each choice of parameters. The true positive rates are given in the file 4_nonlin_coupled_ar1.xlsx.

#### 5.2 Bidirectionally coupled processes with random coupling delays

**Autoregressive process:** We simulated the bidirectionally coupled autoregressive process where $x$ and $y$ both influence each other with random strengths and random coupling delays, with iid Gaussian process noise of mean zero and variance 1:

$$x_t = 0.4x_{t-1} + \rho_{yx}y_{t-\tau_{yx}} + \epsilon_{xt}$$
$$y_t = 0.4y_{t-1} + \rho_{xy}x_{t-\tau_{xy}} + \epsilon_{yt}$$

The interaction strengths are chosen randomly from a continuous uniform distribution, and they are made sum to 0.4:

$$\rho_{yx} \sim \text{Unif}(0, 0.4)$$
$$\rho_{xy} = 0.4 - \rho_{yx}$$

Note that for any particular realization of the process, we have $\rho_{xy} \neq \rho_{yx}$ to mimic real world where variables are rarely symmetric.

The interaction delays $\tau_{yx}$ and $\tau_{xy}$ are drawn independently from the set $\{1, 2, ..., \tau_{max}\}$ with equal chance. Thus, $\tau_{max}$ determines the amount of uncertainty in the coupling delays. Here, we varied $\tau_{max}$ from 1 to 5. Additionally, the initial conditions of $x$ and $y$ were chosen independently at random from a standard normal distribution and the system was allowed to equilibrate for 500 steps before recording measurements used for testing.

We tested for dependence using the absolute value of the Pearson correlation between $\{x_t\}$ and $\{y_t\}$. To account for the delay uncertainty, we pre-shifted data by several choices of $s$ and performed a multiple-testing correction. We tried different numbers of pre-shifts centered at $s = 0$. That is, we preshifted by:

$$-s_{max}, ..., s_{max}$$

and performed a Bonferroni correction for the number of pre-shifts ($= 2s_{max} + 1$). We varied $s_{max}$ from 1 to 5. Larger values of $s_{max}$ correspond to an assumption of greater uncertainty in the coupling delay. As in Fig 3, we used the lowest value of $r$ that would enable significance at the 0.05 level after the Bonferroni correction (i.e. $r = 59$ for 3 pre-shifts, $r = 99$ for 5 pre-shifts, and so on). In order to handle up to 11 different choices of $s$ (and thus 11 different hypothesis tests to correct for), we used a time series length of 800. We used the parametric test for Pearson correlation and IAAFT surrogate test for comparison. We used 1499 IAAFT surrogates rather than 99, to account for the lower $p$-values demanded by the Bonferroni correction. To calculate true positive rates, we repeated the trial $10^4$ times for TTS and parametric tests, and 500 times for the slower IAAFT test. The true positive rates are given in the file 5_bidirectional_ar.xlsx.

**Logistic map:** We simulated the bidirectionally coupled logistic map process where $x$ and $y$ influence each other with random coupling delays:

$$x_t = x_{t-1}(r_x - r_x x_{t-1} - 0.1y_{t-\tau_{yx}})$$
$$y_t = y_{t-1}(r_y - r_y y_{t-1} - 0.1x_{t-\tau_{xy}})$$

where the initial conditions of $x$ and $y$ were chosen independently at random from a uniform distribution between 0.2 and 0.8, and the system was allowed to equilibrate for 500 steps before recording measurements used for testing. The terms $r_x$ and $r_y$ are randomly drawn from the set:

$$\{3.7, 3.72, 3.74, 3.76, 3.78, 3.8\}$$

without replacement (i.e. $r_x \neq r_y$). We require $r_x \neq r_y$ to avoid pathological synchrony [6, 12]. As in the above section, the interaction delays $\tau_{yx}$ and $\tau_{xy}$ are randomly drawn independently from the set $\{1, 2, ..., \tau_{max}\}$, and we varied $\tau_{max}$ from 1 to 5. Again, we used a time series length of 800 for this benchmark.

We tested for dependence between the two time series using cross-map skill with the same direction and parameters as in Fig 3 (except for when using the parametric test, which relies on Pearson correlation). As in the section above, we used the TTS test with multiple pre-shift values and a Bonferroni multiple test correction, and the IAAFT and parametric correlation tests. All other details of the analysis, such as choices of $r$ and trial numbers, are the same as in the above section. The true positive rates are given in the file 6_bidirectional_logistic.xlsx.

### 6 Detailed methods and results for the orbital-climate dependence example

#### 6.1 Obtaining orbital parameter time series and deglaciation event times.

We obtained obliquity, precession, and eccentricity time series from [60] using the web interface at `http://vo.imcce.fr/insola/earth/online/earth/online/index.php`, with a sampling frequency of 1 kyr. The solar constant was left as the default value of 1368 watts per square meter. Deglaciation event times [65] were obtained from `https://www.ncei.noaa.gov/pub/data/paleo/contributions_by_author/huybers2006/huybers2006.txt`.

#### 6.2 Testing for dependence using a nonlinear correlation statistic

In Fig 4 we used a correlation function based on state space reconstruction (SSR) with delay vectors. One of the most popular state space reconstruction correlation statistics is the so-called "cross-map skill" of the convergent cross-mapping approach [6]. Our technique, while inspired by cross-map skill, deviates from it in two ways: the weighting function and the number of neighbors used for prediction. We first describe this correlation function and use a simulation to justify why these changes are important for the orbital-climate dependence example.

To obtain a correlation between two time series $\{x_t\}$ and $\{z_t\}$, we begin by constructing delay vectors from the $\{z_t\}$ series:

$$v_t = (z_t, z_{t-D}, ..., z_{t-(E-1)D})$$

where $D$ is the delay amount and $E$ is the embedding dimension (we will discuss the choice of the parameters $D$ and $E$ later). $v_t$ is thus a point in an $E$-dimensional "delay space". We now have the paired time series $\{x_t, v_t\}_{t=1}^{n}$. Next, we try to predict the $\{x_t\}$ series using the delay vectors $\{v_t\}$. Specifically for each value of $i = 1, 2, ..., n$, we find the $k$ nearest neighbors of $v_i$ in the delay space. Note that $v_i$ cannot be its own neighbor. We use these nearest neighbors to predict $x_i$:

$$\widehat{x}_i = \frac{\sum_{j=1}^{n} x_j w(v_i, v_j) I_k(i,j)}{\sum_{j=1}^{n} w(v_i, v_j) I_k(i,j)}$$

$$I_k(i,j) = \begin{cases} 1 & \text{if } v_j \text{ is one of } v_i\text{'s } k \text{ nearest neighbors} \\ 0 & \text{otherwise} \end{cases}$$

where $\widehat{x}_i$ is the predicted value of $x_i$ and $w(v_i, v_j)$ is a weighting function which is larger when $v_i$ and $v_j$ are closer together (to be discussed further below). Our correlation statistic is then given by the negative mean squared error of the predictions:

$$-\frac{1}{n}\sum(x_i - \hat{x}_i)^2. \tag{7}$$

We choose negative error for the statistic so that higher values of the statistic (lower error) will correspond to stronger coupling signal, as per the TTS test. The popular cross-map skill procedure [6] typically uses an exponential weighting function:

$$w_{exp}(v_i, v_j) = e^{-|v_i - v_j|/d_i} \tag{8}$$

where $|v_i - v_j|$ is the Euclidean distance from $v_i$ to $v_j$ and where $d_i$ is the Euclidean distance from $v_i$ to its closest neighbor. However, we found that these choices do not perform well for time series that resemble the deglaciation series (Fig S12D). Specifically, our $\{x_t\}$ series (i.e. the series being predicted) is a series of 2000 kiloyears in which 36 deglaciations occurred. That is, 36 values of the $\{x_t\}$ series are 1s and the rest are 0s. We have found that in simulations where $\{x_t\}$ is a sparse event time series, the exponential weight function has abysmal detection power. Instead, here we use a simple inverse-distance weighting function:

$$w_{inv}(v_i, v_j) = 1/|v_i - v_j| \tag{9}$$

which we found to have high detection power for sparse event time series (Fig S12D).

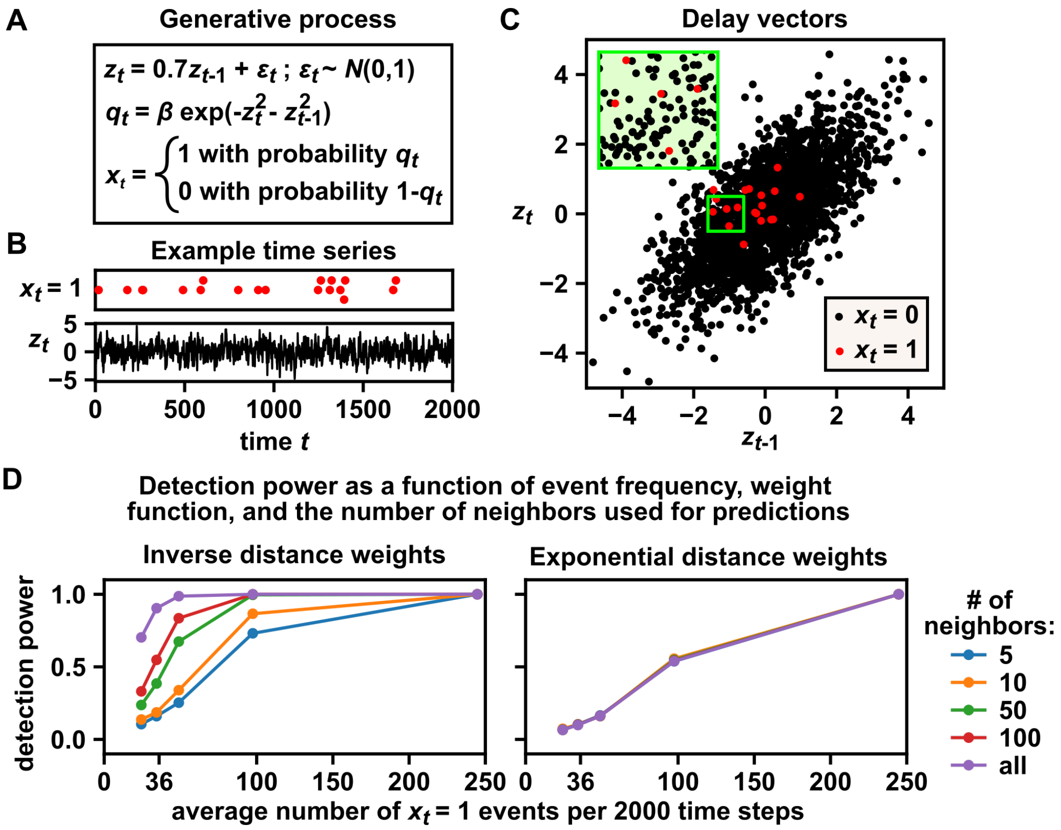

Figure S12: Inverse distance weights and larger numbers of neighbors improve detection power for a simulated sparse event time series. (A) $\{z_t\}$ was generated according to a simple first-order autoregressive process. $\{x_t\}$ is an event time series ($x_t$ is either 1 or 0) and depends on $\{z_t\}$, with 1 occurring more frequently when $z_t$ and $z_{t-1}$ are near the origin. The parameter $\beta$ determines the frequency of events. (B, C) Time series and delay vector plot of a realization with $\beta = 0.05$. In the event time series, overlapping dots are vertically separated so that they can be distinguished (i.e. the vertical axis is meaningless). Delay vectors were constructed as: $v_t = (z_t, z_{t-1})$. When events are sparse, event points are generally surrounded by non-event points. Note that events are clustered around the origin. Thus, nearest-neighbor prediction techniques with a small number of nearest neighbors may suffer poor prediction accuracy. (D) The TTS test with various SSR-based correlation functions (Eq. 7) was used to detect the dependence between $\{x_t\}$ and $\{v_t\}$. Time series were of length 2198 and $r$ was set to 99, so that truncated $\{x_t\}$ and $\{v_t\}$ series would be of length 2000, mirroring the length of the deglaciation series. The event frequency parameter $\beta$ was set to 0.05, 0.07, 0.1, 0.2, and 0.5, which correspond to an average of 24, 34, 49, 97, and 245 events within the length-2000 truncated window. (The deglaciation time series has 36 events among its 2000 time points). The detection power (i.e. the proportion of 1000 trials in which the dependence between $\{x_t\}$ and $\{z_t\}$ was detected at the 0.05 significance level) is shown as a function of the average number of events within the truncated $\{x\}$ window. Either inverse distance weights (Eq. 9) or exponential distance weights (Eq. 8) were used and the number of neighbors used for prediction (i.e. $k$) was varied from 5 to all possible neighbors (1999 in this case). For sparse event series, inverse distance weighting with all possible neighbors provides substantially higher power than all other options. Curves for exponential distance weights are superimposed because the number of neighbors did not affect detection power.

In order to compute correlations using this approach, we must pick embedding parameters (the delay lag $D$ and the embedding dimension $E$) for each orbital parameter time series. As is typical, we selected embedding parameters that maximized predictions of future values using past values of the same time series. There are three further considerations in our embedding parameter selection procedure: (1) Which portion of the data did we use as training data to select embedding parameters? (2) We chose embedding parameters to minimize the error in forecasts $q$ kyr into the future; what was $q$? (3) Which possible parameter values did we search over and what other hyperparameters were needed? Below we discuss each of these in turn.

Which data did we use for parameter value selection? We randomly selected 25 windows of 2 Myr (the same as the length of the deglaciation time series) from between $-12$ Myr and $+10$ Myr (where 0 is the present time). For each 2-Myr window (2000 data points), we constructed delay vectors from the orbital parameter time series and used the SSR correlation function to predict future values of the same time series. We thus obtained 25 mean squared error values (one for each window). Embedding parameters were chosen to minimize the average over all 25 mean squared error values.

Embedding parameters were chosen by minimizing the error of future predictions; how far into the future (forecast horizon) did we predict? We chose the forecast horizon to be a fixed proportion of the cycle period instead of a fixed amount of time in order to subject time series with different autocorrelation structures to prediction problems of similar "difficulty". Cycle periods for obliquity, precession and eccentricity, estimated as average interpeak distances, were 41 kyr, 21 kyr, and 99 kyr respectively (broadly consistent with known values [61]). Thus, for embedding parameter selection, we used forecasts of 8, 4, and 20 kyr ahead for obliquity, precession, and eccentricity respectively.

Which possible embedding parameter values did we search over? We scanned embedding dimensions from 2 to 6 and scanned delay lags from 1 to 55 kyr, and chose embedding parameters that minimized average prediction error. Also, since these predictions were on continuous time series (not event data), we used the standard [6] exponential distance weights (Eq. 8) and used $k = 5$ nearest neighbors for predictions.

From this procedure with all of the above ingredients, we obtained delay vector parameters for obliquity ($E = 6$ kyr, $D = 31$ kyr), precession ($E = 5$ kyr, $D = 4$ kyr), and eccentricity ($E = 6$ kyr, $D = 45$ kyr).

We then computed the nonlinear correlation statistic (Eq. 7 and 9) with all neighbors (i.e. all delay vectors except self, or equivalently, $k = 1999$) by predicting deglaciation events based on the delay vectors of orbital parameters. Finally, we used the TTS test to assess the significance of the statistic. For the $x^{trunc}$ window we used the 2000 kyr of deglaciation event data ($-1999$ to 0 kyr) from [65], with event times rounded to the nearest kyr. We used a flanking radius of $10,000$ kyr.

#### 6.3 Statistical power and effect of pre-shift under a simulated dependence model

We expected that small pre-shifts would not dramatically alter power because the orbital parameter time series exhibit deterministic dynamics and thus delay vectors occurring at nearby times would likely share similar amounts of information about deglaciation. To verify this intuition, we simulated a possible model of dependence between orbital parameters and deglaciation, and performed TTS tests with various pre-shifts.

For these simulations, we could in principle construct a model where deglaciation and one of the orbital parameters are dependent with various coupling lags (but no preshift), and assess how varying the true coupling lag would affect detection power. However, repeating the simulation for many different coupling lags is computationally expensive. As a computationally cheaper (but essentially equivalent) alternative, we constructed a model where the dependence has no coupling lag, and then tested how preshifting might affect detection power. Our model incorporates three features from the deglaciation data set: the number of deglaciation events in the dataset ($n_{event} = 36$), the average measurement uncertainty, and the minimum observed time between deglaciation events. Additionally, we constructed the model such that the correlation statistic described above would be well-positioned to capture the dependence relationship. Below, we describe the model for obliquity, but the steps are analogous for precession and eccentricity.

**1.** Select a value for the parameter $k$, which will set the minimum time between deglaciation events. We use $k = 24$ kyr because this is the minimum time between two deglaciation events in the data set.

**2.** Choose a random time $t^*$ from among the deglaciation time range ($-1999$ to 0 kyr), and designate this time $t^*$ as the "anchoring" deglaciation event. The anchoring deglaciation event will play a central role in determining all other deglaciation events.

**3.** Find the corresponding $D$-dimensional anchoring obliquity delay vector $\overrightarrow{z}(t^*) = [z(t^*), z(t^*-E), ..., z(t^*-(D-1)E)]$. See section 6.2 for the values of $D$ and $E$.

To model dependence between deglaciation and obliquity, the rest of the procedure assigns the remaining deglaciation events to the obliquity delay vectors that are most similar to the anchoring obliquity delay vector (while obeying the constraint that all deglaciations must be separated in time by at least $k$). This captures the assumption that deglaciations occur at times whose obliquity delay vectors resemble the anchoring delay vector $\overrightarrow{z}(t^*)$.

**4.** Let $\overrightarrow{\boldsymbol{Z}}$ be the entire list of obliquity delay vectors, sorted by Euclidean distance to $\overrightarrow{z}(t^*)$, from least to greatest distance. In other words, $\overrightarrow{\boldsymbol{Z}}(1)$ is $\overrightarrow{z}(t^*)$ itself, $\overrightarrow{\boldsymbol{Z}}(2)$ is the nearest neighbor of $\overrightarrow{z}(t^*)$ in the $D$-dimensional delay space, $\overrightarrow{\boldsymbol{Z}}(3)$ is the second-nearest neighbor of $\overrightarrow{z}(t^*)$ and so on.

**5.** Delete from $\overrightarrow{\boldsymbol{Z}}$ all delay vectors $\overrightarrow{z}(t)$ for which $|t^* - t| < k$. This step, along with step 7, enforces the requirement that $k$ is the minimum time between deglaciation events.

**6.** Let $\overrightarrow{z}(t')$ be the (new) first entry of $\overrightarrow{\boldsymbol{Z}}$. Designate $t'$ as a deglaciation event.

**7.** Delete from $\overrightarrow{\boldsymbol{Z}}$ all delay vectors $\overrightarrow{z}(t)$ for which $|t' - t| < k$.

**8.** Repeat steps 6-7 until $n_{event}$ deglaciation events have been designated.

This left us with a list of 36 deglaciation times, each between $-1999$ and 0 kyr. Lastly, to model measurement noise, we added Gaussian random noise with zero mean and standard deviation of 6.14 kyr (a specially chosen value whose origin we explain below) to each deglaciation time, and then rounded to an integer in units of kyr. Any values below $-1999$ kyr or above 0 kyr were simply set to $-1999$ or 0 respectively.

To explain where the standard deviation of 6.14 kyr (variance of 37.7 kyr$^2$) comes from, first note that the set of deglaciation times was derived from a time series of $\delta^{18}O$ measurements (a proxy for water temperature) [65]. During the past 2000 kyr, the variance due to uncertainty in $\delta^{18}O$ measurements was on average 75.5 kyr$^2$. Huybers designated each deglaciation $t_{deglac}$ as the midpoint in time between a local minimum in temperature $t_{cold}$ and a local maximum in temperature $t_{hot}$. That is, $t_{deglac} = \frac{1}{2}(t_{cold} + t_{hot})$. Under the simplifying assumption that $t_{hot}$ and $t_{cold}$ are independent random variables each with a variance of 75.5 kyr$^2$, the variance of $t_{deglac}$ is $(2 \times 75.5)/2^2 \approx 37.7$ kyr$^2$, which is the value used for deglaciation time uncertainty in our simulations.

Using this generating model, we performed 100 simulations of dependence between each orbital parameter and deglaciation, and we tested for the dependence relationship using the TTS test with the nonlinear correlation statistic described in the previous section. By construction, this generating model assumes that there is no coupling lag between orbital parameters and deglaciation. To assess the effect of a coupling lag, we pre-shifted orbital time series by up to 65 kyr in either direction (Fig S13B). For obliquity (Fig S13B, blue), power is insensitive to preshift. For precession (Fig S13A, orange), power does not seem to be dramatically decreased by pre-shifts up to the precession-versus-ice-volume lags observed by Imbrie et al. [66] (6 kyr). Detection of the deglaciation-eccentricity dependence also maintains good power up to about a pre-shift of 9 kyr (Fig S13A, green).

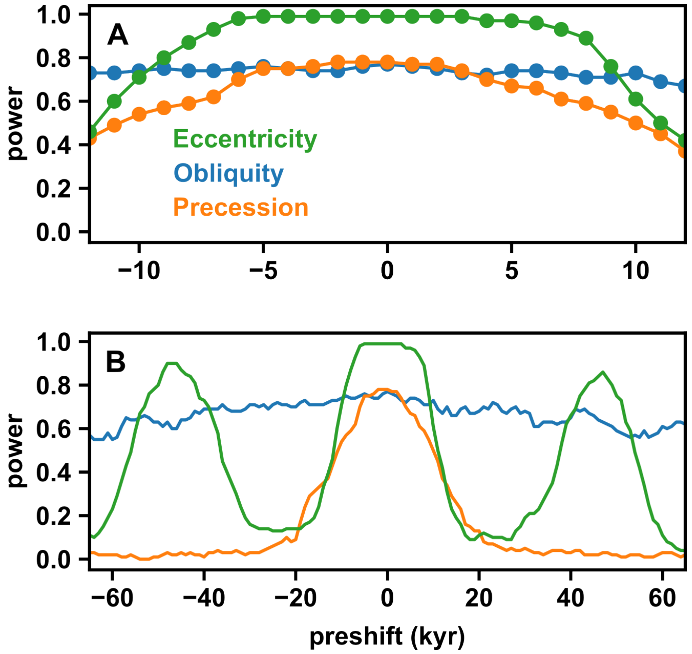

Figure S13: Effect of pre-shifts on the statistical power of the TTS test in simulated models of orbital-deglaciation dependence. (A) Power for all three orbital parameters is mostly stable for preshifts up to $\sim 6$ kyr in either direction. (B) Power is mostly stable over larger pre-shifts for obliquity, but not precession or eccentricity. A pre-shift value of $s$ refers to a test that tries to correct for a coupling lag where deglaciation dynamics at time $t$ are best estimated by orbital delay vectors at time $t + s$.

### 7 Detailed methods and results for the cross-site microbiome dependence example

#### 7.1 Testing for dependence between population at the single-species level is impeded by compositional data.

Microbiome surveys often rely on 16S ribosomal RNA sequencing approaches in which relative abundance levels of taxa are observed (“compositional data”), but not absolute abundance levels [72]. Because of this, detecting species-level dependence relationships directly from data is difficult (Fig S14). Specifically, it is difficult to make valid species-level inferences about dependence between subpopulations using relative abundance data, both within a body site (Fig S14B-iii) and between body sites (Fig S14B-iv). For this reason, we focus on testing for dependence between body sites as a whole, a task that is not impeded by compositional data, as we prove in the following subsection.

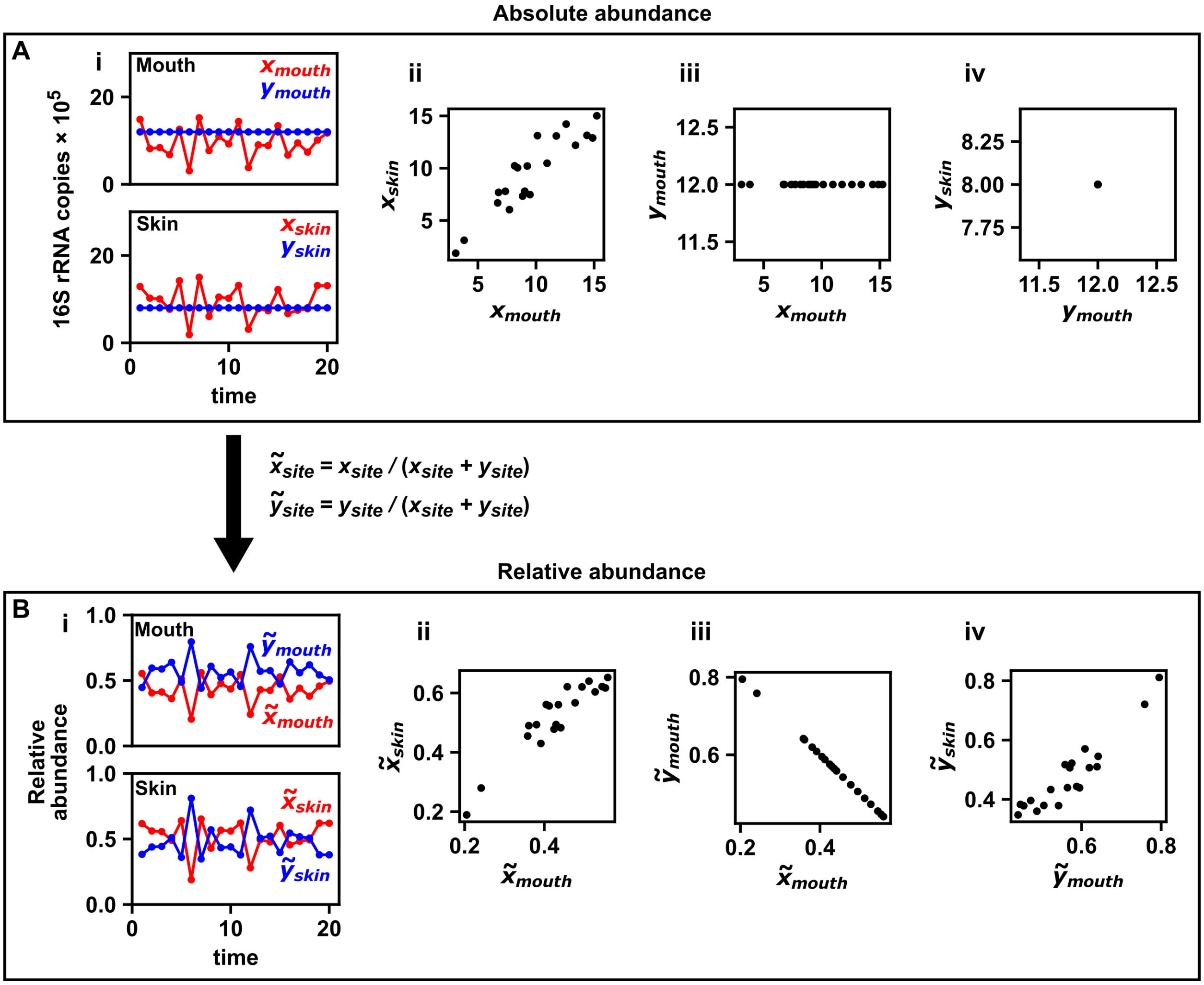

Figure S14: Naive species-level analysis of relative abundance data can result in spurious correlations. (A) Absolute abundance estimates of 16S rRNA copy numbers (a proxy for biomass) of two microbial species ($x$ and $y$) measured over time at two body sites (mouth and skin). (A-i) shows that $x_{mouth}$ and $x_{skin}$ covary, whereas $y_{mouth}$ and $y_{skin}$ stay constant over time. Scatter plots show that $x$ is correlated between the two body sites (A-ii), whereas there is no evidence of a correlation between the $x$ and $y$ populations in a single site (e.g. A-iii), and there is also no evidence of a correlation between the $y$ populations at the two sites (A-iv). (B) Same as A, but now using relative abundance measurements ($\tilde{x}$ and $\tilde{y}$), calculated as $\tilde{x}_{site} = x_{site}/(x_{site} + y_{site})$. The original correlation still exists (B-ii), but now spurious correlations have appeared, both within (B-iii) and between (B-iv) body sites.

#### 7.2 Testing for dependence between whole microbial communities at different body sites is not impeded by compositional data

Here we formally show that testing for dependence between the microbial communities of two body sites is not impeded by compositional data. The argument relies primarily upon Theorem 11 from Appendix 1.2.

Let the absolute population densities of $m$ taxa at a certain body site be given by the matrix $X$ where $X_{i,t}$ is the density of the $i$th taxon on day $t$. Then, the compositional (or relative abundance) matrix $h(X)$ is given by:

$$(h(X))_{i,t} = \frac{X_{i,t}}{\sum_{j=1}^{m} X_{j,t}}.$$

As long as a nonzero total microbial load is measured on each day, $h$ is continuous and therefore measurable. Thus we can apply Theorem 11. In particular, if $X$ and $Y$ are independent random variables, then their compositional counterparts $h(X)$ and $h(Y)$ must also be independent. In fact, the contrapositive of this statement is more useful for statistical testing: If we can show by statistical testing that $h(X)$ and $h(Y)$ are dependent, then it follows that $X$ and $Y$ are also dependent.

#### 7.3 Justification of median intraspecies correlation

In the microbiome example (Fig 5), we use the TTS procedure paired with a custom statistic (here called the "median intraspecies correlation") to test for dependence between the microbiomes at different body sites across time. Specifically, for two body sites $X$ and $Y$, we had time series of the relative abundance of all shared bacterial species. For the $i$th bacterial species, we computed the Pearson correlation between the time series of the $i$th species in sites $X$ and $Y$; the median intraspecies correlation statistic is then the median of all such correlations. The idea behind this statistic is that (as suggested by Caporaso et al. [71]) species-level correlations between body sites likely arise from frequent transfers of bacteria among body sites (such as from hands rubbing together). In this case, we expect sites that transfer bacteria among one another to have primarily positive correlations. Thus, the median correlation seems to be a sensible statistic.

In this section, we verify the above intuition using a simulation that captures essential properties of the system (e.g. frequent transfer among sites and compositional data). We also compare the median intraspecies correlation to the canonical correlation, which is another way to perform a correlation between two vectors.

For this simulation, we model population dynamics as a first order autoregressive process:

$$\begin{aligned} X_{i,t+1} =& 5 + 0.9X_{i,t} + r_{YX} y_{i,t} + \epsilon_{X,i,t} \\ Y_{i,t+1} =& 5 + 0.9Y_{i,t} + r_{XY} X_{i,t} + \epsilon_{Y,i,t} \end{aligned}$$

where $X_{i,t}$ is the population size of species $i$ in site $X$ at time $t$ and $Y_{i,t}$ is the same but for site $Y$. The noise terms $\epsilon_{X,i,t}$ and $\epsilon_{Y,i,t}$ are iid standard normal random variables. This model captures the mixing of populations between different sites via the cross-terms $r_{YX}$ and $r_{XY}$. We model bidirectional transfer by setting

$$r_{YX} = r_{XY} = 0.03$$

and we model unidirectional transfer by setting

$$r_{YX} = 0.03 \text{ and } r_{XY} = 0.$$

Since the microbiome data studied in this work are relative abundance values (i.e. compositional), we apply the compositional transformation to simulate the measurement process:

$$\tilde{X}_{i,t} = \frac{X_{i,t}}{\sum_{j=1}^{m} X_{j,t}}$$

where $m$ is the number of species. Finally, since the data set we used has a sampling frequency of once per day [71], and mixing events (e.g. hand-to-mouth, hand-to-hand) likely occur more frequently than once per day, we subsample the data. Thus, the time series used in this simulation are 400 time points taken once every 4 time steps. That is, the relative abundance measurements of species $i$ at sites $X$ and $Y$ are given by:

$$\begin{aligned} &\{\tilde{X}_{i,4}, \tilde{X}_{i,8}, \tilde{X}_{i,12}, ..., \tilde{X}_{i,1604}\} \\ &\{\tilde{Y}_{i,4}, \tilde{Y}_{i,8}, \tilde{Y}_{i,12}, ..., \tilde{Y}_{i,1604}\}. \end{aligned} \tag{10}$$

We use $m = 50$ species, which is within the range of the observed shared species counts (Fig S16A&D). This subsampling step was used for all simulations and benchmarks in this section. Fig S15 shows that for both bidirectional transfer and unidirectional transfer, the median Pearson correlation is positive, as expected.

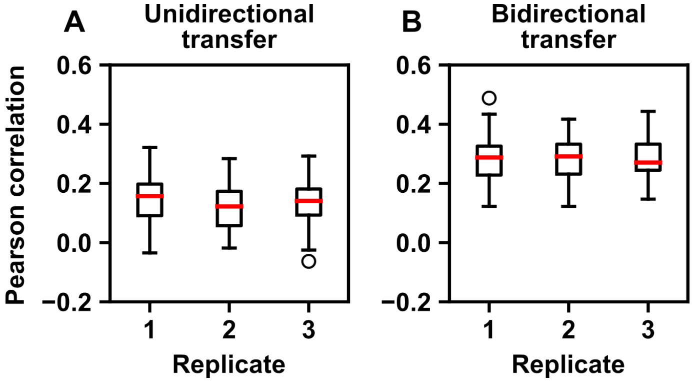

Figure S15: Distribution of intraspecies Pearson correlation coefficients in a simulation of cross-site dependence. (A) Pearson correlations between $\{\tilde{X}_{i,t}\}$ and $\{\tilde{Y}_{i,t}\}$ (i.e. Eq. 10) in the case of unidirectional transfer, where $r_{YX} = 0.03$ and $r_{XY} = 0$. Each box plot shows the distribution of intraspecies Pearson correlation coefficients from a single replicate simulation. Since $m = 50$, each box plot represents 50 numbers. The median of the box plot, shown in red, represents the median intraspecies correlation (i.e. the statistic used in Fig 5). (B), as in A, but for the case of bidirectional transfer, where $r_{YX} = r_{XY} = 0.03$.

We assessed how well the TTS test could detect dependence between $X$ and $Y$ when paired with either the median intraspecies Pearson correlation coefficient, or the "first canonical correlation", which is an alternative way of calculating a correlation from two matrices [117, 118]. To explain the latter method, let $W$ and $Z$ both be $m \times n$ data matrices with $m$ variables and $n$ measurements. The first canonical correlation of $W$ and $Z$ is given by:

$$\max_{(a,b)}(\mathrm{Corr}(a^T W, b^T Z))$$

where $a^T$ and $b^T$ are both $m$-dimensional row vectors, and Corr is the sample Pearson correlation. In other words, the first canonical correlation is the maximum Pearson correlation between a linear combination of $W$ and a linear combination of $Z$. As a technical detail, since the values within the relative abundance matrices are linearly dependent (due to the compositional transformation), taking the canonical correlation of $\tilde{X}$ and $\tilde{Y}$ can lead to numerical conditioning issues. Therefore, we arbitrarily delete one of the $m$ species from $\tilde{X}$ and $\tilde{Y}$ before computing the first canonical correlation.

Using the system of Eq. 10, we determined the power of the TTS test with either the median intraspecies Pearson correlation coefficient or the first canonical correlation (as implemented in the 'CanCorr' function in the Statsmodels package [75]). We used $r = 79$ and calculated power as the proportion of 25 simulations in which dependence was detected at the 0.05 level. Using the median intraspecies Pearson correlation, the TTS test detected dependence in $19/25 = 76\%$ of simulations in the case of unidirectional transfer, and $25/25 = 100\%$ of simulations with bidirectional directional transfer. Using the first canonical correlation as the correlation statistic, the TTS test detected dependence in $1/25 = 4\%$ or $2/25 = 8\%$ of simulations in the unidirectional and bidirectional cases respectively. Overall, the median intraspecies correlation provided superior performance, justifying its use in Fig 5 instead of the canonical correlation approach.

#### 7.4 Obtaining OTU tables

The OTU count table and sample information were obtained from the Qiita platform [73]. The OTU table we used is available in BIOM format at https://qiita.ucsd.edu/download/219436. A file with sample information (such as collection dates, body sites, and the human subject a sample was taken from) is available at https://qiita.ucsd.edu/download/773810. Microbiome survey time series were reported from two human subjects (M3 and F4) [71]. We used data from subject M3 as the time series are longer.

The OTU table was generated by a standard workflow that trimmed sequences to 90 base pairs and then generated an OTU table by a closed reference picking method. This workflow can be viewed in a visual flowchart at https://qiita.ucsd.edu/study/description/550. The OTU table ID is 44973.

#### 7.5 Local similarity analysis

Local similarity analysis has been implemented in various different ways [5, 74]. We used the original procedure of [5] with the maximum delay parameter set to zero. Specifically, to compute the local similarity score of two time series $s(\{x\}_1^n, \{y\}_1^n)$, we begin by taking the rank of each time series. Let $r_{x,t}$ and $r_{y,t}$ be the rank of $x_t$ and $y_t$ respectively. We dealt with tied points by assigning them the average of the ranks they spanned. For example, if the $x$ series was

$$x_1 = 10, x_2 = 10, x_3 = 5, x_4 = 20$$

then the ranks would be

$$r_{x,1} = 2.5, r_{x,2} = 2.5, r_{x,3} = 1, r_{x,4} = 4.$$

"Normalized" $x$ and $y$ time series are then computed as:

$$\tilde{x}_t = \Phi^{-1}\left(\frac{r_{x,t}}{n+1}\right); \tilde{y}_t = \Phi^{-1}\left(\frac{r_{y,t}}{n+1}\right)$$

for $t = 1, ..., n$, where $\Phi$ is the cumulative distribution function of the standard normal distribution. Next, a recursive procedure is used to look for positive ($S^+$) or negative ($S^-$) local correlations, which can vary over time:

$$S_1^+ = 0; S_1^- = 0$$

$$\begin{aligned} S_{t+1}^+ &= \max(0, S_t^+ + \tilde{x}_t\tilde{y}_t) \\ S_{t+1}^- &= \max(0, S_t^- - \tilde{x}_t\tilde{y}_t). \end{aligned}$$

We then find the maximum of these local (anti)correlations:

$$\begin{aligned} S_{\max}^+ &= \max(S_1^+, S_2^+, ..., S_{n+1}^+)/n \\ S_{\max}^- &= \max(S_1^-, S_2^-, ..., S_{n+1}^-)/n. \end{aligned}$$

Finally, the local similarity score $s$ is given by the maximum correlation or maximum anticorrelation, whichever stronger:

$$s = \begin{cases} S_{max}^+ & \text{if } \mathrm{S}_{\max}^+ > \mathrm{S}_{\max}^- \\ -S_{\max}^- & \text{if } \mathrm{S}_{\max}^+ < \mathrm{S}_{\max}^- \\ 0 & \text{if } S_{\max}^+ = S_{\max}^- \end{cases}$$

#### 7.6 Detailed results

**Gaps filled by linear interpolation**

**A** Number of shared OTUs

| | palm (L) | palm (R) | tongue | gut |
|---|---|---|---|---|
| palm (L) | 240 | | | |
| palm (R) | 234 | 507 | | |
| tongue | 56 | 91 | 180 | |
| gut | 16 | 22 | 4 | 367 |

**B** Pearson correlation-based TTS test *u*-values

| x time series \ y time series | palm (L) | palm (R) | tongue | gut |
|---|---|---|---|---|
| palm (L) | | 0.0125 | 0.0250 | 0.2250 |
| palm (R) | 0.0125 | | 0.0125 | 0.5375 |
| tongue | 0.0250 | 0.0125 | | 0.2375 |
| gut | 0.2000 | 0.5000 | 0.2000 | |

**C** Local similarity-based TTS test *u*-values

| x time series \ y time series | palm (L) | palm (R) | tongue | gut |
|---|---|---|---|---|
| palm (L) | | 0.0125 | 0.0125 | > 1 |
| palm (R) | 0.0125 | | 0.0125 | > 1 |
| tongue | 0.0125 | 0.0125 | | 0.2625 |
| gut | > 1 | > 1 | 0.2875 | |

**Gaps filled by random assignment**

**D** Number of shared OTUs

| | palm (L) | palm (R) | tongue | gut |
|---|---|---|---|---|
| palm (L) | 239 | | | |
| palm (R) | 237 | 517 | | |
| tongue | 57 | 93 | 184 | |
| gut | 16 | 22 | 4 | 388 |

**E** Pearson correlation-based TTS test *u*-values

| x time series \ y time series | palm (L) | palm (R) | tongue | gut |
|---|---|---|---|---|
| palm (L) | | 0.0125 | 0.0125 | 0.3500 |
| palm (R) | 0.0125 | | 0.0125 | 0.1750 |
| tongue | 0.0125 | 0.0125 | | 0.2000 |
| gut | 0.4250 | 0.2125 | 0.1750 | |

**F** Local similarity-based TTS test *u*-values

| x time series \ y time series | palm (L) | palm (R) | tongue | gut |
|---|---|---|---|---|
| palm (L) | | 0.0125 | 0.0125 | > 1 |
| palm (R) | 0.0125 | | 0.0125 | > 1 |
| tongue | 0.0125 | 0.0125 | | 0.2500 |
| gut | > 1 | > 1 | 0.2125 | |

Figure S16: Detailed results from the cross-site microbiome dependence example of Fig 5. We only used measurements from day 42 to day 418 to avoid the longest gaps ($>$ 6 days). Each of the resulting time series contained no more than 41 short (1-day) gaps, no more than 11 medium (2-, 3-, or 4-day) gaps, and no more than 2 longer (5- or 6-day) gaps. Gaps in time series were filled by either linear interpolation (A-C) or random assignment (D-F; each missing taxon abundance value was filled with an abundance value of the same taxon at the same body site from a random time). All random assignments were performed independently (i.e. "with replacement"). (A,D): The number of shared taxa between each body site after removing taxa that were either rare (i.e. absent in half of measurements) or nonstationary (according to an augmented Dickey-Fuller test at the 0.05 significance level). Values along the diagonal denote the total number of taxa at a site after preprocessing. (B,E): TTS test $u$-values from the test based on Pearson correlation. (C,F): TTS test $u$-values from the test based on local similarity scores. All $u$-values shown are raw (i.e. before FDR adjustment). The $y$ time series denotes the series that was used to generate surrogates. Note that tables in B, C, E, and F do not need to be symmetric.

### 8 Detailed methods and results for the zebrafish behavior example

#### 8.1 Obtaining speed and direction time series

We obtained fish trajectory data from the web address at `https://drive.google.com/drive/folders/1UmzlX-yJhzQ5KX5rGry8wZgXvcz6HefD`. We used the file at the path "100/1/trajectories_wo_gaps.npy" (where 100 is for the number of fish per tank, and 1 indicates the first video). This file contains sequences of fish positions indexed by video frame, as well as constants such as frame rate and approximate fish body length. We preprocessed these data using custom scripts inspired by the authors' tutorial at `https://gitlab.com/polavieja_lab/idtrackerai_notebooks/-/blob/master/trajectories_analysis/T1_trajectories_analysis.ipynb`. Where applicable, preprocessing scripts were validated against the authors' trajectorytools package [83].

First, we estimated the position of the center of the tank using the miniball algorithm [119] (as implemented in `https://github.com/weddige/miniball`), and used this to center the data. We then computed fish velocity as

$$\overrightarrow{v}_t = \frac{\overrightarrow{x}_{t+1} - \overrightarrow{x}_{t-1}}{2} R$$

where $\overrightarrow{x}_t$ is a 2-dimensional column vector specifying fish position in units of body length at video frame $t$, $\overrightarrow{v}_t$ is a 2-dimensional column vector specifying the fish velocity in units of body length per second at video frame $t$, and $R$ is the frame rate (32 frames per second). Fish speed was simply defined as $|\overrightarrow{v}_t|$. We then computed direction as the angle $\varphi_t$ between the position and velocity vectors, as described pictorially in Fig 6. Mathematically, $\varphi_t$ is the angle that satisfies:

$$\cos(\varphi_t) = \left(\frac{\overrightarrow{x}_t}{|\overrightarrow{x}_t|}\right)^\top \frac{\overrightarrow{v}_t}{|\overrightarrow{v}_t|}$$
$$\sin(\varphi_t) = \left(\begin{pmatrix} 0 & -1 \\ 1 & 0 \end{pmatrix} \frac{\overrightarrow{x}_t}{|\overrightarrow{x}_t|}\right)^\top \frac{\overrightarrow{v}_t}{|\overrightarrow{v}_t|}.$$

Note that the matrix in the equation for $\sin(\varphi_t)$ is the rotation matrix [120] for a 90° rotation in 2 dimensions. A few fish trajectories had a small number of frames with zero speed. If speed is zero, then direction of movement is, strictly speaking, undefined. To deal with this, we filled in the direction values at zero-speed frames by linear interpolation between the nearest frames with nonzero speeds. Most of the fish trajectories had data gaps, coded as "nan" values in the data files. Since there were still 44 fish trajectories without any gaps, we only performed the TTS test on fish trajectories without gaps.

#### 8.2 Selecting approximately stationary time series

Groups of zebrafish are known to reduce their swimming speed in the days after being introduced into a tank [85]. Indeed, visual inspection of the video from [84] appears to reveal a trend of slowing average fish speed throughout the 10 minutes of tracking in the video, suggesting that fish behavior may be nonstationary (Fig S17A). In fact, this trend is replicable (Fig S17B) across the three replicate videos from [84]. Although the TTS test was robust to nonstationarity arising from a simple linear trend in Fig 2, we still wished to more closely align the data with the theoretical assumption of stationarity. We therefore restricted our analysis to the first $10,000$ frames ($\approx 5$ minutes) of video 1 as there is relatively little systematic trend in the average fish speed in this segment (dotted box in Fig S17A).

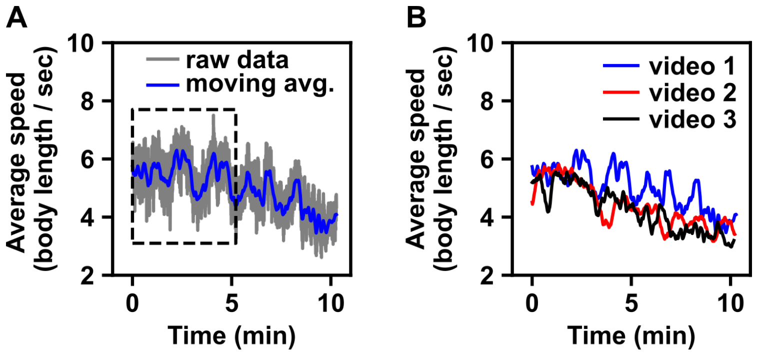

Figure S17: Average fish speeds. (A) Average fish speed in the first video from [84]. The average speed of the 100 fish in the tank is shown in grey. An 11-second moving average of the grey trajectory is shown in blue. The black dotted box shows the first 10,000 frames, which were selected for analysis. (B) 11-second moving averages of all 3 videos from [84].

#### 8.3 Detailed results

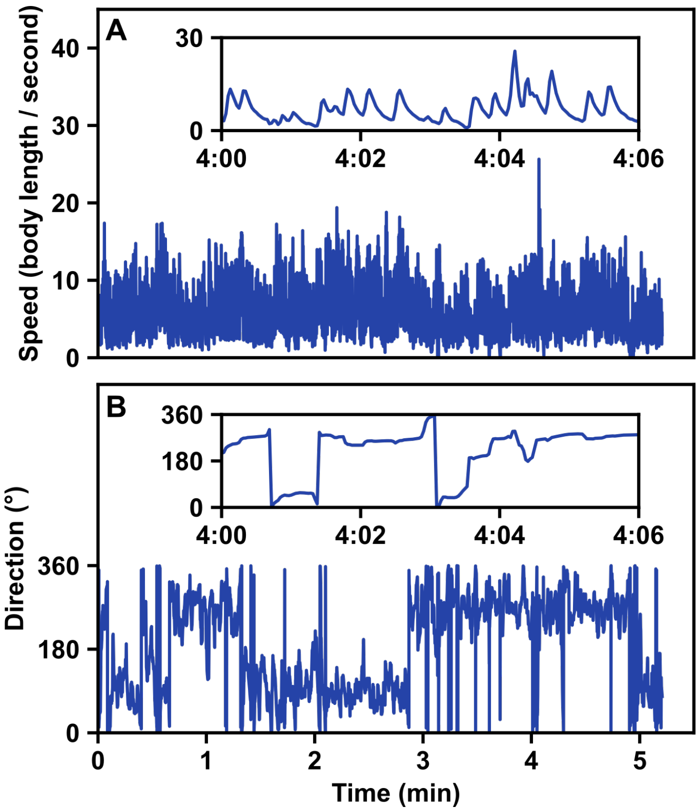

Figure S18: Time series of speed and direction from the fish ("individual 74") analyzed in Fig 6C.

### 9 Difficulty of testing for stationarity

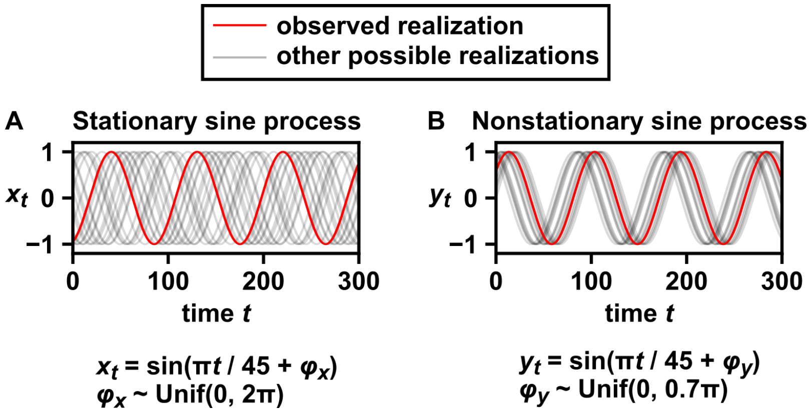

Figure S19: Stationarity is a property of an ensemble of time series, not any single time series. (A) A stationary process that produces a sine wave. Twenty possible realizations of the process are shown. An arbitrary realization is colored red and called the "observed realization". (B) A nonstationary process that produces a sine wave. To see that this process is nonstationary, note that the ensemble mean of processes changes with time. Although only one of the two processes is stationary, a single realization of one process is functionally indistinguishable from a single realization of the other.
